# Functional Convergence of Genetically Diverse B-Cell Receptors in Simian-HIV Infected Rhesus Macaques

**DOI:** 10.64898/2026.01.09.698730

**Authors:** Shengli Song, Hui Li, Chen-Hao Yeh, Akiko Watanabe, Masayuki Kuraoka, Hongmei Gao, Xiaoying Shen, Celia C. LaBranche, Wilton B. Williams, Kevin O. Saunders, M. Anthony Moody, Kevin Wiehe, David C. Montefiori, Barton. F. Haynes, George M. Shaw, Garnett Kelsoe

## Abstract

Germline-targeting or lineage-based vaccine strategies are being used to induce HIV broadly neutralizing antibody (bnAb) responses. These strategies assume that genetically diverse individuals respond similarly to the same immunogen by mobilizing comparable germline precursors, although outcomes vary across bnAb epitopes. Here we explored this premise using a simian-HIV infection model in rhesus macaques. Antigen-unbiased Env-reactive B-cell populations were profiled, followed by systematic analysis of antibody function and B-cell receptor (BCR) genetics. We find that while global Env-reactive B-cell response profiles and antibody functional properties are similar across individuals, underlying BCR genetics are diverse, particularly among non-bnAbs. These results indicate that functional convergence of Env-reactive antibody responses does not necessarily require genetic convergence and suggest that specific germline-targeting may not be an absolute prerequisite for successful vaccine design. These findings support an epitope-focused framework in which bnAb epitopes are engineered to enhance population-level immunogenicity, with potential applicability to other challenging pathogens.

## INTRODUCTION

HIV-1 infection and AIDS threaten not only the health of individuals but also societies and nations. Despite recent breakthroughs in pre-exposure prophylaxis [1, 2], a protective vaccine remains the preferred form of prevention [3, 4] and could be transformative in controlling the HIV-1 pandemic. However, despite decades of research, a safe and effective HIV-1 vaccine remains elusive [5–7], largely due to the virus’s extraordinary genetic diversity [8], complex mechanisms of immune evasion [9–11], and the unique characteristics of the HIV envelope (Env) glycoprotein.

The HIV Env is the sole viral protein on the virion surface that mediates infection and is also the major target of antiviral humoral responses, making it the key immunogen in vaccine design. Env trimers on the surface of infectious virions exhibit considerable structural instability and heterogeneity [12–14]. Nonfunctional gp120/gp41 monomers and gp120-depleted gp41 stumps are readily observed on HIV particles [15]. Furthermore, even the native Env trimer transitions among three distinct prefusion conformations [16]. With few exceptions [17–19], the majority of primary HIV isolates expose high levels of non-neutralizing epitopes [20] that are often immunodominant and distract humoral responses from the frequently sub-dominant neutralizing epitopes [15]. These intrinsic properties of the Env trimer may partially explain why the antibody responses induced by natural HIV-1 infection consist largely of non-neutralizing antibodies (nnAbs) [15, 21].

The failure of early HIV-1 vaccine trials that did not induce broadly neutralizing antibodies (bnAbs), has made the induction of bnAbs the primary goal of HIV vaccine development [22, 23]. The atypical characteristics of many bnAbs, including rare combinations of immunoglobulin (Ig) gene elements, incipient poly-reactivity, and high frequencies of improbable somatic mutations [24–26], suggest that vaccines targeted to specific Ig gene rearrangements (germline-targeting) and sequential (lineage-based) vaccine strategies to guide the somatic evolution of bnAb progenitor cell [27–30] may be necessary to elicit protective HIV-1 bnAb responses. These strategies rely on immunogens designed to recruit rare precursors and recreate similar pathways of bnAb evolution observed in small numbers of human subjects. Many advances have been made in developing germline-targeting immunogens to prime inferred bnAb precursors, including those specific for the CD4-binding site (CD4bs) [31–33], V2 glycan [34–36], V3 glycan [37–39], and membrane-proximal external region (MPER) [40, 41]. Most recently, promising progress has also been made in boosting precursor responses and affinity maturation using more native-like immunogens in human Ig gene knock-in mouse models [42–44].

Despite these advances, remarkable challenges remain in the pursuit of a generalizable vaccine capable of consistently inducing bnAb responses across genetically diverse populations. First, the efficacy of initial germline priming is highly dependent on precursor frequencies [45, 46] and their affinities for the designed immunogen [46, 47]. Second, population-wide allelic diversity among Ig loci further complicates germline selection and immunogen design [48]. Third, even with successful recruitment of germline bnAb progenitors, their shepherding along specific evolutionary trajectories to ensure entry of mature bnAb B cells into the long-lived plasma cell compartments remains a major hurdle [7].

A fundamental, yet less-addressed question of HIV-1 vaccine design concerns the similarity or dissimilarity of humoral responses to identical immunogens among genetically diverse individuals. The hypothesis underlying germline-targeting or lineage-based vaccine strategies posits that engineered priming immunogens can engage defined bnAb germline precursors, when present in the naïve repertoire, and guide their somatic evolution along related pathways toward bnAb responses. More succinctly, can the same series of priming, boosting and polishing immunogens elicit similar humoral responses in different individuals? If humoral responses to identical immunogens are largely similar across diverse individuals, these responses can be considered deterministic, supporting the general applicability of vaccine strategies aimed at recapitulating rare or disfavored B-cell lineages to elicit bnAb production. Conversely, if humoral responses to identical immunogens vary substantially between individuals, such variability must be explicitly considered in vaccine design to ensure practical efficacy.

Our study addresses this fundamental question using a well-established simian-HIV (SHIV) infection model in rhesus macaques (RMs) [49, 50]. The SHIV strains used in this study bear Env glycoproteins derived from primary transmitted/founder (T/F) HIV strains and follow conserved mutation patterns in infected RMs that mimic patterns observed in the corresponding humans from which the Envs originally derived [51, 52], generating dynamic quasispecies [53] of Env antigens in real time. A high-throughput single B-cell culture method was established to profile B-cell populations from these infected RMs in an unbiased manner, enabling systematic analysis of Env-reactive B-cell responses from both the functional and genetic perspectives. Our results demonstrate that while the global B-cell response profiles and the functional properties of reactive antibodies are similar across RMs, the underlying B-cell receptor (BCR) genetics are diverse. These findings indicate that functional convergence can arise despite substantial diversity in underlying BCR genetics, motivating an epitope-focused framework in which shared epitope-level constraints, rather than specific genetic solutions, shape convergent antibody responses across individuals.

## RESULTS

### Similar viral load kinetics and shared Env gene mutation patterns in RMs infected with molecularly cloned SHIVs

Two cohorts of six outbred, Indian-origin RMs were infected with molecularly cloned strains of SHIVs that express the Env glycoproteins of T/F HIV-1 virus strains, BG505.T332N (BG505 for short, Clade A) or CH505 (Clade C), respectively [49]. The original BG505 HIV infection elicited bnAb responses of undetermined specificity in a person living with HIV (PLWH) [54, 55], and two different types of CD4bs bnAb lineages were identified from the human subject infected with CH505 HIV [56, 57]. In the CH505 PLWH, both the HCDR3-binding CH103 CD4bs bnAb lineage and the CD4-mimicking CH235 CD4bs bnAb lineage were isolated [30, 56]. Two different infection routes, intrarectal and intravenous, were used for the two cohorts respectively, representing the two major routes of HIV transmission in humans.

Infections were monitored by determining circulating viral loads in plasma samples taken at different intervals throughout the infection course. All 12 RMs showed similar early kinetics of plasma viral loads (Figure 1A), making the infection responses comparable. Consistent with previous studies with other viral strains [49, 58], the early acute peaks followed by decline and persistence in plasma viral loads indicates successful chronic SHIV infection in these RMs.

**Figure 1.**
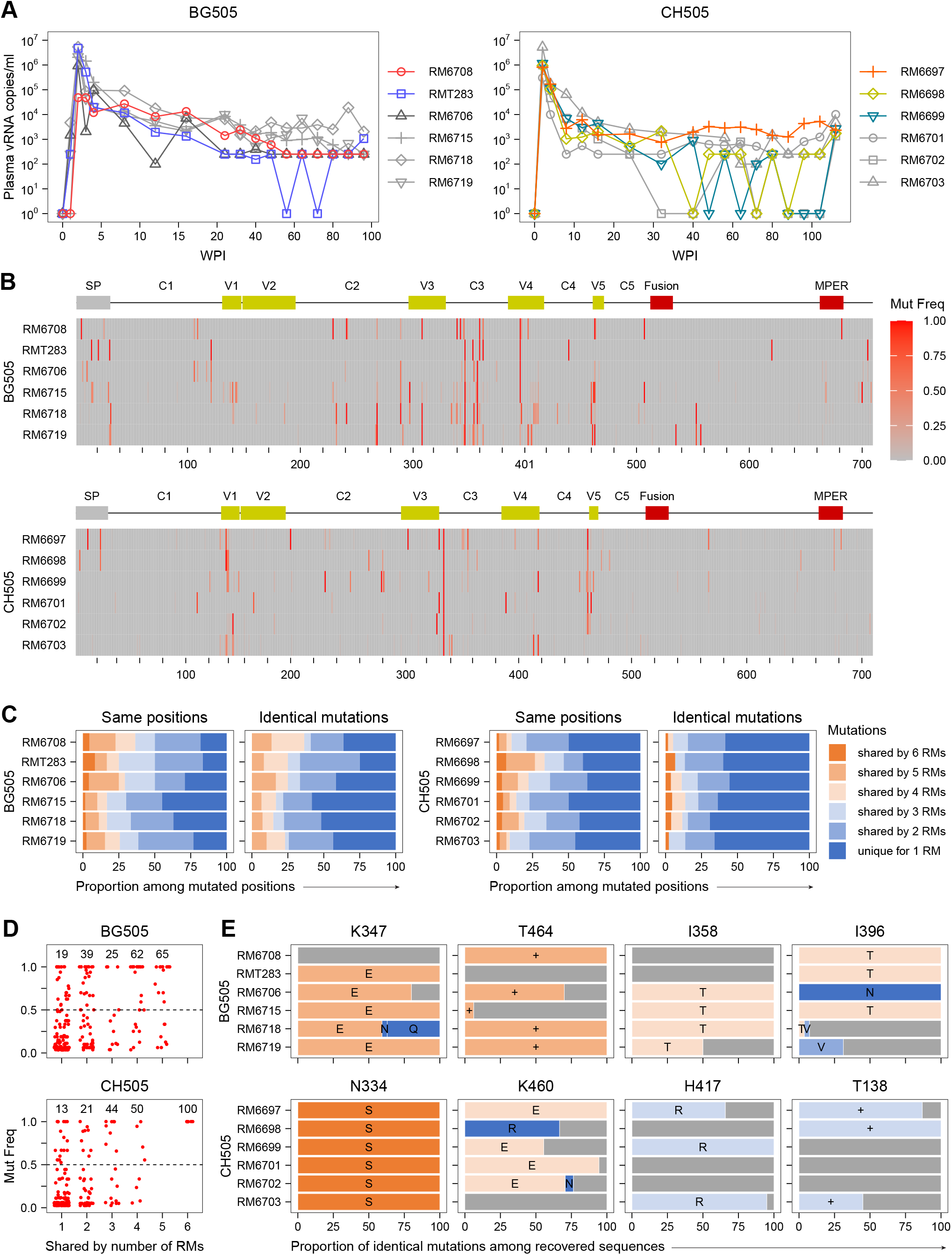
Similar viral load kinetics and shared Env gene mutation patterns in RMs infected with molecularly cloned SHIVs. (A) Plasma viral load kinetics measured by qPCR of viral RNA copies. Values below detection limits were set to 1 for visualization. (B) Mutation frequency at individual amino acid residues based on Env sequences recovered from each RM. Domain (top) and position (bottom) annotations follow HXB2 numbering. (C) Proportion of mutations unique to a single RM or shared by different numbers of RMs per group. Mutations at the same positions or identical mutations are counted separately. (D) Mutation frequencies of shared mutations. Each dot represents a mutation from one RM. Numbers above columns indicate the percentage of mutations with frequencies >0.5 (dashed line). (E) Examples of identical mutations shared among RMs. Titles indicate residues in the infecting T/F virus strain. Letters on individual bar segments indicate the one-letter codes of the mutated amino acid residues. +, insertion; –, deletion. Color-coded as in C, with grey denoting unmutated proportions.

We compared mutated amino acid sequences across all six RMs in each infection group. Consistent with previous findings [51], Env gene mutation patterns were shared among RMs infected with the same SHIV (Figure 1B; Documents S1 and S2). Mutation hotspots were identified in C3, V4, and V5 domains for BG505-infected RMs and in V1, C3, and C4 for CH505-infected RMs. Overall, Env mutation patterns were conserved within each SHIV infection group but differed between groups, suggesting that viral factors shape these patterns despite host genetic differences.

Similar to the observation in humans [57], mutations at the same positions, including identical amino acid changes, were observed among RMs. In the BG505 infection group, 55.0% – 83.3% of mutations occurred at the same positions in at least two RMs, with 41.7% – 75.0% showing identical mutations. In the CH505 infection group, 50.0% – 65.5% of mutations were at the same positions, and 35.7% – 44.8% were identical (Figure 1C). Additionally, a substantial portion of these shared identical mutations occurred at high frequencies (Figure 1D) among recovered viral genome sequences, as exemplified in Figure 1E. This chronic SHIV infection model provides dynamic quasispecies of Env antigens in a real-time fashion and serves as a straightforward system to compare humoral responses across individual RMs.

### Persistent and productive GC responses in RMs with chronic SHIV infection

To elucidate the dynamics of B-cell responses to SHIV infection, we focused our analysis on three RMs that received similar infection regimens and exhibited similar viral load kinetics (top three RMs in Figure 1A) for each infection group. Lymph node (LN) biopsies were taken from selected RMs at various timepoints after infection. Phenotyping of the isolated LN cells identified four major B-cell populations: IgG^+^ germinal center (GC) B cells, IgG^+^ memory B (Bmem) cells, IgM^+^ Bmem cells, and mature follicular (MF) B cells (Figure 2A). All four B-cell populations, and importantly the IgG^+^ GC B-cell population, persisted in all RMs throughout the sampling time course (24-to 56-or 72-weeks post-infection), although the magnitudes and kinetics of each population varied remarkably among different RMs (Figure 2B and Table S1).

**Figure 2.**
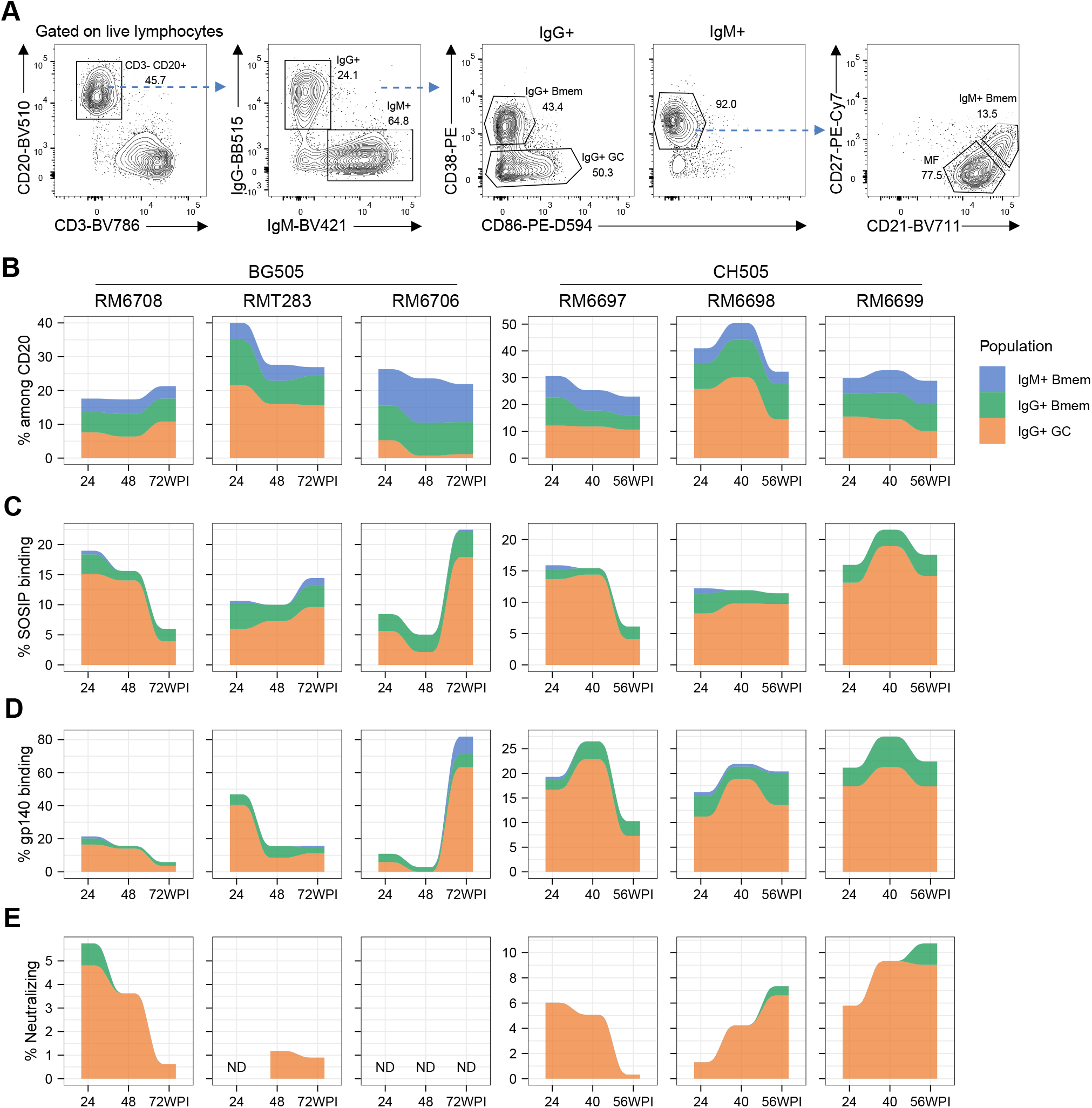
Persistent and productive GC responses in RMs with chronic SHIV infection. (A) Flow cytometry gating strategies to identify B-cell populations in LN cells derived from SHIV-infected RMs. (B) Alluvial plots showing percentages of IgG^+^ GC, IgG^+^ Bmem and IgM^+^ Bmem populations among CD20^+^ LN cells during the infection time course in the three RMs in each infection group. (C-E) Alluvial plots showing percentages of SOSIP-binding (C), gp140-binding (D), and autologous neutralizing (E) IgG^+^ single B-cell cultures from different B-cell populations over the infection course. See also Figures S1 and S2, and Tables S1 and S2.

The MS40L-low feeder line [59] was previously used to establish a robust human single B-cell culture method in our laboratory [60, 61]. To enable high-throughput functional and genetic profiling of GC and Bmem populations in SHIV-infected RMs, we adapted this method for RM B-cell cultures (Figure S1A). We further engineered the MS40L-low line to autonomously express cytokines necessary for supporting B-cell proliferation and differentiation. The resulting cloned feeder cell line, MEC-147, supports single B-cell cultures from both RMs (Figure S1B) and humans (Figure S1C), achieving cloning efficiencies and antibody production comparable to the parental MS40L-low subline plus exogenous cytokines. This streamlined culture process proved more amenable to high-throughput screening than the original MS40L-low method [60, 61].

IgG^+^ GC B, IgG^+^ Bmem, IgM^+^ Bmem, and MF B cells were recovered from lymph nodes of infected RMs by flow cytometry and individually sorted into single B-cell cultures without selection for antigen specificity [61]. From six RMs, 67,536 single B-cell cultures were screened, identifying 17,259 IgG^+^ clonal cultures by ELISA (25.6% cloning efficiency; Tables S1 and S2). Antibodies produced and clonal B cells expanded from these single B-cell cultures provide us with the functional and genetic materials to profile B-cell responses in SHIV-infected RMs. This antigen-unbiased approach profiles the B-cell repertoire by phenotype rather than specificity, allowing population-level analyses of Env-reactive B cells. Notably, given the low surface Ig expression of GC B cells [62], this strategy is essential for capturing their role in chronic humoral responses.

Culture supernatants containing clonal IgGs were tested for binding activity in Luminex assays against an antigen panel that included autologous SOSIP and gp140, CH505 gp120 and MN gp41 domain antigens, an anti-rhesus IgG capture antibody, and non-relevant control antigens such as streptavidin, bovine serum albumin (BSA), and ovalbumin (Table S2). The majority of autologous SOSIP (Figure 2C) and gp140 (Figure 2D) binding cultures originated from IgG^+^ GC B cells, with responses varying among RMs (Figure S2). Of 2,049 Env-binding cultures identified, 1,566 (76.4%) were derived from IgG^+^ GC B cells, 311 (15.2%) from IgG^+^ Bmem, and 161 (7.9%) from IgM^+^ Bmem (Table S1).

In parallel and in a blinded manner, culture supernatants containing clonal IgGs were subjected to standard TZM-bl neutralization assays against autologous SHIVs [63, 64]. Consistent with Env-binding results, the great majority of autologous neutralizing cultures were from the products of IgG^+^ GC B cells (Figures 2E and S1D). In total, 319 autologous neutralizing cultures were identified, among which 308 (96.6%) were derived from IgG^+^ GC B cells, and 11 (3.4%) from IgG^+^ Bmem (Table S1). Similar to the Env-binding results, the magnitude and kinetics of autologous neutralizing GC B-cell responses varied among different RMs. All neutralizing cultures also showed significant binding to Env antigens in Luminex assays. The remaining 1,258 (80.3%) IgG^+^ GC, 300 (96.5%) IgG^+^ Bmem, and 161 (100%) IgM^+^ Bmem cultures demonstrated binding to Env antigens but no detectable autologous neutralizing activity, suggesting that the B-cell responses were predominantly non-neutralizing in these SHIV-infected RMs. Collectively, persistent and productive GC responses were induced by chronic SHIV infection in all infected RMs.

### Distinct clonal dynamics of neutralizing and non-neutralizing Env-reactive IgG+ GC B-cell clones

To study the evolution of Env-reactive B-cell responses during chronic SHIV infection, we determined the V(D)J sequences of neutralizing and/or Env-binding BCRs identified from single-cell cultures and screening. Given the high diversity and incomplete characterization of RM IGV repertoires [65, 66], we developed a versatile and unbiased BCR V(D)J sequencing platform (see Methods). Using this platform, we determined the V(D)J sequences of 1,499 neutralizing and/or Env-binding B cells, covering 95.3% – 100% of identified autologous SOSIP-binding and 91.3% – 99.3% gp140-binding single-B-cell cultures with median fluorescence intensity (MFI) greater than 300 across the six RMs (Figure S3A). V(D)J assignment and clonal analyses were performed based on the sequence data (see Methods). These genetic data were combined with time-course, phenotypic, and functional data (Table S3) and visualized as phylogenetic trees for 159 B-cell lineages (Document S3). Based on these combined data, a systematic analysis was then carried out as detailed below to compare the similarities among individual RMs infected with the same viruses.

We first focused on IgG^+^ GC B cells, as most neutralizing and Env-binding clones originated from this population (Figure 2). B-cell singletons or lineages containing at least one member showing >50% autologous neutralization activities in the initial screening were classified as neutralizing, while all others were defined as non-neutralizing. Clonal expansion/contraction of IgG^+^ GC B cells was visualized (Figure 3A), revealing varied expansion kinetics among RMs and individual clones. However, distinct clonal dynamics between neutralizing and non-neutralizing clones were consistently observed across RMs. First, neutralizing lineages persisted across multiple timepoints more frequently than non-neutralizing ones (Figures 3A and 3B), despite sampling from anatomically separated LNs at different timepoints. This observation implies that Bmem or GC B cells circulate and re-enter GC responses at distal LNs during chronic SHIV infection. Second, neutralizing clones showed more extensive clonal expansion (Figure 3C) but lower repertoire diversity (Figure 3D) compared to non-neutralizing IgG^+^ GC B cells, consistent with the well-established dominance of non-neutralizing Env-reactive responses and the restriction of neutralizing activity to a smaller subset of B-cell clones during HIV infection [24].

**Figure 3.**
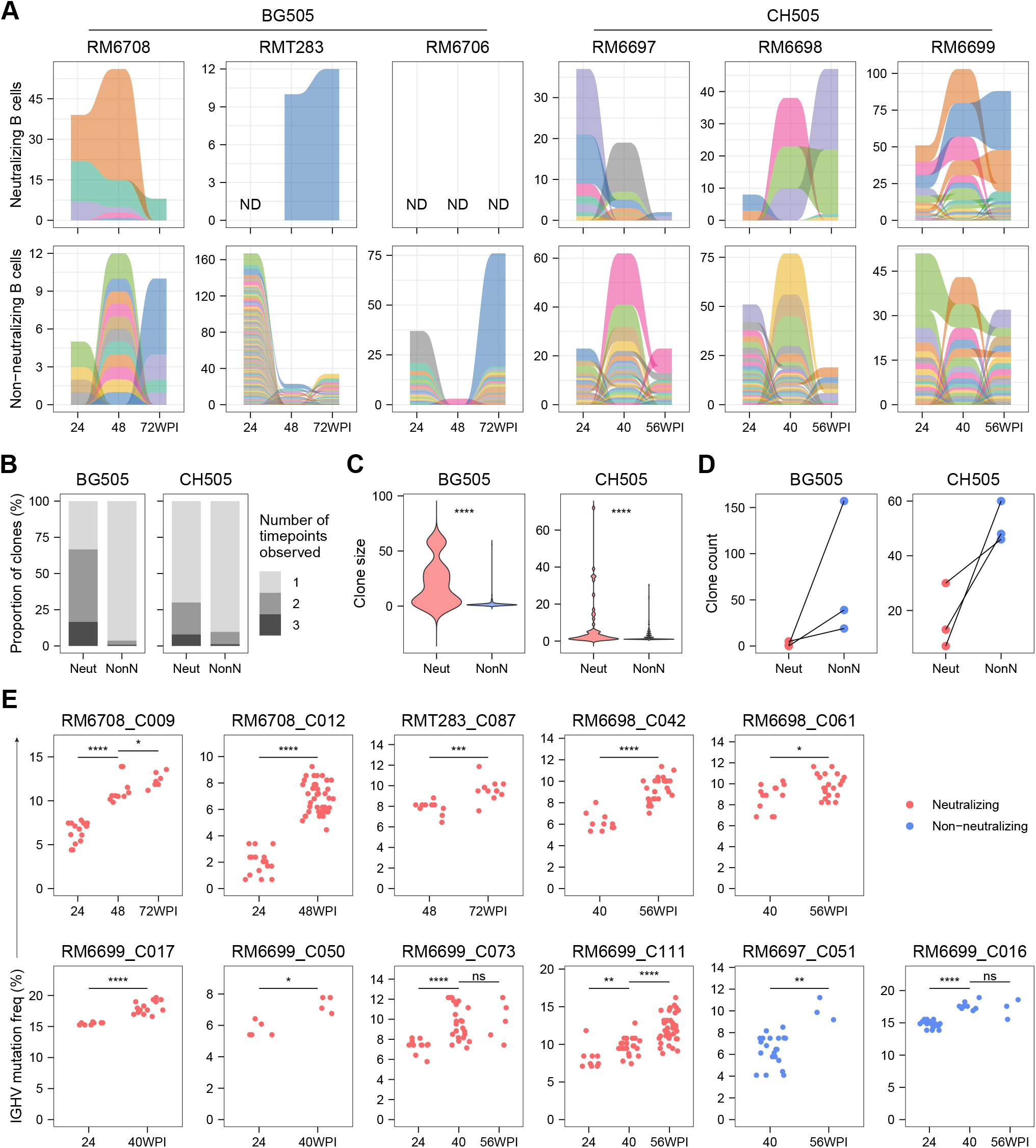
Distinct clonal dynamics of Env-reactive neutralizing and non-neutralizing IgG^+^ GC B-cell clones. (A) Expansion/contraction of Env-reactive neutralizing and non-neutralizing IgG^+^ GC B-cell clones in each infection group over time. ND, not detected. Individual clones are displayed in different colors to facilitate visual distinction between adjacent flows. The colors are assigned arbitrarily and do not represent specific clone identities or biological characteristics. (B-C) Distributions of timepoint occurrence (B) and clone size (C) were compared between neutralizing (Neut) and non-neutralizing (NonN) clones. Two-sided Wilcoxon rank-sum tests at the clone level (pooled across RMs within each SHIV group) were used. (D) Clone counts of neutralizing and non-neutralizing IgG^+^ GC B-cell clones in individual RMs. (E) IGHV mutation frequency changes in Env-reactive clones spanning multiple timepoints. All nine clones (named “RM_Clone-ID”) that persisted across at least two timepoints and contained more than two clonal members at each timepoint are shown. Adjacent timepoints were compared using two-sided Wilcoxon rank-sum tests. Statistical annotations: *, p < 0.05; **, p < 0.01; ***, p < 0.001; ****, p < 0.0001; ns, not significant. See also Table S3 and Document S3.

The IGHV mutation frequencies were compared for those B-cell lineages recovered over multiple timepoints. As shown in Figure 3E, in all cases for nine neutralizing and two non-neutralizing lineages polled from five RMs analyzed, significant accumulation of somatic hypermutations overtime were detected in the IGHV genes. This observation is consistent with the possibility that Bmem or GC B cells circulate and re-enter GCs in distal sites during chronic and systemic SHIV infection.

### Limited clonal relationship between Env-reactive Bmem and IgG^+^ GC B cells

In parallel, we analyzed Env-reactive IgG^+^ and IgM^+^ Bmem cells also recovered in the LN biopsies from infected RMs (Table S3). Compared to the local IgG^+^ GC B cell populations, Bmem populations contained a higher proportion of non-neutralizing B cells across all RMs in general (Figure 4A). This might be due to preferential selection of GC B cells with lower neutralization activity/affinity into the Bmem pool or the consequence of preferential Bmem differentiation from a distinct population of specific B cells comprising a majority of non-neutralizing Env-reactive cells.

**Figure 4.**
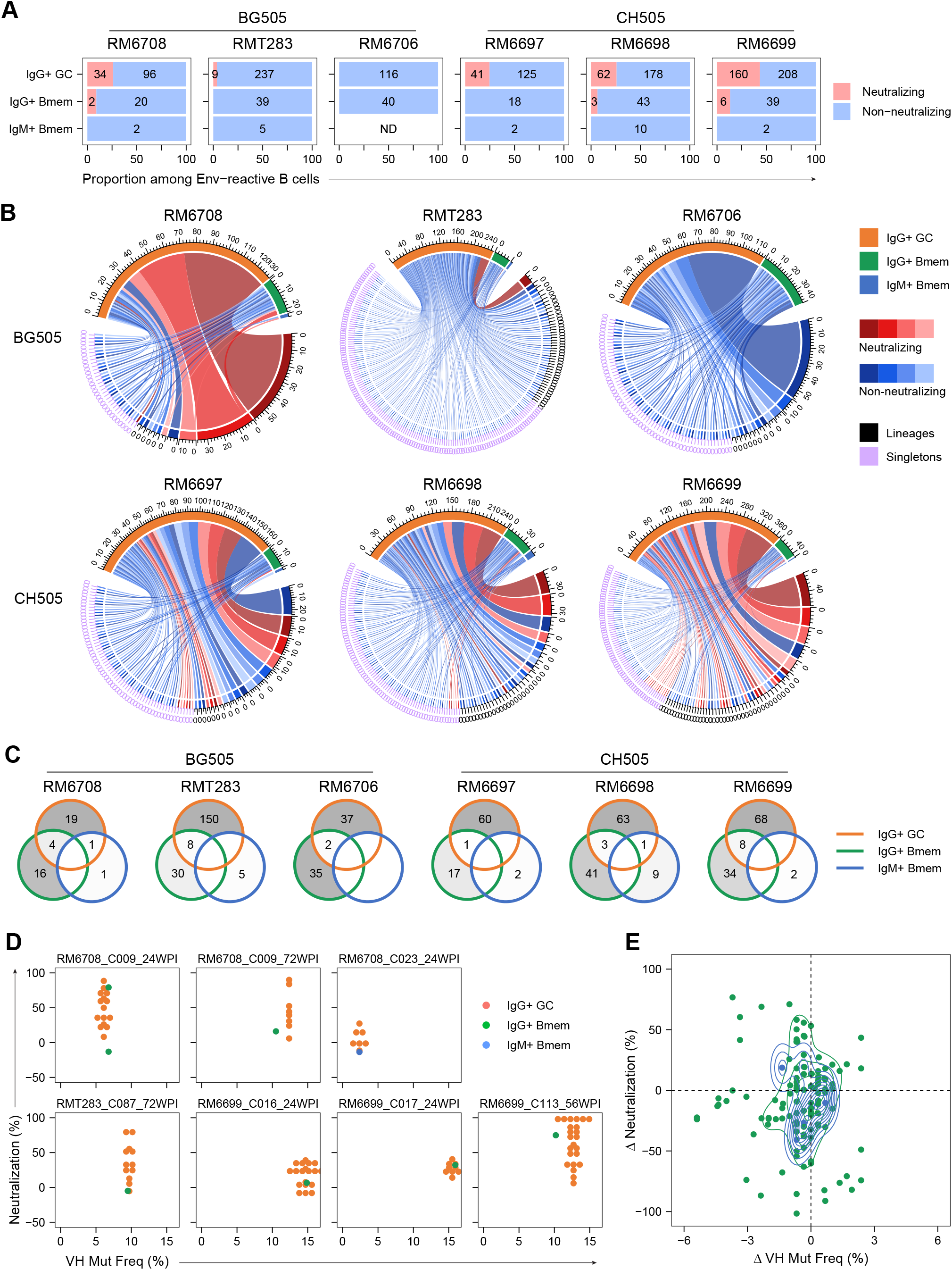
Limited clonal relationship between Env-reactive Bmem and IgG^+^ GC B cells. (A) Proportions of neutralizing and non-neutralizing Env-reactive B cells from different populations. Inserted numbers indicate cell counts. ND, not detected. (B) Clonal relationship between Env-reactive IgG^+^ GC and Bmem cells. Top sectors show B-cell populations (color-coded); lower sectors represent individual neutralizing or non-neutralizing clones, including lineages and singletons. Color gradients (red for neutralizing and blue for non-neutralizing) were used solely to visually distinguish adjacent individual clones. (C) Venn diagrams showing the distribution of B-cell clones across single or multiple populations. Blank regions indicate zero counts. (D-E) For lineages containing both IgG^+^ GC and Bmem members at the same timepoints: (D) V_H_ mutation frequencies and neutralization activities for all seven RM_Clone-ID_Timepoint groups identified with >3 clonal members; (E) V_H_ mutation frequency and neutralization differences between Bmem and IgG^+^ GC B cells for all 19 RM_Clone-ID_Timepoint groups identified. Each dot represents a Bmem–IgG+ GC B cell pair. See also Table S3.

To differentiate between these possibilities, we determined the clonal relationships between Env-reactive IgG^+^ GC and Bmem cells recovered from LN samples (Figure 4B). In all RMs, most (70.2% – 95.0%) Env-reactive Bmem cells were singletons (Figure 4B); and only a minority of Bmem clones (13.1% for IgG^+^ Bmem clones and 9.5% for IgM^+^ Bmem clones) were clonally related to IgG^+^ GC B cells recovered from the same animal (Figure 4C). These infrequent clonal associations support the possibility that the Bmem pools in infected RMs are preferentially derived from activated but non-neutralizing B-cell pools. Given evidence (above) that circulating Bmem cells can re-enter GCs in distal LN sites (Figures 3A and 3B), a third, non-exclusive possibility is that neutralizing, but not non-neutralizing Bmem cells are preferentially selected for GC re-entry.

One limitation in the analysis above originated from the criterion we used to define a B-cell clone as neutralizing or non-neutralizing, which is based on the highest neutralization activity achieved among all member(s) of that B-cell clone. Therefore, B-cell singletons and lineages of small sizes may be undervalued in this binary categorization due to limited sampling sizes. To address this limitation, we pulled out all lineages that consist of members from both GC and Bmem populations at the same timepoints and compared the V_H_ mutation frequencies and neutralization activities of clonal members from different populations (Figure 4D). As shown for individual RMs in Figure S3B and aggregated in Figure 4E, there aren’t consistent differences between Bmem and IgG^+^ GC populations in V_H_ mutation frequencies or neutralization activities, arguing against preferential selection of B cells of lower neutralization activities into Bmem pool from lineages with neutralizing IgG^+^ GC B cells.

### Distinct functional types of Env-reactive B-cell clones defined by antigen binding and neutralization

Considering the intrinsic instability and conformational dynamics of the Env trimers, we reasoned that a comparison of binding activities to the stabilized SOSIP versus Env in “open” conformations would permit assessment of the quality of Env-reactive B-cell responses; this comparison also provides a measure of the quality of humoral responses between individual infected RMs. Unstabilized gp140 proteins, which lack the transmembrane and intracellular domains required for stabilizing the Env spike on cell membranes and virions [20, 67, 68], were therefore selected as antigens representing open conformations.

We tested this measure of antibody quality using reference bnAbs and nnAbs, with the latter binding predominantly to open-conformation epitopes (Figure S4A). As the CD4bs-specific bnAb VRC01 [69] binds SOSIP and gp140 proteins similarly [70], we normalized MFI binding ratios of all tested antibodies to that of VRC01. Relative binding activities correlated with epitope specificities for both BG505 and CH505 Env antigens (Figure 5A). Specifically, CD4bs and V3 glycan-dependent bnAbs behaved similarly to VRC01 by comparable binding to both SOSIP and gp140, whereas V1/V2-specific, and gp120/gp41 interface antibodies preferentially bound SOSIP over gp140. In contrast, V3 domain open-conformation, CD4-induced (CD4i) epitope, and CD4bs-specific tier-1-neutralizing antibodies exhibited a four-to 1200-fold higher binding for gp140 relative to SOSIP. These findings suggest that differential binding to SOSIP and gp140 serves as a reliable indicator to distinguish specificity for closed versus open conformations on Env trimers [71].

**Figure 5.**
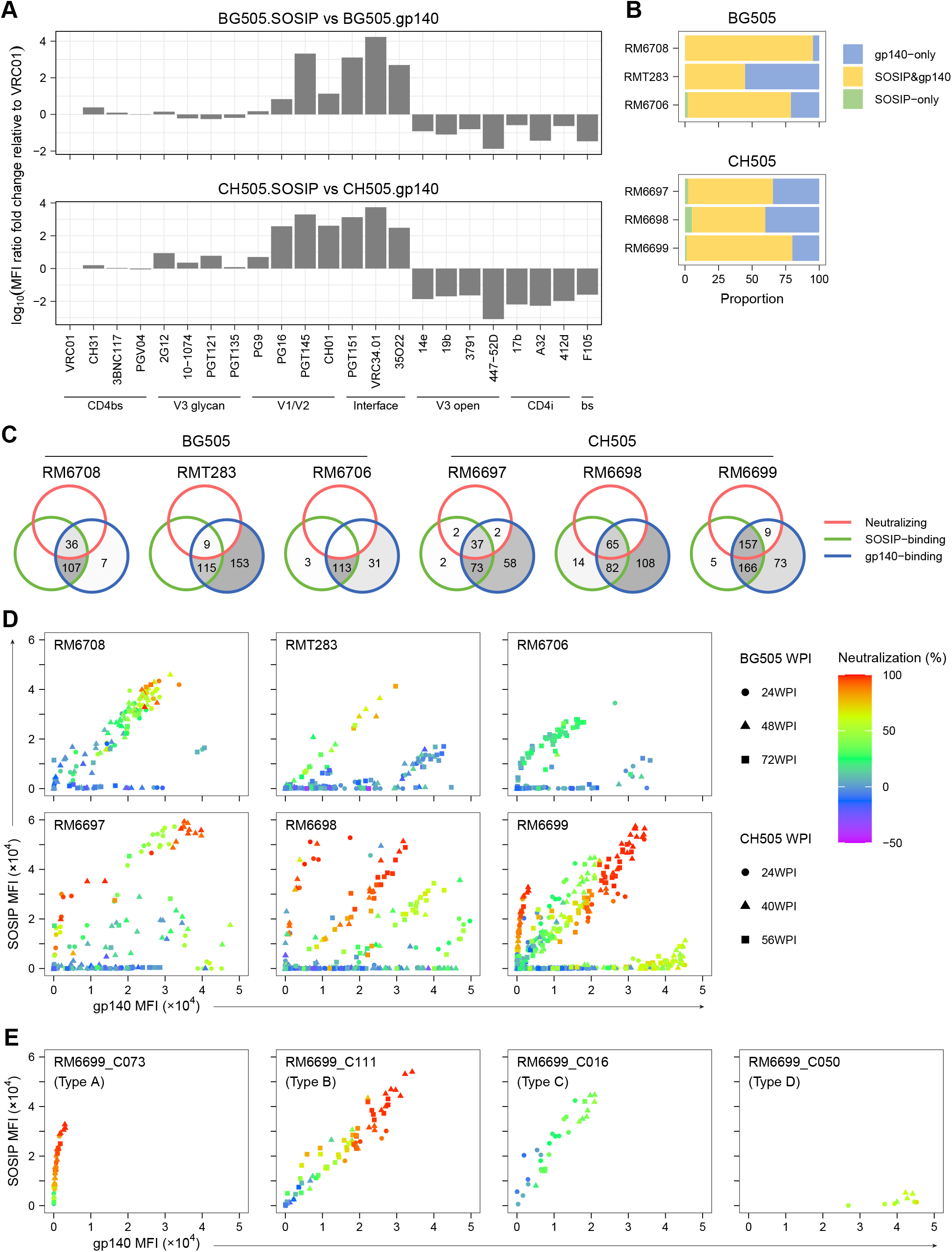
Distinct functional types of Env-reactive B-cell clones defined by antigen binding and neutralization. (A) Log_10_-transformed fold-change of MFI ratios (SOSIP/gp140) for selected bnAbs and nnAbs, normalized to VRC01. MFI values at a same concentration (206 ng/ml, dashed lines in Figure S4A) were used; similar trends observed at other concentrations. bs, CD4bs-specific tier-1-neutralizing. (B) Composition of antigen specificities of Env-reactive antibodies identified from SHIV-infected RMs. (C) Venn diagrams categorizing Env-reactive antibodies by autologous neutralization and SOSIP/gp140 binding. Blank regions indicate zero counts. (D) Dot plots of SOSIP-vs. gp140-binding MFI values for Env-reactive antibodies in individual RMs, with a color gradient for autologous neutralization. (E) Similar plotting as (D), exemplifying four major types of Env-reactive B-cell clones (“RM_Clone-ID”). See also Figure S4, Table S3, and Document S4.

Using this method, we compared SOSIP/gp140 antigen reactivities of all Env-reactive clonal antibodies recovered from SHIV-infected RMs by single B-cell culture (Table S3). As shown in Figure 5B, while the majority of antibodies bind to both autologous SOSIP and gp140 antigens, substantial fractions (4.7% – 55.2%, average 29.3%) of antibodies bind only to gp140 antigens, suggesting that the immune recognition of Env antigen in open conformation is common in SHIV-infected RMs. Incorporating neutralization data, we found that the vast majority (95.9%) of autologous neutralizing antibodies bound both SOSIP and gp140 (Figure 5C). A small subset (3.5%) recognized only gp140 but exhibited weaker neutralization (51.2–67.2%, Table S3). These findings align with vaccination studies [72] showing that stabilized prefusion Env trimers are optimal for eliciting neutralizing B-cell responses to difficult-to-neutralize HIV isolates.

We compared the functional similarities of Env-reactive antibodies among SHIV-infected RMs based on autologous neutralization and SOSIP vs. gp140 binding activities. The antibodies formed clusters based on relative binding activities (Figure 5D), with autologous neutralization correlating with binding strength (increasing MFI) within each cluster. Although these clusters form unique binding/neutralization patterns for each RM analyzed, similar clusters can be identified from different RMs, especially within the same SHIV infection group. This observation suggests the functional similarity in the scope of antigen specificities but different composition of Env-reactive B-cell repertoire in these RMs infected with identical SHIVs.

We further incorporated BCR genetics into the functional analysis of neutralization/antigen binding activities. Individual B-cell lineages identified were pulled out and plotted in the same way as in Figure 5D. As expected, the binding activities for SOSIP and gp140 antigens correlated with each other in general for clonal members of the same lineages (Document S4). Importantly, the distribution of these lineages corresponds to the distinct clusters observed in the plots above with total Env-reactive antibodies (Figure 5D). There are four major types (see Methods) of Env-reactive B-cell clones based on this plotting (Figure 5E). Type A clones bind preferentially to SOSIP antigen and the majority show strong neutralization activities; Types B and C clones bind to both SOSIP and gp140 vigorously, with Type Bs showing strong (≥ 50%) and Type Cs showing weak (< 50%) autologous neutralization activities; Type D clones bind preferentially to gp140 protein and most exhibit weak neutralization activities. Considering the conformational differences between SOSIP and gp140 antigens, these different functional types presumably represent specificities for different types of epitopes exposed differentially on SOSIP and gp140 antigens. These functional types can then be used as surrogate indications to compare the response patterns in different RMs.

### Shared functional characteristics of Env-reactive recombinant antibodies derived from different RMs

To characterize further Env-reactive B cells across SHIV-infected RMs and compare their functional similarities, we produced 146 recombinant antibodies (rAbs, Table S3) from representative clones based on the four functional types (A – D) shown in Figure 5E and confirmed their Env-binding activities (Figure S4B and Document S5). The relative binding activities of these rAbs for autologous SOSIP and gp140 antigens matched that observed for culture supernatants (Figure S4C). We also detected the binding activities towards heterologous SOSIP and gp140 antigens, with rAbs derived from BG505 (clade A) infected RMs for CH505 (clade C) Env antigens, and vice versa. Notably, cross-clade binding was observed in 98.7% (75/76) of Type D and 38.1% (8/21) of Type C rAbs, but only 13.3% (2/15) of Type A and none (0/34) of Type B (Figure 6A). All cross-clade Type D and Type C rAbs were specific for the gp120 subunit, as determined from binding data with the corresponding culture supernatants during initial screening (Figure S3C). This observation suggests that non-neutralizing epitopes recognized by Types C and D antibodies are relatively conserved among HIV-1 viruses of different clades and, likely, among progeny viruses within the infected individual.

**Figure 6.**
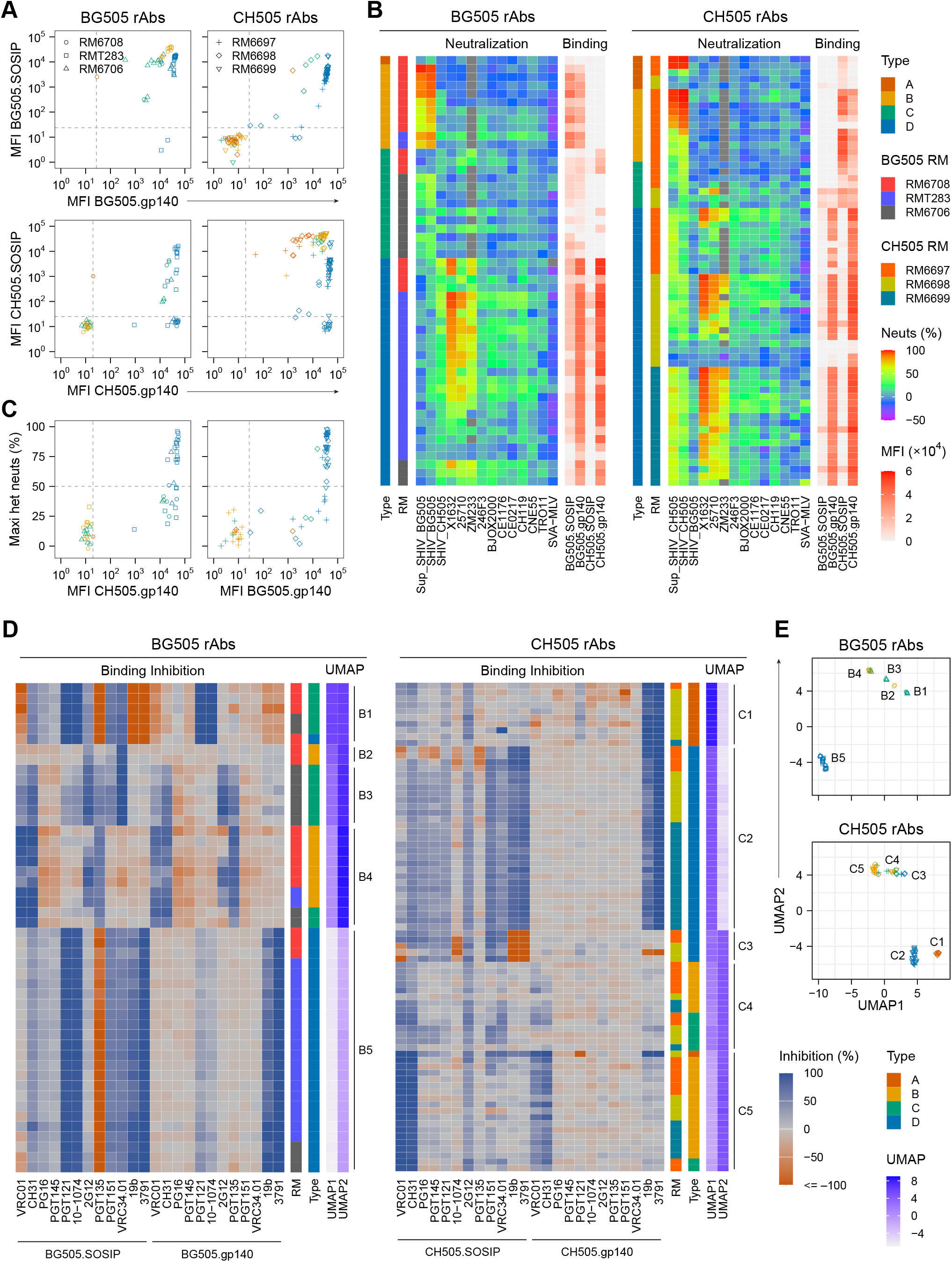
Shared functional characteristics of Env-reactive recombinant antibodies derived from different RMs. (A) Env-binding MFI values of rAbs to autologous and heterologous SOSIP/gp140. Functional types of source B-cell clones are color-coded (legend in B). Dashed lines indicate binding cutoffs. (B) Heatmaps of rAb (rows) neutralization against autologous and heterologous pseudo-viruses. Autologous SHIV neutralization using culture supernatants (Sup_SHIV) and SOSIP/gp140 binding (panel A) are included for comparison. (C) Maximum heterologous neutralization vs. gp140 binding of rAbs (panel B), with dashed cutoffs for neutralization and binding. (D-E) Heatmaps (D) showing percentages of inhibition of analyte rAbs (rows) binding to autologous SOSIP or gp140 antigens in the presence of bnAbs or nnAbs competitors. UMAP coordinates are displayed in parallel within (D) and as 2D plots in (E), labeled with clusters identified using the DBSCAN algorithm. RMs and Types in panel D use the same color-coding as in B. Color-coding in panel E shows types as in B, and shape-coding shows RMs as in A. See also Table S3, S4, and S5, Figures S3, and S4, and Document S5.

Neutralization breadth of rAbs was assessed using TZM-bl assays [63, 64] against 11 heterologous tier 2 pseudo-viruses [73] and the autologous SHIV strain (Table S4). Autologous neutralization results were consistent with initial culture supernatant screenings (Figure 6B). Surprisingly, heterologous neutralization was most often mediated by rAbs from Type D and, to a lesser extent, Type C clusters (Figure 6C). Notably, the highest heterologous neutralization activities were detected for X1632 (Clade G), 25710 (Clade C), and ZM233 (Clade C), regardless of the originating SHIV strain (Figure 6B). Interestingly, the same subset of antibodies exerted heterologous neutralization activities for these three viruses (Figure 6B). And these heterologous neutralizing antibodies overlap with the subset exhibiting binding activities for heterologous gp140 antigens (Figure 6B, 6C). In contrast, whereas rAbs from Type A or B clusters were enriched for autologous neutralization, neutralization breadth was largely absent. This functional pattern holds true across different RMs, regardless of the infecting SHIV clade, indicating a consistent functional profile in the antibody responses to SHIV infections.

To explore further functional similarities of SHIV elicited humoral responses, we determined epitope specificity patterns of selected rhesus rAbs through competition binding assays. Binding of these rAbs (analytes) to autologous SOSIP and gp140 (antigens) were measured in the presence of various human bnAbs and nnAbs (competitors) with known specificities (Table S5). We plotted the percentage of inhibition of analyte-antigen binding by these competitors (Figure 6D). Subsequently, we applied Uniform Manifold Approximation and Projection (UMAP) [74] to the binding inhibition data, which identified clusters of analyte antibodies (Figure 6E). The UMAP clusters correlate with specific binding inhibition patterns and align generally with previously defined functional types. Except for cluster C4, all other clusters consist predominantly or exclusively of rAbs from one functional type (Figure 6D). This clustering analysis not only supports the previous functional type categorization but also constitutes an additional dimension to the assessment of functional similarities across RMs.

Based on the specificity clustering, we compared the similarities across RMs. The majority of the specificity clusters (8 out of 10) are comprised by rAbs from multiple RMs (Figure 6D), consistent with the conserved specificity profile among Env-reactive BCRs/Abs derived from different RMs infected with the same virus. In contrast, UMAP comparisons of rAbs from BG505 and CH505 infection groups showed minimal overlap (Figure 6E), indicating that differences in SHIV strains and/or infection routes strongly influence the host’s B-cell responses, leading to distinct foci – epitopic sites – of humoral dominance. The observation that distinct SHIV infections promote dissimilar humoral responses is likely critical to vaccine design strategy but we cannot rule out the possibility that our limited sample sizes could artifactually generate this divergence.

### Heterogeneous BCR genetics underlying similar functional profiles in genetically diverse RMs

Given the functional similarities of Env-reactive antibodies elicited in individual RMs by SHIV infection (Figure 6), it was important to determine the similarity of BCR repertoires between individual RMs, and to compare the genetics of Env-reactive BCRs relative to the naïve BCR repertoires in these SHIV-infected RMs. Does the phenotypic similarity of Env antibody responses equate to an equally similar usage of V(D)J gene segments or is the plasticity inherent in combinatorial genetic association sufficient for functional similarity to be generated by diverse genetic solutions? Answers to these questions shed light on the feasibility of an HIV vaccine to induce functionally similar antibody responses in genetically diverse human populations.

We extended a strategy from a recent study [66] to establish individualized RM IGV germline databases for both heavy and light chains. Multiplex primer sets were developed for rhesus IGHV, IGKV, and IGLV segment families (see Methods). Using the *IgDiscover* program [75], individualized IGV germline repertoires were determined via repertoire-sequencing (Rep-Seq) of MF B cells isolated from the six RMs in this study (Table S6). By these methods, we inferred 79–136 (average 114) IGHV, 98–111 (average 106) IGKV, and 70–103 (average 86) IGLV germline alleles from these RMs. The number of IGHV germline alleles inferred is comparable to that reported for five randomly selected Indian-origin RMs in a previous study [66] (74–105, average 95), suggesting a similar sequencing depth and recovery rate in our study.

Consistent with previous findings [65, 66, 75], the three RMs in each cohort are genetically diverse in germline repertoire, with 15.4–48.5% (average 30.6%) of IGHV, 27.0–34.6% (average 31.6%) of IGKV and 26.3–37.1% (average 33.5%) of IGLV germline alleles shared with the other two RMs in the same cohort (Figure 7A). When considering only Env-reactive BCRs, the shared allele percentages dropped further to 2.9–21.1% (average 11.8%), 2.8–27.8% (average 11.0%), and 10.8–25.0% (average 15.0%) for IGHV, IGKV, and IGLV, respectively (Figure 7B). This reduced allele sharing in Env-reactive BCRs raises the possibility at repertoire level that genetically diverse RMs utilize heterogeneous BCR genetics to achieve functionally similar antibody responses to the SHIV infection.

**Figure 7.**
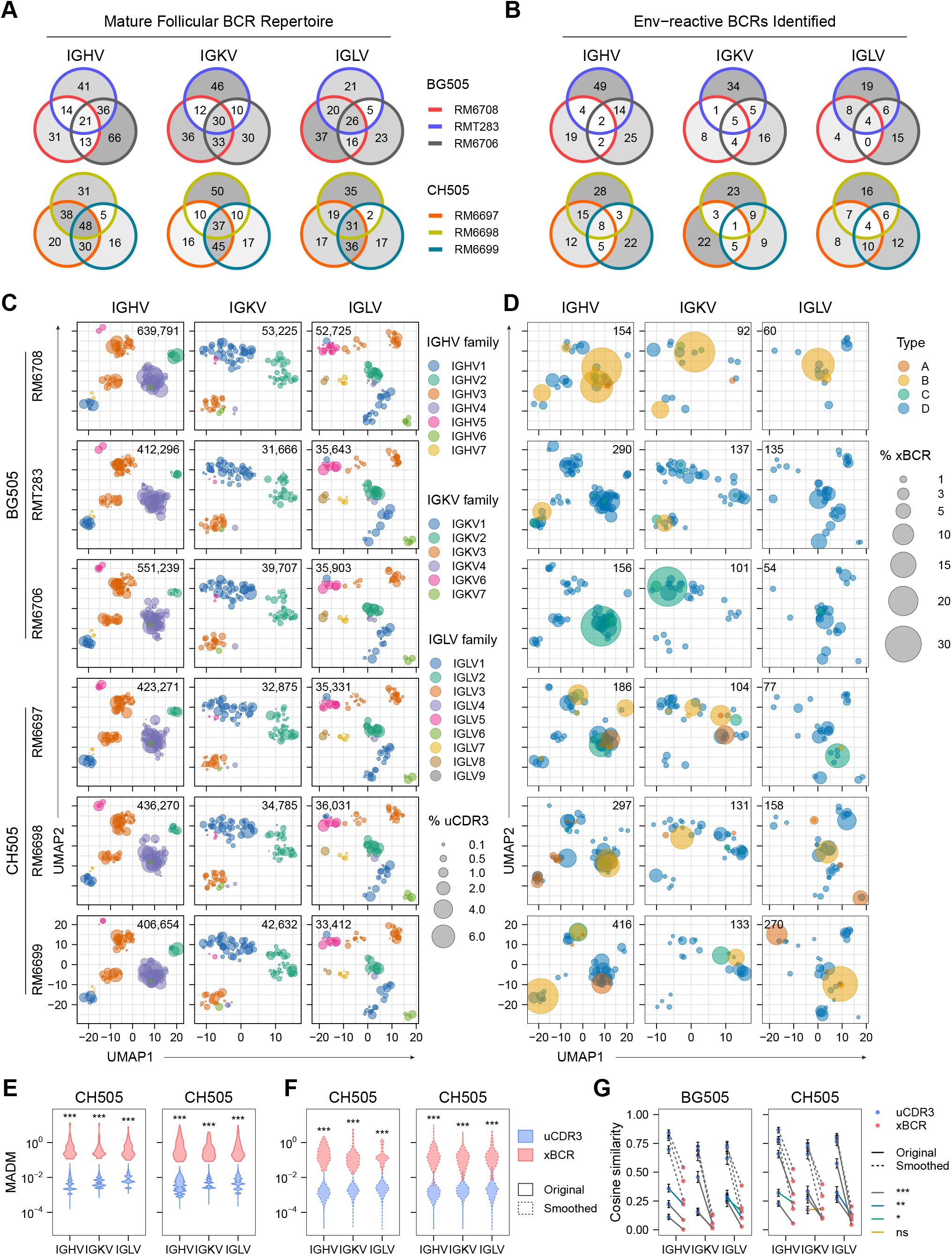
Heterogeneous BCR genetics underlying similar functional profiles in genetically diverse RMs. (A-B) Venn diagrams showing the numbers of shared and distinct IGHV, IGKV, and IGLV alleles in the MF BCR repertoire (A) and Env-reactive BCRs identified (B) across three RMs per infection group. (C-D) Bubble plots of IGV allele usage in the MF BCR repertoire (C) and Env-reactive BCRs (D), mapped to UMAP coordinates based on sequence similarity. Bubbles are color-coded by IGV family (C) or functional type (D) and scaled by usage percentages among unique CDR3s (uCDR3, C) or Env-reactive BCRs identified (xBCR, D). IGKV and IGLV are combined as light chain repertoire in percentage calculation. Numbers indicate counts of unique CDR3s from Rep-Seq (C) or Env-reactive BCRs identified (D). (E–F) Distribution of MADM of IGV allele usage frequencies across RMs, calculated from original percentages (E; solid lines) or percentages smoothed within IGV genes (F; dashed lines). The distribution for uCDR3 represents the combined MADM values from 1000 multinomial down-sampled simulations (see Methods). Empirical *p* values were calculated between the median MADM of xBCR and 1000 median values of randomly-sampled and size-matched uCDR3 subsets (see Methods). ***, empirical *p* < 0.001. Alleles with zero frequencies across all three RMs were excluded to prevent distortion by an excess of MADM = 0 values. (G) Cosine similarity scores of IGV allele usage between RMs. For the uCDR3 source, dots indicate medians from 1000 multinomial down-sampling simulations (see Methods), with error bars showing 25th – 75th percentiles. Lines connect the same RM pair comparisons from uCDR3 or xBCR sources, with solid lines for original percentages and dashed lines for percentages smoothed within IGV genes. Line colors indicate the significance of empirical *p* values: grey (***) for *p* < 0.001, blue (**) for *p* < 0.01, green (*) for *p* < 0.05, and orange (ns) for not significant. See also Table S6 and Figure S3.

We then sought evidence at the individual IGV allele level. Based on aligned IGV allele sequences, we constructed a distance matrix using the Kimura 2-parameter model [76] to represent similarity among alleles. We employed unified UMAP layouts, embedded with the similarity between individual alleles, to visualize IGV repertoires and compare IGV allele frequencies across different RMs. The percentages of unique complementarity-determining region 3 (CDR3) for each IGV allele in the MF BCR repertoire were plotted (Figure 7C). These UMAP plots reveal that the overall expression profiles of IGV families are generally similar among the analyzed RMs, although individual allele frequencies can vary. In contrast, Env-reactive BCRs exhibit distinct expression profiles among different RMs in each infection group (Figure 7D), particularly when different functional types are also considered.

To quantify variation in frequencies, mean absolute deviation from the median (MADM) values of the frequencies for individual IGV alleles were calculated among the three RMs in each infection group. To control for differences in sampling depth between MF BCR repertoires and Env-reactive BCRs, MF repertoires were down-sampled 1000 times using a multinomial distribution based on allele frequency probabilities derived from Rep-Seq data (see Methods). For each RM, the multinomial sample size was matched to the number of Env-reactive BCRs. Allele usage frequencies were then recalculated from the down-sampled repertoires. Compared with the size-matched MF BCR repertoires, Env-reactive BCRs exhibit significantly elevated MADM values for all IGV segment families in both infection groups (Figure 7E).

However, this analysis has a limitation. Due to the high levels of allelic variation of IGV genes in RMs [65], an allele-wise comparison might overestimate differences at the IGV gene level among RMs, given that closely related IGV alleles can produce BCRs with similar functions/specificities. To address this limitation, we devised a strategy to smooth the allele frequencies among IGV alleles (see Methods). First, using the sequence similarity-based allelic distance matrix established earlier, threshold values that best discriminate IGV genes were determined (Figure S3E). The frequencies for each individual IGV allele were then redistributed among all alleles within the threshold distance, weighted by the distance between them. This smoothed expression profile normalizes minor allelic variation and better reveals functional similarity among RMs. Accordingly, the MADM values decreased after smoothing for all IGV segment families in all experimental groups, with Env-reactive BCRs still exhibiting significantly higher variation than the MF BCR repertoire (Figure 7F). This trend holds even when smoothing is extended within IGV families (Figures S3F and S3G).

To assess the overall similarity in IGV expression profiles among RMs, the frequencies for individual IGV alleles from Env-reactive BCRs or from 1000 down-sampled MF repertoires, either as original values or after smoothing, were treated as multi-dimensional vectors. Cosine similarity scores [77] were calculated between every two RMs (see Methods). Except for IGKV for one RM pair in the CH505 infection group, these scores were significantly lower for Env-reactive BCRs compared to the MF BCR repertoires across all IGV segment families in both infection groups, regardless of whether the analysis used original frequencies or values after smoothing within IGV genes (Figure 7G) or families (Figure S3H).

To extend this analysis beyond IGV allele usage, we examined whether the shared functional response patterns were accompanied by similarities in HCDR3 length and joint IGHV-family, IGHJ-family, and HCDR3-length profiles. HCDR3 length distributions of Env-reactive clonotypes broadly overlapped those of the corresponding MF repertoires, although individual profiles varied among RMs (Figure S5A). To account for limited sampling, we compared observed repertoire distances with reference distributions generated from 1,000 MF subsamples matched to the number of Env-reactive clonotypes in each animal. Within-RM Wasserstein-1 distances between Env-reactive and MF repertoires fell within the central 95% reference intervals in five of six animals, with RMT283 exceeding the upper reference bound (Figure S5B). Between RMs infected with the same SHIV, five of six pairwise distances fell within their respective reference intervals, whereas the RM6708–RMT283 distance fell below the lower bound, indicating greater similarity in HCDR3 length distributions for this pair (Figure S5C). Analysis of joint IGHV-family, IGHJ-family, and HCDR3-length profiles using Jensen–Shannon distance yielded a similar within-RM pattern, with RMT283 again representing the sole animal whose distance exceeded the reference interval (Figure S5D). All six between-RM distances for these joint profiles fell within the corresponding reference intervals (Figure S5E). Thus, the shared functional response patterns were not accompanied by a consistent increase in between-animal similarity of these combined repertoire features beyond that observed with size-matched sampling of the MF repertoires.

## DISCUSSION

This is a systematic study of the B-cell responses during chronic SHIV infection in cohorts of genetically diverse RMs. The molecularly cloned T/F viruses underwent conserved mutation patterns or even shared identical amino acid substitutions in the Env gene (Figure 1), providing dynamic quasispecies [53] of Env antigens in real time. This infection model offers a straightforward model system to compare the similarity of humoral responses in genetically diverse RMs.

Antigen-unbiased BCR sampling is another strategic component enabling this study. Rather than using specific antigen probe for pre-selection, MF, IgM+ or IgG+ Bmem, and IgG+ GC B cells were sampled in an antigen-independent manner. Antigen specificity, binding strength, neutralization and BCR genetics analyses were subsequently based on these unbiased B-cell cultures. This approach allowed us to interrogate the evolution of Env-reactive B cells at the population level and to study systemically the similarity of B-cell responses in genetically diverse RMs elicited by identical SHIV infections.

### Dissimilarity in the magnitude and kinetics of B-cell responses

We compared the magnitude and kinetics of B-cell responses across different RMs at multiple levels, including the proportions of various B-cell populations, autologous neutralizing responses in GC B-cell populations, autologous SOSIP and gp140 binding GC B-cell responses (Figure 2), and the expansion/contraction of individual B-cell clones (Figure 3A). As expected, all these aspects varied among the outbred RMs. This infection model thus embraces individual differences as seen in human populations, and provides a natural basis to investigate similarities in BCR function and genetics.

### Similarity in the general functional properties of the B-cell responses

Despite variations in the magnitude and kinetics of humoral responses, the functional properties of the B-cell responses were similar between the different RMs infected with a common SHIV. In all RMs tested, GC responses persisted through all the timepoints analyzed and IgG^+^ GC B cells produced the majority of autologous neutralizing and SOSIP/gp140 binding antibodies after unbiased sampling and culture (Figure 2). The persistence and effective somatic evolution within GC responses lend support to the potential of serial vaccination to induce long-lasting GC responses that support and guide bnAb development.

In all RMs tested, autologous neutralizing GC B-cell clones tend to persist longer and undergo more extensive clonal expansions than their non-neutralizing counterparts (Figure 3). It would be interesting to determine whether neutralizing clones have higher precursor frequency and germline BCR affinities than non-neutralizing clones as reported [46, 78], or if other factors contribute to this selection process [79]. Uncovering the underlying mechanisms, which is advantageous from a vaccination perspective, presents an intriguing opportunity. Determining whether this selection is mediated by infection with replicating viruses or is intrinsic to the Env antigen itself is particularly interesting. If infection-dependent, exploring how vaccination can recapitulate this process would be of great interest.

In all RMs tested, the memory compartments consist of primarily diverse non-neutralizing Bmem cells that are clonally unrelated to neutralizing GC B cells (Figure 4). Future studies are needed to ascertain if this discrepancy indicates distinct selection mechanisms during primary responses [80, 81] or a preferential GC re-entry of neutralizing Bmem cells during recall responses, leading to a functionally biased Bmem repertoire. These results also highlight the necessity of unraveling the nuanced process of Bmem generation for the success of future vaccine designs that favor the generation of broadly neutralizing Bmem cells.

### Similarity in the antigen specificity and neutralization profiles of identified Env-reactive antibodies

Exploring the relationship between autologous neutralization and antigen binding, we identified four functional types of Env-reactive BCRs/Abs representing distinct antigen specificities (Figure 5). Detailed analysis of representative rAbs shows that BCRs/Abs from different RMs exhibit remarkable similarities within each functional type. These similarities span cross-clade gp140 antigen binding, heterologous neutralization, and epitope specificity clustering determined by competition binding assays (Figure 6). These findings underscore a deeper layer of functional convergence in B-cell responses across different RMs.

A notable observation is the conservation of non-neutralizing epitopes recognized by Type D and, to a lesser extent, Type C antibodies (Figure 6). These epitopes, present in cross-clade Env antigens and recognized across RMs, likely face minimal selection pressure from host humoral responses. Their similarity among RMs suggests a common viral strategy to divert immunity from neutralizing epitopes. Notably, Types D and C antibodies primarily contribute to heterologous neutralization against three specific tier 2 strains, highlighting the complexity of Env-mediated infection across HIV/SHIV strains. This raises questions about the pathophysiological significance of non-neutralizing antibody responses. Understanding these responses in the context of BCR-virus coevolution could inform vaccine design.

### Dissimilarity in the BCR genetics underlying functional similarities

In contrast to the functional similarities, Env-reactive BCRs employ heterogeneous genetics in these genetically diverse RMs (Figure 7). At IGV repertoire level, further diversified IGV allele usages were detected for Env-reactive BCRs as compared with MF BCR repertoire. At individual IGV allele level, Env-reactive BCRs exhibit distinct IGV allele expression profiles among different RMs, with significantly elevated variation in frequencies for individual alleles and consistently lower overall similarity in IGV expression profiles relative to that for MF BCRs. Extending these comparisons to HCDR3 length and joint IGHV-family, IGHJ-family, and HCDR3-length profiles revealed no consistent increase in between-RM similarity among Env-reactive clonotypes relative to size-matched MF subsamples (Figure S5), consistent with functional convergence across genetically diverse BCR repertoires.

These observations contrast with the emphasis on biased IGHV gene usage in previous studies of HIV Env-reactive antibodies [82–84]. The germline-targeting or lineage-based vaccine strategies for certain bnAb classes, most notably CD4bs bnAbs such as VRC01-and CH235-like antibodies, are also largely based on the assumption that specific V(D)J usage or certain CDR3 properties are prerequisites for defining a bnAb germline precursor. The presence of “public epitopes” may explain shared germline usage in some cases [85]. However, the proportion of BCRs recognizing public epitopes within the Env-reactive repertoire, especially those leading to bnAb responses, remains to be determined.

Even in HIV vaccine studies, multiple observations challenge restricted V(D)J usage and support BCR genetic diversity toward a given epitope. In VRC01 germline knock-in mice immunized with germline-targeting immunogen eOD-GT8, only 3.2% of CD4bs-specific IgG BCRs are VRC01-class [86]. Similarly, in a clinical trial with eOD-GT8 [87], VRC01-class B cells constituted a minority among CD4bs-specific IgG^+^ B cells, whereas non-VRC01-class B cells were highly polyclonal, exhibited diverse V(D)J usage and CDR3 lengths, and underwent more efficient affinity maturation in GCs. In RMs immunized with N332-GT5, targeting the V3 glycan-specific bnAb BG18, an antibody lacking BG18 sequence features, except for a long heavy-chain CDR3, bound the immunogen in a highly homologous manner [88]. Likewise, in humans immunized with the MPER peptide, a polyclonal, heterogeneous BCR repertoire with neutralizing activity was induced, dominated by an IGHV gene not previously described for MPER-specific bnAbs [40].

The shared patterns of epitope recognition among genetically diverse Env-reactive B-cell receptors observed here are conceptually analogous to ecomorph evolution [89], in which genetically distinct populations independently evolve similar phenotypes in response to shared ecological constraints. In the SHIV infection model studied here, infection of genetically diverse RMs with genetically identical SHIV strains within each infection group gives rise to temporally dynamic Env antigenic environments that exhibit convergent mutation patterns across hosts. Within this shared antigenic context, independently expanding B-cell clones exhibit convergent functional recognition despite heterogeneous underlying BCR gene rearrangements, resulting in functionally similar yet genetically distinct Env-reactive B-cell populations. This conceptual analogy emphasizes the epitope itself as a possible shared selective constraint rather than the specific genetic solutions by which individual BCRs achieve recognition. While the present study does not address the elicitation of bnAbs, these observations provide a conceptual framework for considering how similar principles might be explored in vaccine design.

### Implications for vaccine design

The functional convergence of Env-reactive BCRs with heterogeneous genetics in genetically diverse RMs may be encouraging for vaccine development. Our results suggest that individual differences in BCR genetics may not be a barrier to a generalizable vaccine. These observations also suggest that specific germline-targeting may not be an absolute prerequisite for a successful vaccine for all bnAb epitopes. Given the limited immunogenicity of many bnAb epitopes [5, 90], immunogen design might focus on enhancing the immunogenicity of bnAb epitopes themselves at the population level, while allowing the immune system of individual vaccinees to engage these epitopes using V(D)J and CDR3 configurations permitted by the structural constraints of the epitope, rather than specifically targeting one or a few known germline BCRs capable of somatic evolution to bnAb activity. One possible approach is to use a rationally or empirically designed immunogen bearing one or more bnAb epitope(s) engineered to enhance their intrinsic immunogenicity across genetically diverse populations, potentially in combination with strategies that limit the immunodominance of non-neutralizing epitopes. Sequential immunizations with the immunogens evolving towards the native epitopes may then guide humoral responses toward bnAb epitopes present in circulating HIV strains. Recent studies by our laboratories using genetically-engineered, epitope-focused Env trimers expressed by replicating SHIVs, ferritin nanoparticles or mRNA-lipid nanoparticles to induce potent bnAbs targeting V2 apex and V3 glycan epitope supersites support this approach [91, 92]. State-of-the-art technologies, including deep mutational scanning [93] and artificial intelligence-driven biomolecular interaction prediction [94], could further aid immunogen design. Insights from public epitope studies [85] could help engineer bnAb epitopes into “public epitopes” for priming. Such bnAb epitope-focused Env designs may also extend to vaccines for other challenging pathogens, including coronaviruses, influenza, and hepatitis C viruses.

### Limitations of this study

As noted earlier, the infection model offers a simpler way to generate dynamic Env quasispecies in real time. However, the complexity of host-virus interactions and the uncontrolled evolution of viral antigens make it difficult to compare BCR genetics across different RMs responding to the same temporarily static immunogen. Future studies, informed by the insights gained from this work, should use serial immunization to explore improved vaccine design strategies. Although SHIV-infected RMs can recapitulate features of human Env–antibody coevolution and bnAb maturation [51, 52], the BG505 and CH505 Envs were originally derived from human infections and are being evaluated in a different host immune repertoire. Species differences in Ig germline repertoires and precursor availability may influence which BCRs are recruited, and we cannot exclude the possibility that these human-derived Envs more readily elicit neutralizing antibodies in RMs than in humans. This potential mismatch limits extrapolation to human vaccine responses, and the extent to which the repertoire-level patterns observed here extend to humans remains to be determined.

Another limitation is the lack of plasma cell and plasmablast analysis due to the technical constraint that our single B-cell culture system cannot support their culture. While the single B-cell RT-PCR, V(D)J cloning, and rAb production workflow allows for a detailed interrogation of these populations, this workflow is subject to limited throughput based on state-of-the-art techniques. The development of a high-throughput workflow is underway and is beyond the scope of this study.

A final limitation is that only a small number of potential bnAbs are represented in our overall Env-induced datasets. Unlike strain-restricted neutralizing antibodies, bnAbs retain neutralizing activity across diverse HIV strains, which may impose additional genetic and maturation requirements. Whether the genetic flexibility observed here extends to bnAb development therefore remains to be determined. A key question for future studies is how to recruit and expand rare bnAb precursors and guide their maturation toward breadth in the presence of competing non-bnAb responses.

### Technical innovations in this study

RMs are essential preclinical models, yet methods for characterizing antigen-specific B cells and immunogenetics remain underdeveloped [65, 66]. In this study, we established or refined key techniques for studying RM humoral responses, including a feeder cell line for high-throughput B-cell culture and cloning (Figure S1), a versatile and unbiased BCR V(D)J sequencing platform, and multiplex primers for establishing individualized RM Ig heavy and light chain germline databases (see Methods). These techniques, usable individually or in combination, could greatly enhance other studies of humoral responses using RMs as a model system. The engineered feeder cell line, MEC-147, also supports efficient single human B-cell cloning (Figure S1), representing a streamlined method for fully human antibody discovery and potentially useful for the identification of diagnostic and therapeutic antibodies in industry.

## RESOURCE AVAILABILITY

### Lead contact

Requests for further information, resources, and reagents should be directed to the lead contact, Garnett Kelsoe.

### Materials availability

The feeder cell line MEC-147 is available for non-commercial use under a Material Transfer Agreement (MTA) and for commercial use under a license. All other materials from this study are available from the lead contact upon reasonable request.

### Data and code availability

V(D)J sequences derived from the 146 Env-reactive BCRs selected for rAb production and characterization have been deposited in NCBI GenBank under accession numbers PV497204–PV497495. V(D)J sequences and associated metadata for all 1,499 Env-reactive BCRs identified in this study are provided in Table S3. SHIV Env genomic sequences were deposited previously [51, 95] in NCBI GenBank under accession numbers MN468576–MN468585, MN468748–MN468751, MN468914–MN468930, MN469235–MN469265, MN469516– MN469531, MN469591–MN469592, MT486140–MT486182, MT486279–MT486282, MT486349–MT486358, MT486603–MT486624, MT486654–MT486672, and MT486741–MT486762. BCR Rep-Seq raw data have been deposited in the NCBI Sequence Read Archive (SRA) under accession numbers SRX28288042– SRX28288065 for 5’RACE libraries and SRX28339869–SRX28339904 for multiplex (MTPX) libraries (BioProject: PRJNA1247978). The source code for the C# program *AbSolute* is available on GitHub at https://github.com/shengli-song/AbSolute. All source data and custom R and Python scripts used for data processing and figure generation are available via Zenodo under DOI 10.5281/zenodo.23606983.

## ACKNOWLEDGMENTS

We thank Xiaoyan Nie, Xiaoe Liang, Dongmei Liao, Nikita Hall, Yunhan Jiang, and Jiaying Garnett for technical assistance in B-cell related assays. We thank Dr. Cliburn Chan for reviewing the manuscript and providing advice on data analysis. We thank Joel Finney for insightful discussion. We thank Jiayu Chen, Nathan Newton, Alice Kimbell, Shi-Mao Xia, and Peter Gao for technical assistance and management in neutralization assays. We thank Steven Slater and Amanda Foreman for technical assistance in flow cytometry. We thank Thad Gurley, Yousef Abuahmad, and Lawrence Armand for technical assistance and management in lymph node biopsy processing, sample allocation, and Env antigen conjugation. We thank Ashley Trama in Env antigen preparation. We thank Nicolas Devos and Graham Alexander Jr for technical assistance in MiSeq sequencing. This study was supported by NIH grants AI128832 (G.K.), AI131251 (G.M.S.) and AI100148 (G.M.S.).

## AUTHOR CONTRIBUTIONS

G.K. and S.S. design the research. S.S., H.L., H.G., X.S. and C.C.L. performed the research. C.Y., A.W., and M.K. provided crucial assays. W.B.W., K.O.S. and M.A.M. provided crucial materials. S.S., H.L., H.G., X.S., C.C.L., K.W., D.C.M., G.M.S., and G.K. analyzed the data. S.S. visualized the data and drafted the manuscript. G.K., G.M.S., B.F.H, D.C.M., K.W., and K.O.S. edited and revised the manuscript. G.K. directed the study.

## DECLARATION OF INTERESTS

The authors have the following disclosures. Garnett Kelsoe, Shengli Song, and Duke University, non-exclusively licensed the engineered feeder cell line MEC-147 to *Moderna Inc*. Royalties from this license were used to fund this work as reported here. All other authors have no declarations.

### Supplemental Figures

**Figure S1.**
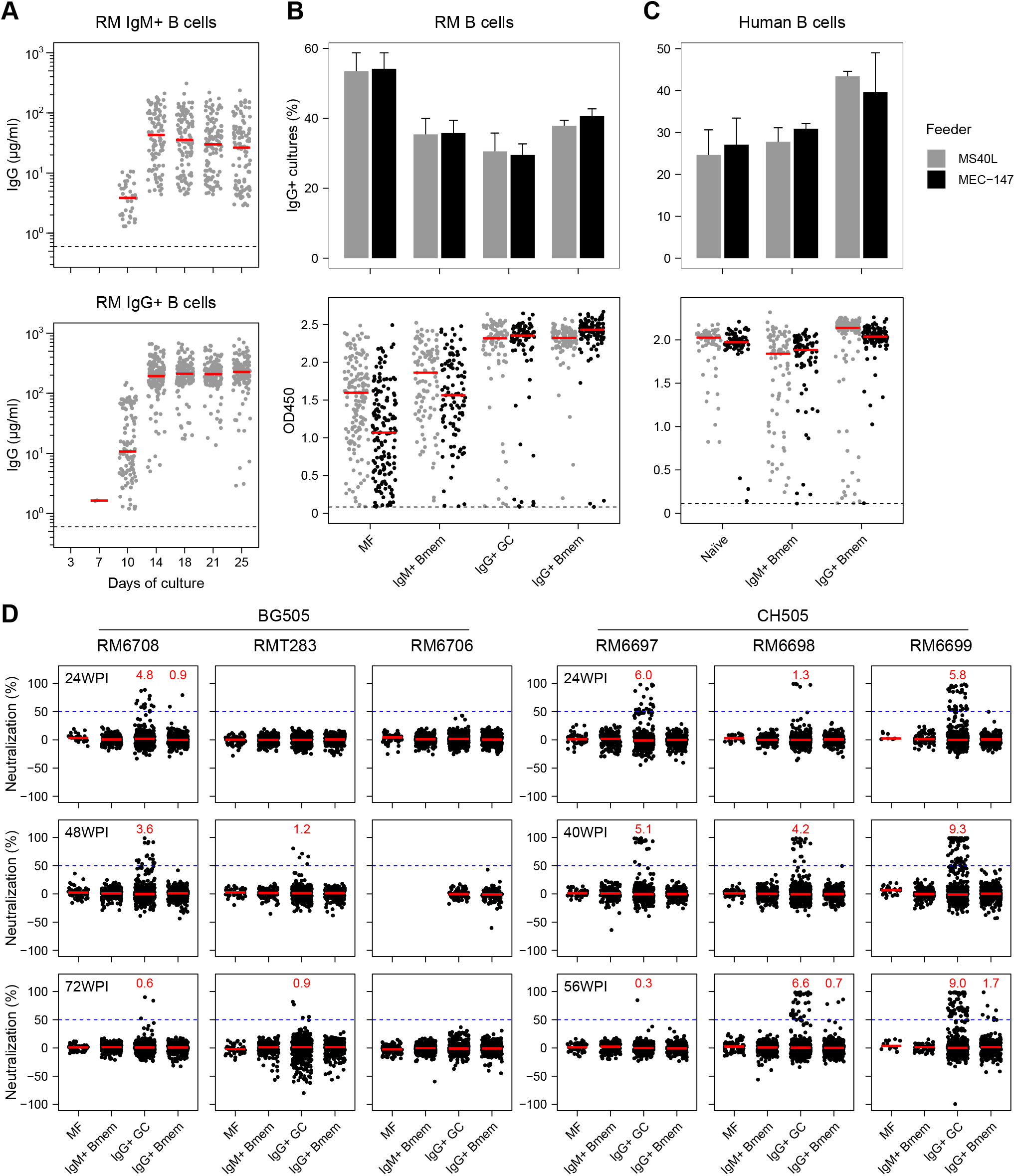
Feeder cell lines for single B-cell cultures, and autologous neutralization activities from the culture supernatants, related to Figure 2. (A) Kinetics of IgG production in culture supernatants from RM IgM^+^ and IgG^+^ single B cells cultured with the MS40L feeder cell line. Each dot represents an individual IgG^+^ culture well. Dashed lines indicate detection limits, and error bars (in red) represent the median values. (B-C) Comparison of the original MS40L and engineered MEC-147 feeder cell lines in supporting single B-cell cultures from different B-cell populations. Panel (B) shows the culture of B cells isolated from the lymph nodes of an RM after DTaP immunization, while panel (C) shows the culture of B cells isolated from PBMCs of a healthy human donor. The top panels display the percentage of IgG^+^ culture wells among all single-cell sorted wells. Error bars indicate the mean + SD across three 96-well culture plates. The bottom panels show OD450 values from IgG ELISA assays, with each dot representing an individual IgG^+^ culture well. Dashed lines indicate detection limits, and error bars (in red) represent the median values. (D) Single nucleated cells were isolated from LN biopsies of infected RMs at various time points following SHIV infection. B cells of different phenotypes, including mature follicular (MF), IgM^+^ Bmem, IgG^+^ GC, and IgG^+^ Bmem cells, were identified from isolated LN cells by flow cytometry and sorted into single B-cell cultures with MEC-147 feeder cells. Supernatants were harvested after 18 days of culture, and IgG-producing cultures were identified using standard ELISA assays. IgG-containing culture supernatants were tested for neutralization activity against autologous SHIV pseudo-viruses using standard TZM-bl assays. Each dot represents an individual single B-cell culture supernatant containing clonal IgG. Error bars in red indicate median values, while blue dashed lines indicate cutoff thresholds for defining neutralizing versus non-neutralizing cultures. Inserted red numbers represent the percentage of neutralizing cultures among IgG^+^ cultures for the corresponding B-cell populations (see Table S2 for details).

**Figure S2.**
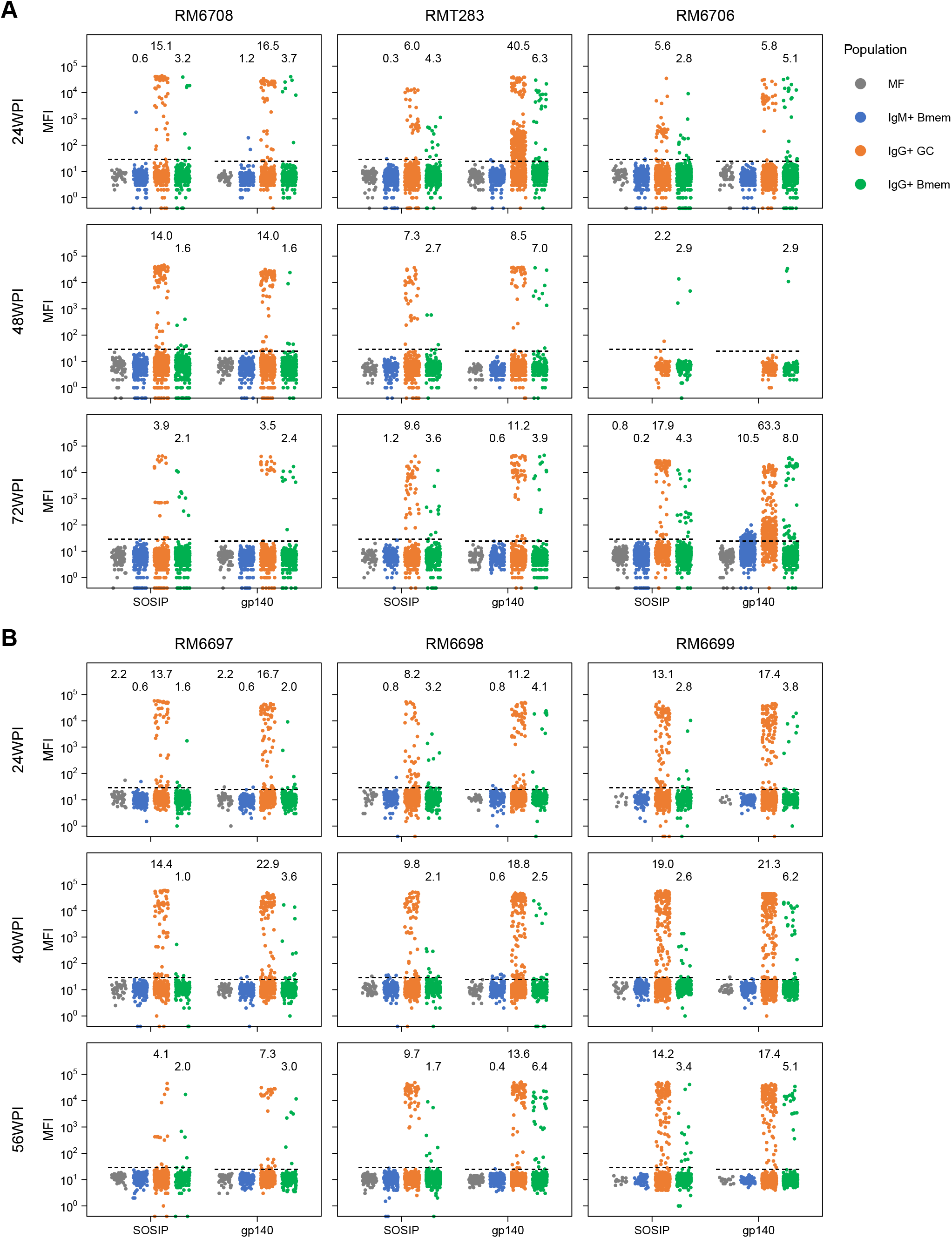
Autologous Env antigen-binding activities of single B-cell culture supernatants, related to Figure 2. The same IgG-containing single B-cell culture supernatants prepared as described in Figure S1D were tested for binding activity against autologous SOSIP and gp140 antigens using standard Luminex assays, conducted in parallel and in a blinded manner to the autologous neutralization assays shown in Figure S1D. RMs were infected with either the BG505 (A) or CH505 (B) SHIV strains. B-cell populations are color-coded as indicated in the keys. Dashed lines indicate cutoff thresholds for defining antigen-binding versus non-binding cultures. Inserted numbers represent the percentage of antigen-binding cultures among IgG^+^ cultures for the corresponding B-cell populations. Additional antigens or capture antibodies, including CH505.gp120, MN.gp41, anti-RhIgG, and non-relevant controls (streptavidin, BSA, and ovalbumin), were included in the Luminex bead panel (see Table S2 for details).

**Figure S3.**
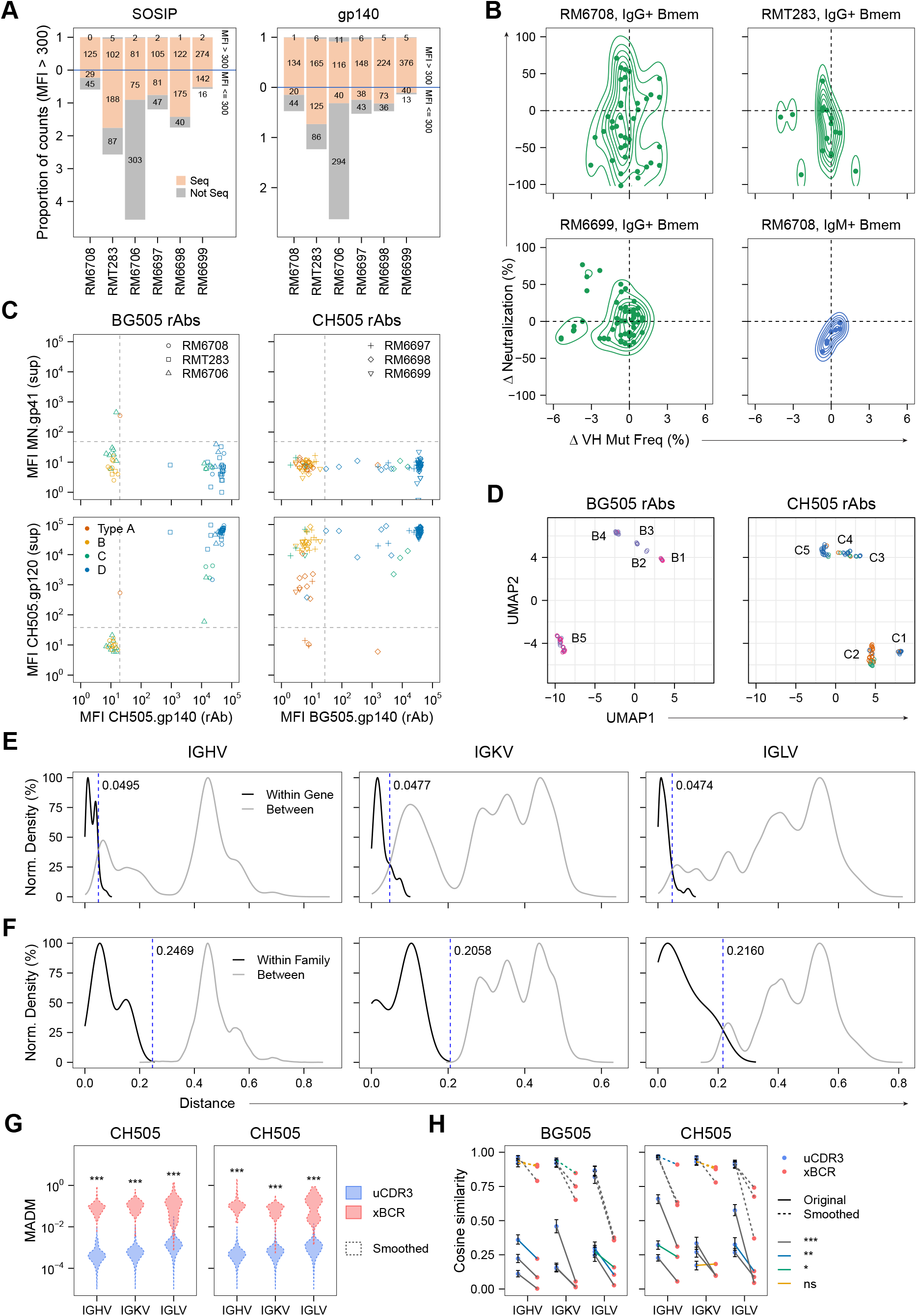
Verification and supporting analyses, related to Figures 4, 6, and 7, Table S3, and Methods. (A) Related to Table S3. Proportions of identified autologous SOSIP-or gp140-binding B cells for which BCR sequences were determined. The blue horizontal line separates two categories: cultures with binding MFI > 300 (upper) and MFI ≤ 300 (lower). Stacked bars represent proportions normalized to the total counts of Env-binding B cells with MFI > 300, with pink indicating sequenced and grey indicating not sequenced. Numbers within bars denote the counts of Env-binding B cells in each category. (B) Related to Figure 4E. VH mutation frequency and neutralization differences between Bmem and IgG^+^ GC B cells for lineages containing both IgG^+^ GC and Bmem members at the same timepoints in individual RMs. Each dot represents a Bmem–IgG^+^ GC B cell pair. Only RM–Bmem population groups with more than two data points are shown. (C) Related to Figure 6A. Relationship between rAb binding MFI values to heterologous gp140 protein and the binding MFI values of corresponding single B-cell culture supernatants to CH505.gp120 and MN.gp41 proteins in the initial screening. Functional types of source B-cell clones are color-coded, and dashed lines indicate binding cutoffs. (D) Related to Methods. UMAP plots showing the same coordinates and clusters as in Figure 6E, with colors indicating individual assay batches. (E-F) Related to Figure 7. Normalized density plots display the distribution of distances calculated from the pairwise IGV gene distance matrix at two categorical levels: IGV gene (E) and family (F). Distances within the same category (“Within”) are shown in black, while distances between different categories (“Between”) are shown in grey. Dashed vertical lines in blue represent cutoff values, determined either at the intersection of within and between distance distributions or at the point of minimum difference when no intersection is observed. (G) Related to Figure 7. Distribution of MADM of IGV allele usage frequencies across RMs, calculated from percentages smoothed within IGV families. The distribution for uCDR3 represents the combined MADM values from 1000 multinomial down-sampled simulations (see Methods). Empirical *p* values were calculated between the median MADM of xBCR and 1000 median values of randomly-sampled and size-matched uCDR3 subsets (see Methods). ***, empirical *p* < 0.001. Alleles with zero frequencies across all three RMs were excluded to prevent distortion by an excess of MADM = 0 values. (H) Related to Figure 7. Cosine similarity scores of IGV allele usage between RMs. For the uCDR3 source, dots indicate medians from 1000 multinomial down-sampling simulations (see Methods), with error bars showing 25th – 75th percentiles. Lines connect the same RM pair comparisons from uCDR3 or xBCR sources, with solid lines for original percentages and dashed lines for percentages smoothed within IGV families. Line colors indicate the significance of empirical *p* values: grey (***) for *p* < 0.001, blue (**) for *p* < 0.01, green (*) for *p* < 0.05, and orange (ns) for not significant.

**Figure S4.**
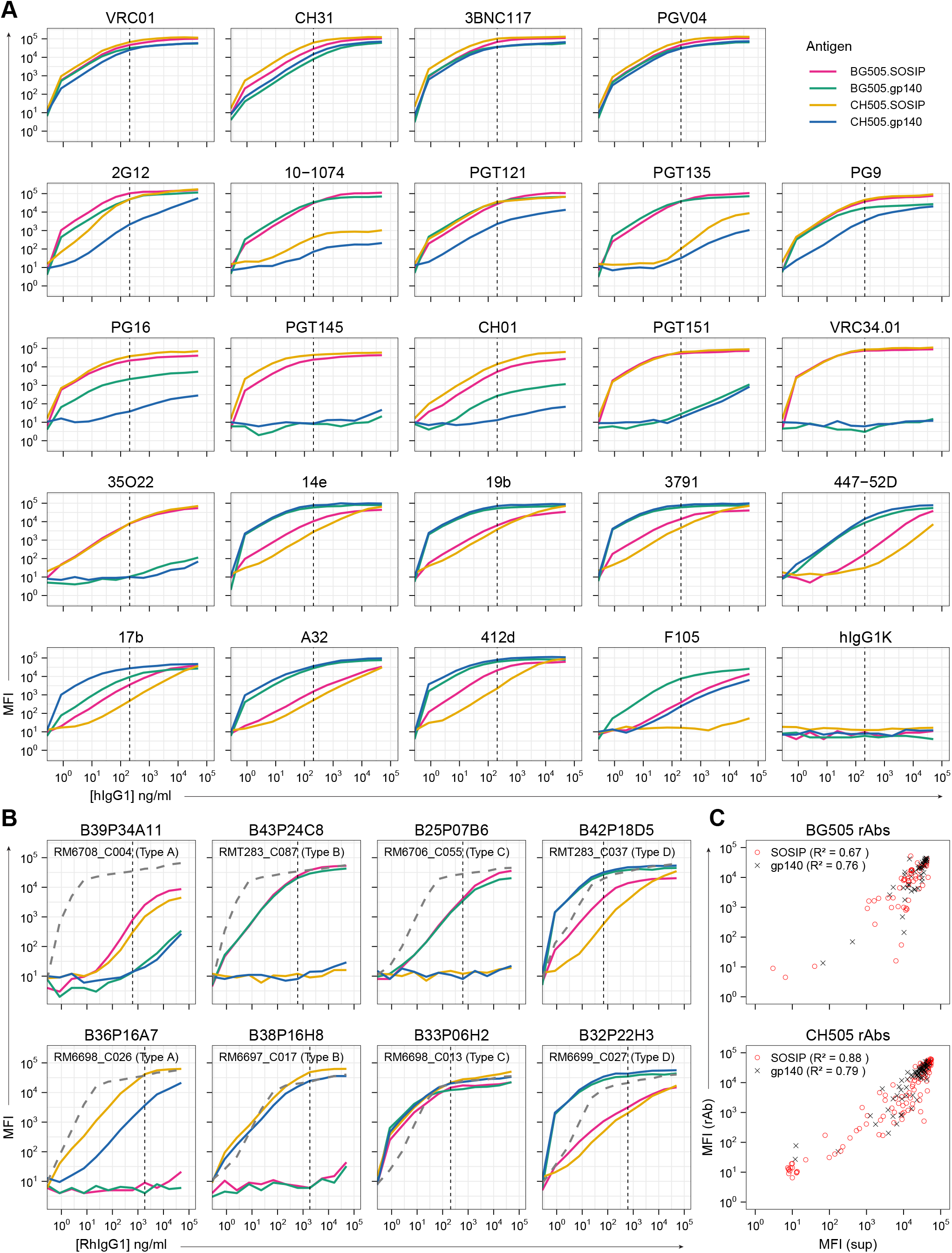
Env antigen-binding activities of rAbs, related to Figures 5 and 6. (A) Binding activities of reference bnAbs and nnAbs (human IgG1 format) for BG505 and CH505 SOSIP and gp140 antigens were tested in standard Luminex assays using serial dilutions. Vertical dashed lines indicate the concentration point (206 ng/ml) used for calculating MFI ratios in Figure 5A. (B) Binding activities of rAbs (rhesus IgG1 format) derived from representative B-cell clones of different functional types were similarly tested. Results for selected rAbs from BG505-infected (top row) and CH505-infected RMs (bottom row) are shown (named as “RM_Clone-ID (Type)”). Full data for all 146 rAbs are provided in Document S5. Antigens are color-coded as in panel (A). Dashed grey curves represent MFI values detected with Luminex beads conjugated to an anti-RhIgG capture antibody. Vertical dashed lines indicate the concentration points where anti-RhIgG MFI values align closely with those detected using culture supernatants during initial screening. (C) Comparison of MFI values for autologous SOSIP and gp140 antigens detected using culture supernatants (MFI (sup)) during initial screening and rAbs (MFI (rAb)) at selected concentration points (indicated by vertical dashed lines in panel (B) and Document S5). Linear regression models were fitted, and R^2^ values indicating the proportion of variance explained are shown in parentheses in the legends for corresponding antigens.

**Figure S5.**
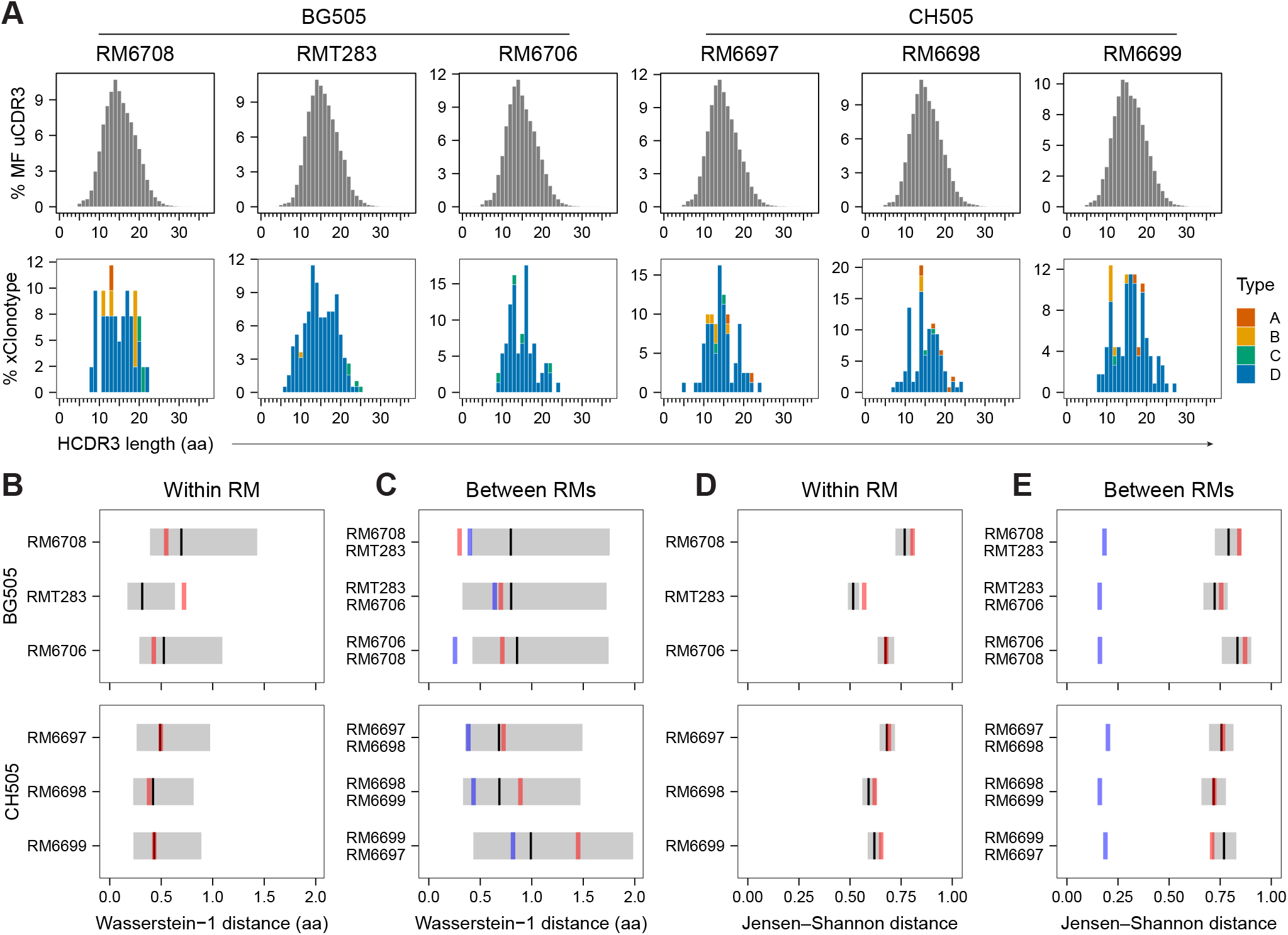
Comparison of HCDR3 length distributions and joint IGHV–IGHJ–HCDR3 profiles in MF and Env-reactive BCR repertoires, related to Table S8. (A) HCDR3 amino acid length distributions in MF B-cell repertoires (upper row, grey) and Env-reactive BCR clonotypes (xClonotypes; lower row) in individual RMs. MF sequences were deduplicated by HCDR3 nucleotide sequence and V/J assignment (uCDR3s), and Env-reactive BCRs were counted once per heavy-chain clonotype. Histograms show percentages within each repertoire. Stacked colors indicate functional types. (B-C) Within-RM (B) and between-RM (C) comparisons of HCDR3 length distributions using Wasserstein-1 distance, expressed in amino acid units. In B, red lines indicate distances between each RM’s xClonotype and complete MF repertoires. In C, red and blue lines indicate distances between xClonotype repertoires and between complete MF repertoires, respectively, for pairs of RMs within the same infection group. To account for sampling variability, 1,000 random subsamples were drawn without replacement from each RM’s MF repertoire, each containing the same number of entries as that animal’s xClonotype repertoire. Reference distances were calculated between each MF subsample and its corresponding complete MF repertoire in B, and between MF subsamples from the two animals in C. Grey bars indicate the 2.5^th^–97.5^th^ percentiles of the resulting distance distributions, and black lines indicate their medians. (D-E) Within-RM (D) and between-RM (E) comparisons using the same repertoires, subsampling strategy, and graphical conventions as in B and C, respectively. Here, repertoire profiles were defined by the frequencies of joint IGHV-family, IGHJ-family, and HCDR3-length combinations and compared using Jensen–Shannon distance. For both distance metrics, smaller values indicate greater repertoire similarity.

## SUPPLEMENTAL INFORMATION

**Document S1. Annotated Env amino acid sequence alignment from BG505-infected RMs**.

Env sequences (n = 74) recovered from six RMs at 88 WPI are aligned with the original Env sequences used to produce the BG505 SHIV for infection as references. Sequences are grouped by RM, with sequence identifiers, GenBank accession numbers, and sample sizes indicated. The alignment spans HXB2 positions 1– 709, with Env regions annotated above the alignment. Dots indicate identity to the reference; red letters indicate substitutions, blue letters indicate inserted residues, and orange dashes indicate deletions. Gray dashes indicate gaps in insertion columns, and X denotes unresolved residues. Asterisks in the numbering row identify insertion columns without assigned HXB2 positions. Identical sequences are retained as separate rows.

**Document S2. Annotated Env amino acid sequence alignment from CH505-infected RMs**.

Env sequences (n = 105) recovered from six RMs at 24, 48, or 56 WPI are aligned with the original Env sequences used to produce the CH505 SHIV for infection as references. The layout and annotations are as described for Document S1.

**Document S3. Phylogenetic trees of identified lineages, related to Figure 3**

BCR sequences were determined from identified autologous neutralizing and/or Env-binding B cells. V(D)J usages were assigned based on individualized germline databases, and clonal relationships were determined. Germline sequences were inferred, and phylogenetic trees were reconstructed. The V(D)J genetics were integrated with time-course, phenotypic, and functional data. Phylogenetic trees incorporating these metadata were visualized using the R package ggtree. The inferred germline sequences are represented as gray-filled circles (●) at the root of the trees, while identified sequences are shown as circles color-filled according to autologous neutralization activities. Squares (▪) and diamonds (♦) to the right of the circles are color-filled based on binding activities to autologous SOSIP and gp140 antigens, respectively. Sequence names are color-coded by the timepoints (WPI) at which they were identified. Sequences derived from IgM^+^ Bmem (◄) or IgG^+^ Bmem (◄◄) populations are indicated, while the remaining sequences originate from the IgG^+^ GC population.

**Document S4. Autologous neutralization and SOSIP– and gp140-binding activities for all Env-reactive B-cell lineages identified, related to Figure 5**

Dot plots illustrate the autologous SOSIP-binding versus gp140-binding MFI values for individual Env-reactive B-cell clones identified from each RM. Each dot represents an Env-reactive B cell identified, color-coded with a gradient to indicate autologous neutralization activities and shape-coded to denote the timepoint (WPI) of its identification. B-cell clones are labeled as “RM_Clone-ID,” with functional types specified in parentheses.

**Document S5. Env antigen-binding activities for all 146 rAbs, related to Figures 6 and S3**

Rhesus IgG1 rAbs derived from representative B-cell clones (labeled as “RM_Clone-ID”) of various functional types (specified in parentheses) were produced and tested for their binding activities to BG505 and CH505 SOSIP and gp140 antigens. Binding activities were assessed using standard Luminex assays with serial dilutions. Antigens are color-coded, and dashed grey curves represent MFI values detected using Luminex beads conjugated to an anti-RhIgG capture antibody. Vertical dashed lines indicate the concentration points where anti-RhIgG MFI values align closely with those detected using culture supernatants during initial screening.

**Table S1. Summary of B-cell phenotyping and functional screening data after single B-cell cultures, related to Figure 2**

For each RM in each SHIV infection group and at each timepoint (WPI), the percentages of the four B-cell populations among CD20^+^ B cells are presented. The table includes the total number of single B-cell cultures set up with each B-cell population, the number of IgG^+^ cultures, and the counts of cultures identified as autologous neutralizing, Env protein-binding, autologous SOSIP-binding, or gp140-binding, derived from IgG^+^ cultures.

**Table S2. Autologous neutralization and antigen-binding screening data with 17,259 individual IgG-containing single B-cell culture supernatants, related to Figure 2**

For each RM in each SHIV infection group and at each timepoint (WPI), four B-cell populations were identified by flow cytometry and sorted into single B-cell cultures. IgG-containing culture supernatants were screened for neutralization activities using standard TZM-bl assays for autologous SHIV pseudo-viruses and for binding activities against autologous SOSIP and gp140 proteins using standard Luminex assays. Additional proteins or capture antibodies, including CH505.gp120, MN.gp41, anti-RhIgG, and non-relevant controls (streptavidin, BSA, and ovalbumin), were included in the Luminex bead panel. Neutralization activities and binding MFI values are listed. Rows highlighted in yellow indicate batch-specific cutoff thresholds that define neutralizing versus non-neutralizing and binding versus non-binding cultures.

**Table S3. Combined phenotyping, functional, and V(D)J genetics data for 1499 neutralizing and/or Env antigen-binding B cells identified, related to Figures 3, 4, 5 and 6**

The V(D)J sequences were determined for 1499 neutralizing and/or Env antigen-binding B cells. The V(D)J genetics data were integrated with corresponding timepoint, phenotyping, and functional data as detailed in Table S2. Column titles are shaded to represent data categories: timepoint and phenotyping data in grey, functional data in light blue, heavy chain genetics in light green, light chain genetics in orange, and genetics of a second light chain (if applicable) in pink. Functional types, determined through combined analysis of neutralization, SOSIP– and gp140-binding activities, and heavy chain clonal data, are listed under a column with a purple-shaded title. The IGHV and IGLV columns list IGV allele calls based on individualized germline databases generated using the *IgDiscover* program. Unique IGV alleles, excluding duplicated calls with variations only in terminal sequence lengths, are listed in the uni-IGHV or uni-IGLV columns.

**Table S4. The neutralization breadth of produced rAbs, related to Figure 6**

The produced rAbs were evaluated at specified concentrations in standard TZM-bl assays against a panel of 11 heterologous tier 2 pseudo-viruses and the autologous SHIV strain used for initial infection. Neutralization percentages for each rAb across the pseudo-virus panel are listed. Columns with titles shaded in grey display meta-information for the rAbs, derived from Table S3. SVA-MLV, a negative control virus pseudotyped with murine leukemia virus envelope, was included in the assays to confirm assay specificity.

**Table S5. Inhibition of analyte rAbs binding to autologous Env antigens by competitor antibodies, related to Figure 6**

Binding activities of produced rhesus IgG1 rAbs (analytes, shaded in orange) to autologous SOSIP and gp140 (antigens, shaded in grey) were measured in the presence of various human IgG1 bnAbs and nnAbs (competitors, with column titles shaded in light blue) of known specificities. The averaged binding MFI values from two duplicated wells with labeling buffer only and without competitor antibodies were set as baseline (with column titles shaded in light green) for calculating inhibition. A cutoff threshold (with column titles shaded in yellow) was established to exclude analyte rAbs with maximum MFI values across all assay wells below 1000 from the analysis. The percentages of inhibition (with column titles shaded in pink) were calculated for each competitor antibody relative to the baseline MFI values.

**Table S6. Individualized IGV germline repertoires of the six RMs involved, related to Figure 7**

Using the *IgDiscover* program, individualized germline repertoires for IGHV, IGKL, and IGLV were determined through Rep-Seq analysis of MF B cells isolated from the six RMs involved in this study. IGV alleles assigned by the *IgDiscover* program are listed in the column labeled “V_allele”. Unique IGV alleles, excluding duplicated entries differing only in terminal sequence lengths, are presented in the column labeled “uni_V_allele”, with the longest germline sequences recorded in the column labeled “uni_seq”. Corresponding columns provide the total number of reads (n_reads) and the number of unique CDR3s (uniCDR3) identified for each unique IGV allele. The percentage of unique CDR3s was calculated for each unique IGV allele out of the total unique CDR3s identified within a given RM, with IGKV and IGLV combined for the light chain calculation.

**Table S7. Primers used in BCR V(D)J sequencing and Rep-Seq, related to Methods**

Primers are listed with the column “Purpose” indicating their use in BCR V(D)J sequencing and 5’ RACE (Rep-Seq, 5’ RACE) or MTXP (Rep-Seq, MTXP) library preparation for Rep-Seq. Reference 1: Vazquez Bernat, N., et al., Rhesus and cynomolgus macaque Ig heavy-chain genotyping yields comprehensive databases of germline VDJ alleles. *Immunity*, 2021. 54(2): p. 355-366 e4.

**Table S8. IGHV and IGHJ assignments and HCDR3 lengths for MF and Env-reactive BCR repertoire comparisons, related to Figure S5**

The “uCDR3_RM” worksheets contain MF Rep-Seq entries from individual RMs, deduplicated by IGHV assignment, IGHJ assignment, and HCDR3 nucleotide sequence. The “xClonotypes” worksheet was derived from Table S3 and contains one entry per Env-reactive heavy-chain clonotype, with each feature represented by its most frequent non-missing value within that clonotype.

## METHODS

### Animals and study design

Indian-origin rhesus macaques (*Macaca mulatta*), aged 3–6 years, both male and female, were used in this study. Animals were housed at *Bioqual Inc*. in accordance with guidelines of the Association for Assessment and Accreditation of Laboratory Animal Care (AAALAC). The study design and all experimental procedures were approved by the University of Pennsylvania and Bioqual Institutional Animal Care and Use Committees. All animals tested negative for Mamu-A*01, B*08, and B*17, had not previously been immunized with HIV proteins or infected with SHIVs, and were assigned to experimental cohorts prior to SHIV infection without formal randomization due to practical constraints inherent to non-human primate studies. All animals were sedated for blood draws, biopsies, and SHIV inoculations.

Two cohorts of six RMs were infected with molecularly cloned strains of SHIVs that express the Env glycoproteins of T/F HIV-1 virus strains, BG505.T332N (BG505 for short, Clade A) or CH505 (Clade C). All BG505-infected RMs were inoculated via the intrarectal route. The infection doses corresponded to 1.5 ng p27 antigen for RM6708, RMT283, and RM6706, and to 8 ng p27 antigen for RM6715, RM6718, and RM6719. All CH505-infected RMs were inoculated intravenously with variable doses, corresponding to 10, 1, 0.1, and 0.01 ng p27 antigen for RM6697, RM6698, RM6699, and RM6701, respectively. RM6702 and RM6703 were initially inoculated with viral doses corresponding to 0.001 and 0.0001 ng p27 antigen, which resulted in undetectable plasma viral loads. Both animals were subsequently re-inoculated eight weeks later with a dose corresponding to 10 ng p27 antigen, with the time of re-inoculation designated as 0 WPI. Infections were monitored by determining circulating viral loads (vRNA/ml) in plasma samples taken at different intervals throughout the infection course. Lymph node biopsies were taken from infected RMs at various times: at 24, 48 and 72 week-post-infection (WPI) for RMs in the BG505-infected cohort, and at 24, 40 and 56 WPI for CH505-infected cohort.

Sample size was chosen based on prior studies demonstrating consistent viral kinetics [49] and shared Env mutation patterns across infected RMs [51]. Six animals per group were used for viral load and Env mutation ana ses, and three representative animals per group were analyzed in detail for B-cell responses due to the resource-intensive nature of these assays. Given that the animals are outbred and possess diverse immunoglobulin germline repertoires (Figure 7A), this sample size was considered sufficient to capture inter-individual variability while allowing detailed functional and genetic characterization of Env-reactive B-cell responses. The experimental unit for *in vivo* analyses was the individual animal, while *in vitro* and sequencing-based analyses were conducted at the level of B-cell cultures or clones as specified.

All animals that underwent SHIV infection and completed the planned sampling were included in the analyses. No predefined exclusion criteria were applied, and no animals or samples were excluded from the reported analyses. The primary outcomes were Env-reactive B-cell responses, including antigen binding, neutralization activity, and BCR sequence characteristics. Secondary outcomes included viral load kinetics, germinal center B-cell dynamics, and clonal expansion patterns. Neutralization assays were performed in a blinded manner, while other procedures were not blinded due to the study design. Statistical analyses were performed as described in the figure legends using two-sided tests.

### Plasma viral load quantification

Plasma viral load measurements were performed by the NIH/NIAID-sponsored Nonhuman Primate Virology Core Laboratory at the Duke Human Vaccine Institute. This core facility is CLIA certified and operates highly standardized, quality-controlled Applied Biosystems Real-Time SIV and HIV vRNA PCR assays. QIAsymphony SP and QIAgility automated platforms (QIAGEN) were used for high throughput sample processing and PCR setup. Viral RNA was extracted and purified from plasma, annealed to a target-specific primer, and reverse transcribed into cDNA. The cDNA was treated with RNase and added to a custom real-time PCR master mix containing target-specific primers and a fluorescently labeled hydrolysis probe. Thermal cycling was performed on a QuantStudio3 (ThermoFisher Scientific) real-time quantitative PCR (qPCR) instrument. Viral RNA copies/reaction was interpolated using quantification cycle data. Raw data were quality-controlled, positive and negative controls were checked, and the mean viral RNA copies/ml was calculated. The lower limit of quantification (LLOQ) of this assay was 62 RNA copies/ml.

### Viral single-genome sequencing

Single-genome sequencing of SHIV 3’ half genomes was performed as described [96]. For BG505-infected RMs, Env gene sequencing data at 88 WPI were included in the analysis. For CH505-infected RMs, sequencing data were included at 56 WPI for RM6697, RM6698, RM6699 and RM6701, at 24 WPI for RM6702, and at 48 WPI for RM6703. The Env gene nucleotide sequences were translated into amino acid sequences and filtered to retain only full-length sequences without premature stop codons. The filtered sequences were aligned with the original Env sequences used to produce the SHIVs for infection as references. Multiple sequence alignment of Env amino acid sequences was performed using the MUSCLE algorithm [97], as implemented in the *msa* R package, with log-expectation profile scoring, a gap-opening penalty of 3, and a maximum of 100 iterations. All other alignment parameters were retained at their default values, and the input sequence order was preserved. Amino acid numbering was standardized based on the HXB2 Env gene. Functional domains were assigned according to the corresponding domains in the HXB2 Env (UniProt entry P04578). Insertions at the same position were considered identical mutation events regardless of the inserted sequences and counted as single mutations regardless of their lengths.

### Cell culture media

B-cell culture medium (BCM) was formulated as RPMI-1640 medium (Gibco, Cat# 11875119) supplemented with 10% FBS (HyClone, Cat# SH30071.03), 10 mM HEPES buffer (Gibco, Cat# 15630080), 1 mM sodium pyruvate (GIBCO, Cat# 11360070), 1× MEM/NEAA (GIBCO, Cat# 11140050), 55 μM 2-mercaptoethanol (GIBCO, Cat# 21985023), 100 units/ml penicillin and 100 μg/ml streptomycin (GIBCO, Cat# 15140122). The culture medium for feeder cell lines (IMDM-10) was formulated as IMDM medium (GIBCO, Cat# 12440053) supplemented with 10% FBS (HyClone), 55 μM 2-mercaptoethanol, 100 units/ml penicillin and 100 μg/ml streptomycin (all Gibco). The culture medium for Expi293F cell line was formulated as Expi293 Expression Medium (GIBCO, Cat# A1435101) supplemented with 100 units/ml penicillin and 100 μg/ml streptomycin (Gibco).

### HIV Env antigens

Stabilized trimeric HIV-1 Env was produced as previously described [98, 99]. Plasmids encoding BG505.T332N DS-SOSIP.664 and chimeric CH505 T/F SOSIPv4.1 with a C-terminal AviTag was co-transfected with a furin expression plasmid into FreeStyle™ 293-F cells using 293fectin™ Transfection Reagent (ThermoFisher). Trimeric Env was purified by PGT145 affinity chromatography, biotinylated in a BirA enzymatic reaction (Avidity), purified by Superose™ 6 size exclusion chromatography. CH505 T/F gp120, CH505 T/F gp140C, and BG505.T332N gp140C antigens were expressed in FreeStyle™ 293-F cells using 293fectin™ (ThermoFisher) transient transfection. The gp140C design included a mutated furin cleavage site where amino acids REKR were mutated to REKE. gp140C and gp120 were purified by *Galanthis navalis* lectin chromatography and Superose™ 6 (gp140C proteins) or Superdex™ 200 (gp120 proteins) size exclusion chromatography as described previously [100]. MN.gp41 was purchased commercially (ImmunoDX, Cat# 1091).

### Antibodies

Fluorochrome-conjugated monoclonal antibodies used in this study for RM B cell phenotyping and sorting included: BB515-conjugated anti-human IgG (hIgG-BB515, 1/50 dilution, clone G18-145, BD, Cat# 564581), hCD38-PE (1/50 dilution, clone AT1, Santa Cruz, Cat# sc-7325-PE), hCD86-PE-D594 (1/50 dilution, clone IT2.2, BioLegend, Cat# 305434), hCD27-PE-Cy7 (1/100 dilution, clone M-T271, BD, Cat# 560609), hIgM-BV421 (1/50 dilution, clone G20-127, BD, Cat# 562618), hCD20-BV510 (1/50 dilution, clone 2H7, BioLegend, Cat# 302340), hCD21-BV711 (1/50 dilution, clone B-ly4, BD, Cat# 563163), hCD3-BV786 (1/50 dilution, clone SP34-2, BD, Cat# 563918), and 7-AAD (1/50 dilution, BD, Cat# 559925). Fluorochrome-conjugated monoclonal antibodies used for human B cell phenotyping and sorting included: hIgG-BB515 (1/50 dilution, clone G18-145, BD, Cat# 564581), hIgD-PE-Cy7 (1/50 dilution, clone IA6-2, BioLegend, Cat# 348210), CD19-APC-F750 (1/50 dilution, clone HIB19, BioLegend, Cat# 302258), CD27-BV421 (1/50 dilution, clone M-T271, BioLegend, Cat# 356418), IgM-BV510 (1/50 dilution, clone G20-127, BD, Cat# 563113), CD3-BV650 (1/50 dilution, clone UCHT1, BioLegend, Cat# 300468), CD14-BV650 (1/50 dilution, clone M5E2, BioLegend, Cat# 301836), CD16-BV650 (1/50 dilution, clone 3G8, BioLegend, Cat# 302042), CD24-BV711 (1/50 dilution, clone ML5, BioLegend, Cat# 311136), and 7-AAD (1/50 dilution, BD, Cat# 559925).

For rhesus IgG ELISA assays, the coating antibody used was mouse anti-monkey IgG (2 µg/ml, clone 5C12.D4, ThermoFisher, Cat# SA1-10653), the standard antibody used was rhesus monkey IgG (polyclonal, Southern Biotech, Cat# 0135-01), and the detection antibody used was mouse anti-monkey IgG-HRP (1/2000 dilution, clone SB108a, Southern Biotech, Cat# 4700-05). For human IgG ELISA assays, the coating antibodies used were goat anti-human Igκ (2 µg/ml, polyclonal, Southern Biotech, Cat# 2061-01) and goat anti-human Igλ (2 µg/ml, polyclonal, Southern Biotech, Cat# 2071-01), the standard antibodies used were human IgG1 Kappa (polyclonal, Southern Biotech, Cat# 0151K-01) and human IgG1 Lambda (polyclonal, Southern Biotech, Cat# 0151L-01), and the detection antibody used was goat anti-human IgG-HRP (1/5000 dilution, polyclonal, Southern Biotech, Cat# 2040-05).

For rhesus IgG Luminex assays, the capture antibody conjugated with Luminex microspheres was the same as the coating antibody used for ELISA, and the detection antibody used was mouse anti-monkey IgG-PE (2 µg/ml, clone SB108a, Southern Biotech, Cat# 4700-09). For human IgG Luminex assays, the capture antibody conjugated with Luminex microspheres was goat anti-human IgG (Jackson ImmunoResearch, Cat# 109-005-098), and the detection antibody used was goat anti-human IgG-PE (2 µg/ml, polyclonal, Southern Biotech, Cat# 2040-09). The same standard antibodies as those for ELISA assays were used.

V(D)J sequences of reference antibodies were synthesized (ThermoFisher or Gene Universal) and cloned in to human IgG1, Igκ and Igλ expression vectors, including VRC01 [69], CH31 [101], 3BNC117 [102], PGV04 [103], 2G12 [104], 10-1074 [105], PGT121, PGT135 [106], PG9, PG16 [107], PGT145 [106], CH01 [101], PGT151 [108], VRC34.01 [109], 35O22 [110], 3791 [82] and 447-52D [111], 17b [112], A32 [113], 412d [114] and F105 [115]. 14e and 19b [116] expression plasmids were kindly provided by Dr. James E. Robinson (Tulane University).

### Flow cytometry phenotyping and sorting

Frozen RM LN cells or human PBMCs were thawed in pre-warmed BCM (see cell culture media above), centrifuge at 300× *g* for 2 min, resuspended in BCM, and labeled with fluorochrome-conjugated antibody cocktails (see antibody list above). After incubation with antibodies at 4°C in the dark for 30 min, cells were washed, resuspended in BCM and stored on ice until flow cytometry detection. For RM LN cells, four major B-cell populations were gated for single-cell sorting, including IgG^+^ GC B cells (7-AAD^−^ CD3^−^ CD20^+^ IgG^+^ CD38^−^), IgG^+^ Bmem cells (7-AAD^−^ CD3^−^ CD20^+^ IgG^+^ CD38^+^ CD86^−^), IgM^+^ Bmem cells (7-AAD^−^ CD3^−^ CD20^+^ IgM^+^ CD38^+^ CD86^−^ CD27^+^ CD21^hi^) and MF B cells (7-AAD^−^ CD3^−^ CD20^+^ IgM^+^ CD38^+^ CD86^−^ CD27^−^ CD21^int^). For BCR Rep-Seq, MF B cells were sorted in bulk directly into TRIzol LS Reagent (ThermoFisher, Cat# 10296010) for RNA extraction. For human PBMCs, three B-cell populations were gated for single-cell sorting, including naïve B cells (7-AAD^−^ CD3^−^ CD14^−^ CD16^−^ CD19^+^ CD27^−^ CD24^int^ IgD^+^ IgM^+^ IgG^−^), IgM^+^ Bmem cells (7-AAD^−^ CD3^−^ CD14^−^ CD16^−^ CD19^+^ CD27^+^ CD24^hi^ IgD^+^ IgM^+^ IgG^−^), and IgG^+^ Bmem cells (7-AAD^−^ CD3^−^ CD14^−^ CD16^−^ CD19^+^ CD27^+^ CD24^hi^ IgD^−^ IgM^−^ IgG^+^). Flow cytometry phenotyping and sorting was performed with BD Aria II or Symphony S6 at the Duke Human Vaccine Institute Research Flow Cytometry Facility.

### Generation of feeder cell line MEC-147

The MS40L-low feeder cell line [59] was transduced with lentiviral vectors encoding human IL-2, IL-4, IL-21, and BAFF, along with EGFP and a puromycin-resistance gene as selection markers. Following puromycin (2 µg/ml, Sigma, Cat# P8833) selection, clonal cell lines were established via single-cell sorting based on EGFP and cell surface CD40L expression. These clonal feeder cell lines were then functionally tested and compared in single B-cell cultures. Among them, one cell line, MEC-147, was selected for its high efficiency in supporting single B-cell cultures across all RM and human B-cell populations tested (Figures S1B and S1C).

### Single B-cell culture

MS40L-low or MEC-147 feeder cells were maintained in IMDM-10 medium (see cell culture media above). Twenty-four hours before single B-cell sorting, the feeder cells were harvested by Trypsin-EDTA (0.05%, ThermoFisher, Cat# 15400054) treatment, resuspended in pre-warmed BCM, and seeded into 96-well flat-bottom culture plates at 3000 cells per 100 µl for each well. Prior to single B-cell sorting, 100 µl of additional BCM was added to each well of the feeder cell plates. For RM B-cell cultures with the MS40L-low feeder cell line, the additional BCM was supplemented with 10% heat-inactivated human serum (Gemini, Cat# 100-512) and a 2× concentration of a cytokine cocktail. The 10,000× stock of the cocktail contained 500 µg/ml recombinant human IL-2 (Peprotech, Cat# 200-02), 100 µg/ml recombinant human IL-4 (Peprotech, Cat# 200-04), 100 µg/ml recombinant human IL-21 (Peprotech, Cat# 200-21), and 100 µg/ml recombinant human BAFF (Peprotech, Cat# 310-13). For human B-cell cultures with the MS40L-low feeder cell line, the additional BCM was supplemented with a 2× concentration of the cytokine cocktail. In contrast, for RM or human B-cell cultures with the MEC-147 feeder cell line, no supplements were added to the additional BCM.

B cells from different populations were sorted into feeder cell plates at a density of one cell per well using flow cytometry to initiate single B-cell cultures. The medium was changed every 3∼4 days. Supernatants were collected for IgG ELISA assays on day 18 for RM B-cell cultures and on day 25 for human B-cell cultures. Following supernatant collection, the culture plates with clonally expanded B cells were frozen and stored at –80°C for subsequent RNA extraction and V(D)J sequencing.

### Screening of IgG-producing clones by ELISA assays

Standard sandwich ELISA assays were conducted to screen IgG-producing B-cell clones using single B-cell culture supernatants. Briefly, 384-well high-binding ELISA plates (Corning, Cat# 3700) were coated overnight at 4°C with coating antibodies (listed above) diluted in carbonate buffer (100 mM NaHCO_3_ and 2.5 mM Na_2_CO_3_). The plates were washed three times with washing buffer (PBS with 0.1% Tween-20) using a microplate washer (BioTek ELx405), then blocked for 30 minutes with blocking buffer (PBS with 0.5% BSA)). After removing the blocking buffer, diluted B-cell culture supernatants were added. RM B-cell culture supernatants were diluted 1:10, and human B-cell culture supernatants were diluted 1:100 in dilution buffer (PBS with 0.5% BSA and 0.1% Tween-20). The plates were incubated for 2 hours at room temperature or overnight at 4°C. After three washes, detection antibodies were added and incubated at room temperature for 1 hour in the dark. Following another three washes, TMB substrate (BioLegend, Cat# 421101) was added and incubated at room temperature in the dark for 10 minutes for RM samples and 5 minutes for human samples. Stop solution (1 M H_2_SO_4_) was then added, and optical density (OD) values were measured at 450 nm with background subtraction at 650 nm. Supernatants from culture wells without sorted B cells served as negative controls. Culture supernatants with OD values exceeding the mean plus six times the standard deviation (mean + 6 × SD) of negative control OD values were considered IgG-positive.

### TZM-bl neutralization assays

IgG-positive culture supernatants identified through ELISA screening were transferred to master plates. One set of supernatant aliquots from the master plates was used for standard TZM-bl neutralization assays [63, 64] against pseudo-viruses expressing autologous Env antigens. Briefly, 90 µl of culture supernatant was added to individual wells of 96-well culture plates. A pre-titrated dose of virus (45 µl) was added, mixed, and incubated for 1 hour at 37°C. Freshly trypsinized TZM-bl cells (110 µl) containing an optimized concentration of DEAE-dextran in growth medium were added to each well followed by incubation for 48 hours at 37°C. Infection was quantified as a function of Tat-regulated firefly luciferase activity [63]. Reductions in relative luminescence units (RLUs) were determined relative to RLU in virus control wells (cells + virus in the absence of test sample) after subtraction of background RLU in cell control wells (cells only). Env-pseudotyped viruses were prepared by transfection in 293T/17 cells as described [63].

The neutralization breadth of rAbs produced based on identified Env-reactive BCR sequences was assessed using a similar approach. A panel of twelve tier 2 Env-pseudotyped viruses was tested individually (Table S4), including pseudo-viruses bearing the SHIV Envs, BG505 and CH505, used for the initial infections. Additional pseudo-viruses expressed the Envs of ZM233 (clade C), and nine heterologous strains from the global panel of reference strains [73]. The global panel Envs included X1632 (clade G), 25710 (clade C), 246F3 (clade AC), BJOX2000 (clade CRF07), CE1176 (clade C), CE0217 (clade C), CH119 (clade CRF07), CNE55 (clade CRF01), and TRO11 (clade B) [73]. SVA-MLV, a pseudotyped negative control virus incorporating murine leukemia virus envelope, was included to confirm assay specificity.

### Luminex® binding assays

Carboxylated MicroPlex® microspheres (Luminex Corp.) were activated using Sulfo-NHS (ThermoFisher, Cat# 24510) and EDC (ThermoFisher, Cat# 22980) and subsequently coupled with protein antigens. A panel of HIV Env antigens along with non-relevant control antigens were tested in this study, including BG505.SOSIP, BG505.gp140, CH505.SOSIP, CH505.gp140, CH505.gp120, MN.gp41, streptavidin, BSA, and ovalbumin. For Avi-tagged biotinylated SOSIP and gp140 antigens, the microspheres were first coupled with streptavidin to enable indirect conjugation with the antigens. Additionally, a subset of microspheres was coupled with an anti-monkey IgG capture antibody (ThermoFisher, Cat# SA1-10653) for detecting rhesus IgGs.

In a blinded manner, alongside the neutralization screening assays described above, another set of single B-cell culture supernatant aliquots from the IgG-positive master plates was analyzed using Luminex binding assays against autologous SOSIP and gp140, as well as control antigens. Briefly, the supernatants were diluted 1:10 in Luminex dilution buffer (PBS with 1% BSA, 1% non-fat milk, 0.05% Tween-20, and 0.05% NaN_3_) and incubated with the multiplexed microsphere panel in MultiScreen filter plates (Millipore, Cat# MSBVN1250) under gentle shaking (70 rpm) at room temperature in the dark for two hours. Following three washes using a vacuum manifold with Luminex washing buffer (PBS with 1% BSA, 0.05% Tween-20, and 0.05% NaN_3_), detection antibodies were added and incubated under the same conditions for one hour. After three additional washes, the microspheres were resuspended in washing buffer by vortexing (800 rpm for 30 seconds) and stored at 4°C in the dark until analysis on a Bio-Plex 3D Suspension Array System (Bio-Rad).

For rAbs, including reference bnAbs and nnAbs in the form of human IgG1, as well as identified Env-reactive antibodies in the form of rhesus IgG1, testing was conducted using serial dilutions. The Luminex antigen panel included SOSIP and gp140 antigens from both BG505 and CH505 strains. For competition binding assays, multiplexed microspheres coupled with autologous SOSIP and gp140 antigens were first incubated with competitor human IgG1 rAbs for 30 minutes. Subsequently, the analyte antibodies (Env-reactive rhesus IgG1 rAbs) were added. The final concentrations for competitor and analyte antibodies were 25 and 1 µg/ml, respectively.

### Competition binding assay data analysis

Binding activities of produced rhesus IgG1 rAbs (analytes) to autologous SOSIP and gp140 (antigens) were measured in the presence of various human IgG1 bnAbs and nnAbs (competitors) of known specificities. The percentages of inhibition were calculated for each competitor antibody relative to the baseline MFI values in the absence of competitors (Table S5). Based on the percentages of inhibition data across all antigen-analyte-competitor combinations tested, a unified UMAP embedding was established. Clustering was applied to the 2D UMAP coordinates using the Density-Based Spatial Clustering of Applications with Noise (DBSCAN) algorithm [117] implemented in the *dbscan* R package. A minimum of 2 data points (*minPts = 2*) was allowed during clustering. To select the neighborhood radius parameter *epsilon* (*eps*), we first computed the distance to each point’s second-nearest neighbor (*k = 2*) and plotted the sorted distance values to generate a *k*-nearest neighbor distance curve. We then identified the point of steepest increase in the curve. The midpoint of this maximal slope segment was used as the *eps* value, providing a data-driven threshold that balances cluster sensitivity with robustness to noise. To assess whether clustering patterns were driven by batch effects, we color-coded the UMAP plots based on assay batches and found no apparent clustering by batch (Figure S3D).

### RNA extraction for BCR V(D)J sequencing and Rep-Seq

For single B-cell culture supernatants demonstrating neutralization or Env-binding activities, RNA was extracted from frozen cell pellets stored in corresponding original culture wells. High-throughput RNA extraction was conducted using the Quick-RNA 96 Kit (Zymo Research, Cat# R1053). For Rep-Seq, RNA was extracted from MF B cells that were directly sorted into TRIzol LS Reagent (ThermoFisher) using the Direct-zol RNA Miniprep Kit (Zymo Research, Cat# R2052).

### A versatile and unbiased BCR V(D)J sequencing platform

Since the IGV gene repertoires are highly diverse in RMs and have not been completely characterized [65, 66], a new BCR V(D)J sequencing platform was developed, including a method for unbiased amplification of full-length V(D)J sequences using the SMART (switching mechanism at the 5’ end of the RNA transcript) technology [118], a method using PacBio Sequel technology for high-throughput long-read-length sequencing, and a user-friendly computer program, *AbSolute*, for semi-automatic data analysis. This versatile platform supports human, rhesus and mouse BCR V(D)J sequencing and data analysis in its current version, and can be easily extended to other mammalian species as long as the Ig constant region sequences are known.

RNA samples extracted from cultured clonal B cells were reverse-transcribed into full-length V(D)J sequences using the SMART technology for unbiased amplification. Briefly, RNA samples were mixed with dNTPs (NEB, Cat# N0447L), a template-switching oligo (TSO, with 5’ padding and one of the unique plate-barcoding sequences), and reverse primers specific to the constant regions of IgG, Igκ, Igλ, and optionally IgM (see Table S7 for primer sequences, with “Purpose” column as “BCR V(D)J sequencing”). The mixture was denatured at 72°C for 3 minutes (min) and annealed on ice for 2 min. A reverse transcription (RT) pre-mixture was then added, comprising 1/5 final volume of 5× RT buffer, 1/10 final volume of 20 mM DTT, 1/20 final volume of RNaseOUT™ RNase Inhibitor (ThermoFisher, Cat# 10777-019), and 1/20 final volume of SMARTScribe™ Reverse Transcriptase (TaKaRa Bio, Cat# 639538). The RT reaction was carried out on a thermocycler at 42°C for 90 min, followed by 70°C for 15 min. Final concentrations were 0.5 mM for each dNTP, 1.0 µM for the TSO and 0.2 µM for each reverse primer.

The full-length V(D)J sequences were amplified using two rounds of semi-nested PCRs. In PCR1, a single common forward primer specific to the TSO was used alongside one of the reverse primers targeting the constant regions of IgG, IgK, IgL, or optionally IgM (located upstream of the corresponding RT primers; see Table S7 for primer sequences) in separate reactions. The forward primers for PCR2 comprise a common set of twelve primers specific for the common PCR1 forward priming site, along with a 5’ padding and one of the twelve column-barcoding sequences. Each set of reverse primers for PCR2 is specific for the constant regions of one of the Ig heavy and light chain classes (IgG, IgK, IgL, or optionally IgM) at regions further upstream of the corresponding PCR1 reverse primers. Each set comprises eight primers, with 5’ padding and one of the eight row-barcoding sequences. Together, each combination of forward and reverse primer sets created a 12 × 8 matrix for barcoding a 96-well plate of reactions. The reaction systems for both PCRs included 1× Herculase Reaction Buffer, 0.2 mM of each dNTP, 0.2 µM of each primer, 1/10 final volume of template DNA, and 1/50 final volume of Herculase II Fusion DNA Polymerase (Agilent, Cat# 600679). cDNA from the RT reactions served as templates for PCR1, while PCR1 products were used as templates for PCR2. For the rhesus system, a common cycling condition was applied to both PCRs, as well as to PCR1 for the human system: 95°C for 4 min; 30 cycles of 95°C for 30 seconds (sec), 55°C for 20 sec, and 72°C for 30 sec; followed by 72°C for 10 min. PCR2 for the human system differed slightly, with the annealing temperature increased from 55°C to 59°C.

The individually barcoded PCR2 amplicons were pooled into a single tube, combining IgG, IgK, IgL, and optionally IgM amplicons derived from one original 96-well plate of RNA samples. The pooled amplicons were gel-purified, ethanol-precipitated, and eluted in Tris-Cl (10 mM, pH 8.0). The purified amplicon libraries were submitted for quality check, further pooling, PacBio SMRTbell® library preparation, and sequencing on the PacBio Sequel platform (DNA Link Inc., Korea).

For the processing of circular consensus sequencing (CCS) reads derived from PacBio sequencing, a custom computer program, *AbSolute*, was developed using C# language. Bulk CCS reads from multiple original RNA plates were demultiplexed into individual plates based on plate barcoding sequences. The CCS reads for individual plates were further demultiplexed into individual wells and Ig heavy and light chain classes based on the combination of column and row barcoding and class-specific constant region sequences. Multiple sequence alignment for each Ig heavy and light chain class in each well was performed programmatically using the MUSCLE [97] and Kalign [119] algorithms, and a consensus sequence was determined as the final output of V(D)J sequences. The *AbSolute* program features a graphical user interface and a semi-automatic analysis workflow. Default settings for analyzing human, rhesus, and mouse antibody genes are embedded. A user-defined settings option allows for the easy extension of the analysis to other mammalian species or even different genes.

### Development of multiplex primer sets targeting IGV 5’UTR and leader regions

A strategy for establishing individualized RM IGHV germline databases was developed in a recent study [66]. First, Rep-Seq is performed using 5’RACE (rapid amplification of cDNA ends) libraries prepared based on SMART technology [118] to obtain full-length 5’ UTR and leader region sequences of IGHV repertoires from multiple RMs. Multiplex primer sets are designed to target these regions with broad repertoire coverage. Rep-Seq is then conducted for individual RMs using these multiplex primer sets to generate full-length VDJ sequences, which are used to establish individualized IGHV germline databases with the *IgDiscover* program [75]. We employed and extended this strategy to establish individualized IGV germline databases for both heavy and light chains. First, building on published protocols [120] and as detailed below, we developed multiplex primer sets for rhesus IGKV and IGLV segment families and updated the IGHV primer sets from the initial study [66].

From eight outbred RMs (including the six SHIV-infected RMs involved in this study, a seventh RM (RM6702) infected with SHIV CH505, and an eighth RM (A6E030) immunized with DTaP), approximately 1 × 10^5^ MF B cells were sorted from each RM via flow cytometry for RNA extraction. RT reactions using SMART technology were performed as follows:

1) A total of 20∼100 ng input RNA in 8.0 µl was mixed with 1.0 µl dNTPs (10 mM each) and 1.0 µl of 50 µM Oligo(dT)_20_ (ThermoFisher, Cat# 18418020). The mixture was denatured at 72°C for 3 min, then annealed on ice for 2 min.
2) To the reaction vial, a mixture of 3.5 µl 5× RT Buffer, 2.0 µl 20 mM DTT, 1.0 µl RNaseOUT RNase Inhibitor (ThermoFisher), and 1.0 µl SMARTScribe™ Reverse Transcriptase (TaKaRa Bio) was added. The reaction was incubated at 42°C for 1 hour on a thermocycler.
3) While maintaining 42°C on the thermocycler, the reaction vial was supplemented with a mixture containing 2.0 µl of 10 µM TSO (Read1_TS, primer sequences in Table S7), 0.5 µl 5× RT Buffer, and 0.5 µl SMARTScribe™ Reverse Transcriptase. The reaction was further incubated at 42°C for 90 min, followed by 70°C for 15 min.

Column-purified cDNA was used as the template for unbiased amplification of full-length V(D)J sequences through two rounds of semi-nested PCRs using KAPA HiFi HotStart ReadyMix DNA Polymerase (Kapa Biosystems, Cat# KK2602). A common forward primer (Read1U), targeting the Read1 priming sequences on the TSO, was used in both PCRs at a final concentration of 0.25 µM. Reverse primers specific to the constant regions of rhesus IgM, IgK, or IgL (see Table S7 for primer sequences, with “Purpose” column as “Rep-Seq, 5’RACE”) were included in separate reactions at a final concentration of 0.5 µM. The cycling conditions were as follows: 96°C for 5 min; 20 cycles for PCR1 or 10 cycles for PCR2 of 95°C for 20 sec, 69°C for 20 sec, and 72°C for 20 sec; followed by 72°C for 5 min. The PCR1 products were column-purified and used as templates for PCR2. PCR2 products were gel-purified, eluted in Tris-Cl, and quantified using the Qubit 1× dsDNA HS Assay Kit (ThermoFisher, Cat# Q33230).

Index PCR was performed using KAPA HiFi HotStart ReadyMix DNA Polymerase with 10 ng of purified PCR2 products in a final volume of 50 µl, along with Illumina dual-index primers (NEB, Cat# E7600S) at final concentrations of 0.4 µM each. The cycling conditions were 96°C for 5 min; 10 cycles of 95°C for 30 sec, 68°C for 30 sec, and 72°C for 30 sec; followed by 72°C for 10 min. The index PCR products were purified using AMPure XP beads (Beckman Coulter, Cat# A63881), eluted in Tris-Cl, and submitted for quality check and pooling. The pooled libraries, along with 15% PhiX174 DNA, were sequenced using MiSeq v3 300PE flow cells at the Duke University Sequencing and Genomic Technologies Shared Resource.

The sequencing data from the 5’RACE libraries were analyzed using the *IgDiscover* program [75]. IMGT® [121] rhesus IGV, IGHD, and IGJ allele collections served as the starter databases for V(D)J inference. The pseudo-CDR3 locations for IGH, IGK, and IGL were defined as [-125, -89], [-97, -70], and [-110, -80], respectively. Consensus UTR and leader sequences were extracted using the “*upstream*” subcommand, based on the assigned reads from the final iteration of V(D)J inference. Degenerate primer sets were designed (see Table S7 for primer sequences, with “Purpose” column as “Rep-Seq, MTPX”) to target the UTR and leader regions, covering the consensus sequence collections derived from the eight RMs included in the 5’RACE library sequencing.

### Establishment of individualized IGV germline databases

Using the multiplex primer sets developed above to target IGV 5’UTR and leader regions, MTPX libraries were prepared for BCR Rep-Seq and establishment of individualized IGV germline databases, building on published protocols [120] and as detailed below.

The same RNA samples used for preparing 5’RACE libraries were utilized for MTPX library preparation. RT reactions were performed as follows:

1) A total of 20∼100 ng input RNA in 10.0 µl was mixed with 2.0 µl dNTPs (5 mM each) and 1.0 µl each of 10 µM reverse primers targeting rhesus IgM, IgK, and IgL constant regions (see Table S7 for primer sequences, with the “Purpose” column as “Rep-Seq, MTPX”). The mixture was denatured at 65°C for 5 min, annealed at 37°C for 5 min, and placed on ice for 2 min.
2) To the reaction vial, a mixture of 1.75 µl DNase/RNase-free H_2_O, 2.0 µl 10× RT Buffer, 0.25 µl RiboLock RNase Inhibitor (ThermoFisher, Cat# EO0381), and 1.0 µl Sensiscript RTase (QIAGEN, Cat# 205211) was added. The reaction was incubated at 37°C for 1 hour, followed by 72°C for 10 min.

The cDNA samples were amplified by PCR using a common reverse primer, Read2U, and one of the multiplex primer sets (see Table S7 for primer sequences) in individual reactions with KAPA HiFi HotStart PCR Kit (Kapa Biosystems, Cat# KK2502). The reactions contained dNTPs at 0.3 mM each, Read2U at 0.5 µM, and multiplex primers at 0.1 µM each. The cycling conditions were as follows: 96°C for 3 min; 25 cycles of 95°C for 20 sec, 68°C for 20 sec, and 72°C for 20 sec; followed by 72°C for 5 min. The PCR products were gel-purified, eluted in Tris-Cl, and quantified using the Qubit 1× dsDNA HS Assay Kit. Index PCR was performed using KAPA HiFi HotStart PCR Kit with 10 ng of purified PCR products in a final volume of 50 µl, along with Illumina single-index primers (NEB, Cat# E7335 and E7500) at final concentrations of 0.4 µM each. Library preparation and MiSeq sequencing were performed in the same way as for the 5’RACE libraries.

The sequencing data from the MTPX libraries were analyzed for germline V(D)J inference using the *IgDiscover* program [75], employing the same starter databases as used for the 5’RACE libraries. The pseudo-CDR3 locations for IGH, IGK, and IGL were defined as [-161, -126], [-119, -93], and [-131, -101], respectively. Filtered and assigned reads from the two libraries prepared using 5’UTR and leader primer sets were merged. Inferred germline IGV alleles differing only in terminal sequence lengths were deduplicated, retaining only the longest germline sequences. Individualized IGV germline databases were then established based on these unique germline sequences for each RM. Unique CDR3 counts were determined for each unique inferred germline IGV allele, formulating the germline IGV allele repertoire used for the analysis in Figure 7C.

### V(D)J assignment and clonal analyses

V(D)J assignment for both heavy and light chain sequences determined from cultured clonal B cells was performed using the *IgBLAST* program [122], based on the individualized IGV germline databases generated for each RM. Clonal relationships were inferred based on *IgBLAST*-assigned heavy chain data using the *scoper* program [123] with a hierarchical clustering-based model [124] and an arbitrary threshold of 0.2. For heavy chains, when more than one productive VDJ sequence was recovered from a single B-cell culture well, we retained the sequence with the largest clonal member count and the highest IGHV identity. For light chains, when more than one VJ sequence was recovered from a single B-cell culture well, sequences were first filtered for unique IGLV and IGLJ usage and for the highest IGLV identity. Among these filtered candidates, if any shared identical IGLV and IGLJ segments with clonal member(s) inferred from the heavy chain sequences, that single light-chain sequence was retained. For the heavy chains, germline sequences were inferred with the *Change-O* toolkit [125] using the *CreateGermlines.py* program based on individualized IGV germline databases, and phylogenetic trees were reconstructed with programs *BuildTrees.py* and *IgPhyML* [126]. The filtered heavy chain clonal data and light chain *IgBLAST*-assigned data were combined with time-course, phenotypic, and functional data (Table S3). Incorporating these combined data, the phylogenetic trees of identified Env-reactive B-cell lineages were visualized using the R package *ggtree* [127].

### Functional typing of Env-reactive B-cell clones

Each individual Env-reactive B cell was first assigned with a functional code based on binding MFI values to autologous SOSIP and gp140 proteins and autologous neutralization activities measured with corresponding single B-cell culture supernatants. To facilitate downstream calculation, SOSIP binding MFI values below 2 were offset to 2. The functional code was assigned according to the following successive cases:

1) Code 1: (SOSIP MFI < 3000) or (SOSIP MFI < gp140 MFI × 0.5)
2) Code 4: (SOSIP MFI ≥ 3000) and (SOSIP MFI ≥ gp140 MFI × 3) and (Neutralization < 50%)
3) Code 5: (SOSIP MFI ≥ 3000) and (SOSIP MFI ≥ gp140 MFI × 3) and (Neutralization ≥ 50%)
4) Code 2: Neutralization < 50%
5) Code 3: all the rest cases

Env-reactive B cell clones, defined by heavy-chain clone ID within an animal across all timepoints, were then assigned a clonal type based on the distribution of functional codes among their members. For each clone, the maximum code observed (maxType) was determined. The threshold for strong binding was defined as half of the maximum SOSIP MFI within the clone (capped at 10,000 MFI), and codes from all members exceeding this threshold (strongTypes) were collected. The most frequent strong code (mostType) was then identified; in case of a tie, the higher code value was chosen.

Final clone types were assigned by cross-referencing (maxType, mostType) according to the following mapping: (maxType = 1, mostType = 1) as Type D, (maxType = 2, mostType = 1) as Type D, (maxType = 2, mostType = 2) as Type C, (maxType = 3, mostType = 1) as Type D, (maxType = 3, mostType = 2) as Type B, (maxType = 3, mostType = 3) as Type B, (maxType = 4, mostType = 1) as Type D, (maxType = 4, mostType = 2) as Type C, (maxType = 4, mostType = 3) as Type B, (maxType = 4, mostType = 4) as Type A, (maxType = 5, mostType = 1) as Type D, (maxType = 5, mostType = 2) as Type B, (maxType = 5, mostType = 3) as Type B, (maxType = 5, mostType = 4) as Type A, and (maxType = 5, mostType = 5) as Type A. An exception was made for clones with maxType = 4 and mostType = 2, if their strong binders also included code 3, they were reassigned to Type B. This rule-based procedure ensured that each Env-reactive B cell clone was consistently classified into one of four categories (Types A–D), integrating both the strongest activity observed and the predominant behavior of strong-binding members.

### V(D)J cloning and rAb production

V(D)J sequences of selected Env-reactive BCRs were cloned into mammalian expression vectors containing rhesus IgG1, IgK, or IgL constant region sequences using standard molecular cloning procedures. Endotoxin-free plasmids were prepared with the E.Z.N.A.® Endo-free Plasmid DNA Mini Kit II (Omega Bio-tek, Cat# D6950).

For antibody production, Expi293F™ cells (ThermoFisher, Cat# A14527) were co-transfected with paired heavy and light chain expression plasmids using the ExpiFectamine™ 293 Transfection Kit (ThermoFisher, Cat# A14524). Transfected cells were cultured in Expi293™ Expression Medium (ThermoFisher) with shaking at 150 rpm for 72∼96 hours. rAbs in the form of human or rhesus IgG1 were purified from the transfection supernatants using Pierce™ Protein G UltraLink™ Resin (Thermo Scientific, Cat# 53126) and eluted with IgG Elution Buffer (ThermoFisher, Cat# 21004). The eluted rAbs were buffer-exchanged into PBS using Amicon® Ultra-4 Centrifugal Filter Units (Millipore, Cat# UFC805096), sterilized by filtration through Millex-GV Syringe Filters (Millipore, Cat# SLGV013SL), and quantified using a NanoDrop 2000 (ThermoFisher).

### Quantification of variation in IGV allele usage among RMs

IGV allele usage frequencies were determined for each RM, and variation among three RMs in each infection group was quantified using two complementary approaches: (1) allele-wise dispersion, measured by the mean absolute deviation from the median (MADM), and (2) repertoire-level similarity, measured by cosine similarity. To control for differences in sampling depth between MF BCR repertoires and Env-reactive BCRs, MF repertoires were down-sampled using a multinomial distribution parameterized by allele frequency probabilities obtained from Rep-Seq data. For each RM, the multinomial sample size was matched to the number of Env-reactive BCRs detected. Allele usage frequencies were then recalculated from the down-sampled repertoires. This random sampling was repeated 1000 times to generate null distributions of allele-wise MADM and repertoire-level cosine similarity. Observed values from Env-reactive BCRs were compared against these null distributions, and empirical *p* values were derived to assess significance of differences between Env-reactive and MF repertoires. The same analyses were also applied to allele frequency data after smoothing (see below) within IGV genes or families.

Specifically, the allele usage frequencies for MF BCR repertoire were calculated based on unique CDR3 calls from BCR Rep-Seq data. MADM was calculated as

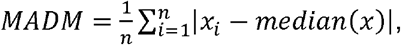

where *x* is the set of allele usage frequencies for a given allele across three RMs in an infection group, and *x_i_* is the allele frequency from the -th RM. Cosine similarity was calculated between every two RMs as a pair. Let 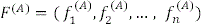 and 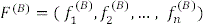 be the allele frequency vectors of two RMs over *n* IGV alleles for a given IG segment family (IGH, IGK or IGL). The cosine similarity between these vectors is calculated as

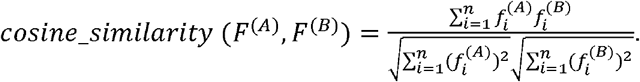

The multinomial down-sampling was performed as follows. Let 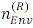 be the number of Env-reactive BCRs detected for RM *R*, and let 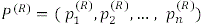 be the observed allele frequencies in the MF repertoire of the same RM, where is the total number of unique IGV alleles identified from the three RMs in each infection group and 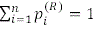. The down-sampled counts of MF BCRs for each allele, 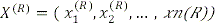 are drawn from a multinomial distribution: 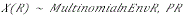. The corresponding allele frequencies after down-sampling are then defined as 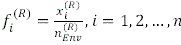.

For the MADM distribution, empirical *p* values were calculated as the proportion of simulated median MADM values greater than or equal to the observed median, with a “+1” correction applied to both numerator and denominator to avoid zero probabilities. Specifically, we first pooled MADM values from 1000 multinomial down-sampling simulations of MF repertoire. From this pooled distribution, we randomly sampled sets of MADM values (matched in size to the Env-reactive dataset) 1000 times to obtain simulated median MADMs from each sampling. Empirical *p* values were then calculated as

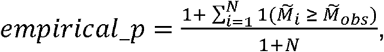

where 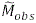 is the observed median MADM of Env-reactive BCRs, 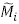 is the simulated median MADM from the *i*-th random sampling, and *N* = 1000. Alleles with zero frequencies across all three RMs were excluded for both observed and simulated datasets to prevent distortion by an excess of MADM = 0 values.

For the cosine similarity analysis, empirical *p* values were calculated as the proportion of simulated cosine similarity values less than or equal to the observed value, again with a “+1” correction. For each pair of RMs within an infection group and for each IGV segment family, cosine similarity was calculated from Env-reactive BCRs (observed) and from 1000 down-sampled MF repertoires (null). Empirical p values were then obtained as

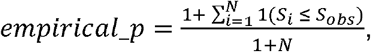

where *S_obs_* is the observed cosine similarity of Env-reactive BCRs, *S_i_* is the simulated cosine similarity from the *i*-th down-sampling, and *N* = 1000. Unlike MADM, alleles with zero frequencies across all three RMs were retained for cosine similarity analysis.

### Smoothing of the allele frequencies among IGV alleles

For an IGV repertoire across three RMs in one infection group, the allele frequencies (percentage among reads with unique CDR3 sequences) in a given RM for each individual IGV allele were redistributed among all the alleles within the threshold distance, weighted based on their pairwise distances. Specifically, let *N* be the number of unique IGV alleles in the repertoire, *x_i_* the detected frequency of the *i*-th IGV allele, and D = [*d_ij_*] the *N* × *N* pairwise distance matrix, where *d_ij_* represents the allelic sequence similarity distance between IGV allele *i* and *j*. Let *τ*, be the threshold distance that best discriminates IGV genes (Figure S4B) or families (Figure S4C), and let – be the minimal smoothing weight at the threshold distance. We define a weight matrix *W* = [*W_ij_*], where

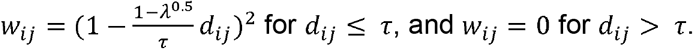

In our analysis, we set *λ* = 0.5 for smoothing within IGV genes and *λ* = 0.0001 for smoothing within IGV families. To normalize the weights so that they sum to 1 for each IGV allele *i*, we define

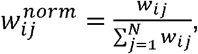

where 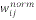 represents the relative contribution of allele *j* to the smoothed frequency of allele *i*. The smoothed frequency 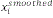 of each allele is then computed as

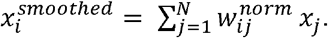

Collectively, this process redistributes the detected frequencies among closely related IGV alleles based on their genetic proximity.

### HCDR3 length and joint repertoire feature analysis

MF BCR repertoire sequences generated using leader– and 5’UTR-targeting primers were pooled for each RM. Records with missing or empty HCDR3 sequences were excluded. Sequences were deduplicated according to the combination of assigned IGHV, assigned IGHJ, and HCDR3 nucleotide sequence to generate the uCDR3 dataset (Table S8). Env-reactive BCRs (Table S3) were grouped by heavy-chain clonotype within each RM, and each clonotype was represented once in the analysis (xClonotype). For clonotypes containing multiple sequences, the most frequent non-missing value was retained separately for IGHV assignment, IGHJ assignment, HCDR3 length, and functional type (Table S8). HCDR3 length distributions were expressed as percentages of the corresponding uCDR3 or xClonotype repertoire (Figure S5A).

Differences between HCDR3 length distributions were quantified using Wasserstein-1 distance, calculated as the integral of the absolute difference between the empirical cumulative distribution functions (Figures S5B and S5C). This distance is expressed in amino acid units. Joint repertoire profiles were constructed by calculating the relative frequency of each combination of IGHV family, IGHJ family, and HCDR3 amino acid length. For each repertoire pair, profiles were aligned over the union of observed combinations, with absent combinations assigned a frequency of zero. Joint profiles were compared using Jensen–Shannon distance, calculated as the square root of the Jensen–Shannon divergence using base-2 logarithms (Figures S5D and S5E).

To evaluate the contribution of limited sampling to repertoire differences, 1,000 random subsamples were drawn without replacement from each RM’s MF uCDR3 repertoire. Each subsample contained the same number of entries as the corresponding RM’s xClonotype repertoire. For within-RM comparisons, the observed distance between the xClonotype and complete MF repertoires was compared with the reference distribution of distances between each MF subsample and its complete parental repertoire. For between-RM comparisons, all three animal pairs within each SHIV infection group were examined. The observed distance between the two xClonotype repertoires was compared with distances between independently drawn MF subsamples from the corresponding animals. The distance between the two complete MF repertoires was also calculated as a reference. All above comparison strategies were applied to both Wasserstein-1 and Jensen–Shannon distances.

### Software and computer programs used for data processing, analysis and visualization

The flow cytometry data were analyzed and plotted using FlowJo (v10.10). BCR V(D)J sequencing data were preprocessed with the custom program *AbSolute* (v1.0.0) and analyzed using the *IgBlast* program (v1.22.0) (https://ftp.ncbi.nih.gov/blast/executables/igblast/release/1.22.0/), the R package *scoper* (v1.3.0), and the Python packages *Change-O* (v1.3.4) and *IgPhyML* (v2.1.0). BCR Rep-Seq data were analyzed using the Python program *IgDiscover* (v0.12). Data from other assays were preprocessed in Microsoft Excel (v1808) and stored as .xlsx files. All data (except flow cytometry data) were processed, analyzed, and visualized using R (v4.5.2) and RStudio (v2025.09.2). Final figures were compiled with Adobe Illustrator (v29.7.1).

R packages utilized included: *openxlsx* (v4.2.8.1) for reading and writing .xlsx files; *tidyverse* (v2.0.0) and *magrittr* (v2.0.4) for data processing; *Biostrings* (v2.76.0) for DNA and protein sequence processing; *doParallel* (v1.0.17) for multi-thread processing; *msa* (v1.40.0) for Env sequence alignment (Figure 1B); *muscle* (v3.50.0) for IGV germline sequence alignment; *ape* (v5.8.1) for pairwise distance calculation with aligned IGV germline sequences; *uwot* (v0.2.3) for UMAP processing; *dbscan* (v1.2.3) for UMAP clustering analysis; *scoper* (v1.3.0) for V(D)J clonal analysis; *ggtree* (v3.16.3) for phylogenetic tree plotting; *jsonlite* (v2.0.0) to incorporate multiple phylogenetic trees into a single HTML file (Document S3); *data.table* (v1.17.8) for reading and processing of large Rep-Seq datasets. For general plotting, the following packages were used: *ggplot2* (v4.0.1), *egg* (v0.4.5), *cowplot* (v1.2.0), *patchwork* (v1.3.2), *scales* (v1.4.0), grid (v4.5.2), *ggnewscale* (v0.5.2), *ggpubr* (v0.6.2), and *gridExtra* (v2.3). Additional visualization packages included *ggalluvial* (v0.12.5) for alluvial plotting, *ggVennDiagram* (v1.5.4) for Venn diagram plotting, and *circlize* (v0.4.17) for chord diagram plotting.

