## Supplementary material for "Functional Convergence of Genetically Diverse B-Cell Receptors in Simian-HIV Infected Rhesus Macaques": Document S1

### BG505 Env amino−acid alignment

6 macaques | 74 sampled sequences | reference coverage: HXB2 1−709

Segment 1/8 | HXB2 1−91 | alignment columns 1−90

**Sequence identifier | GenBank accession**

Env region: SP (HXB2 1–32) | C1 (HXB2 33–91)

HXB2 numbering: 1 2 3 3a 3b 3c 4 5 6 7 8 9 10 15 16 17 18 19 20 21 22 23 24 25 26 27 28 29 30 31 32 33 34 35 36 37 38 39 40 41 42 43 44 45 46 47 48 49 50 51 52 53 54 55 56 57 58 59 60 61 62 63 64 65 66 67 68 69 70 71 72 73 74 75 76 77 78 79 80 81 82 83 84 85 86 87 88 89 90 91

| Sequence identifier \| GenBank accession | Alignment |
|---|---|
| **SHIV.A.BG505.T332N.375Y \| WNK77136** | MRVMGIQRNCQHLFRWGTMILGMIIICSAAENLWVTVYYGVPVWKDAETTLFCASDAKAYETEKHNVWATHACVPTDPNPQEIHLENVTE |
| **RM6708 \| n = 3 \| 88 WPI** | |
| RM6708.d0616.A03 \| MN468749 | ....E..................................................................................... |
| RM6708.d0616.A07 \| MN468750 | ....E..................................................................................... |
| RM6708.d0616.A08 \| MN468751 | ....E...................M................................................................. |
| **RMT283 \| n = 2 \| 88 WPI** | |
| RMT283.d0616.A01 \| MN469591 | .............L.....V.........V............................................................ |
| RMT283.d0616.A02 \| MN469592 | .............L.....V.........V............................................................ |
| **RM6706 \| n = 10 \| 88 WPI** | |
| RM6706.d0616.A03 \| MN468576 | .........Y................................................................................ |
| RM6706.d0616.A04 \| MN468577 | .........Y................................................................................ |
| RM6706.d0616.A05 \| MN468578 | .....T.................................................................................... |
| RM6706.d0616.A06 \| MN468579 | .........Y................................................................................ |
| RM6706.d0616.A10 \| MN468580 | .........Y................................................................................ |
| RM6706.d0616.A12 \| MN468581 | .........Y................................................................................ |
| RM6706.d0616.A13 \| MN468582 | .........Y................................................................................ |
| RM6706.d0616.A15 \| MN468583 | .........Y................................................................................ |
| RM6706.d0616.A16 \| MN468584 | .....T.................................................................................... |
| RM6706.d0616.A18 \| MN468585 | .....T.................................................................................... |
| **RM6715 \| n = 16 \| 88 WPI** | |
| RM6715.d0616.A01 \| MN468914 | ..............K.............T............................................................. |
| RM6715.d0616.A02 \| MN468915 | .............L............................................................................ |
| RM6715.d0616.A05 \| MN468917 | .............L............................................................................ |
| RM6715.d0616.A06 \| MN468918 | ..............K.............T............................................................. |
| RM6715.d0616.A07 \| MN468919 | ..............K.............T............................................................. |
| RM6715.d0616.A09 \| MN468920 | .............L............................................................................ |
| RM6715.d0616.A10 \| MN468921 | ..............K.............T..................................R.......................... |
| RM6715.d0616.A11 \| MN468922 | .............L..............T............................................................. |
| RM6715.d0616.A12 \| MN468923 | ..............K......E......T............................................................. |
| RM6715.d0616.A14 \| MN468924 | ..............K.............T............................................................K |
| RM6715.d0616.A15 \| MN468925 | ..............K.............T............................................................. |
| RM6715.d0616.B02 \| MN468926 | .............L............................................................................ |
| RM6715.d0616.B03 \| MN468927 | .............L............Y............................................................... |
| RM6715.d0616.B04 \| MN468928 | ..............K.............T............................................................. |
| RM6715.d0616.B06 \| MN468929 | .......K......K.............T............................................................. |
| RM6715.d0616.B07 \| MN468930 | .............L............................................................................ |
| **RM6718 \| n = 27 \| 88 WPI** | |
| RM6718.d0616.A01 \| MN469235 | ...........R..................G........................................................... |
| RM6718.d0616.A02 \| MN469236 | ..............................G........................................................... |
| RM6718.d0616.A04 \| MN469237 | ..............................G........................................................... |
| RM6718.d0616.A05 \| MN469238 | ..............................G........................................................... |
| RM6718.d0616.A08 \| MN469240 | ..............................G........................................................... |
| RM6718.d0616.A09 \| MN469241 | ..............................G........................................................... |
| RM6718.d0616.A11 \| MN469242 | .......................................................................................... |
| RM6718.d0616.A13 \| MN469243 | ..............................G........................................................... |
| RM6718.d0616.A14 \| MN469244 | ..............................G........................................................... |
| RM6718.d0616.A17 \| MN469246 | ..............................G........................................................... |
| RM6718.d0616.A18 \| MN469247 | ..............................G........................................................... |
| RM6718.d0616.A19 \| MN469248 | ..............................G........................................................... |
| RM6718.d0616.A21 \| MN469249 | ..............................G........................................................... |
| RM6718.d0616.A22 \| MN469250 | ..............................G........................................................... |
| RM6718.d0616.A24 \| MN469251 | ..............................G........................................................... |
| RM6718.d0616.A26 \| MN469252 | .......................................................................................... |
| RM6718.d0616.A28 \| MN469254 | ..............................G........................................................... |
| RM6718.d0616.A29 \| MN469255 | ..............................G........................................................... |
| RM6718.d0616.A30 \| MN469256 | ..............................G........................................................... |
| RM6718.d0616.A31 \| MN469257 | ..............................G........................................................... |
| RM6718.d0616.B01 \| MN469259 | ..............................G........................................................... |
| RM6718.d0616.B04 \| MN469260 | ..............................G........................................................... |
| RM6718.d0616.B05 \| MN469261 | ..............................G........................................................... |
| RM6718.d0616.B09 \| MN469262 | ...I..........................G........................................................... |
| RM6718.d0616.B10 \| MN469263 | ..............................G........................................................... |
| RM6718.d0616.B11 \| MN469264 | ..............................G........................................................... |
| RM6718.d0616.B13 \| MN469265 | ..............................G........................................................... |
| **RM6719 \| n = 16 \| 88 WPI** | |
| RM6719.d0616.A01 \| MN469516 | ..............................K........................................................... |
| RM6719.d0616.B01 \| MN469517 | ..............................K........................................................... |
| RM6719.d0616.B03 \| MN469518 | .............................V............................................................ |
| RM6719.d0616.B04 \| MN469519 | .............................V............................................................ |
| RM6719.d0616.B05 \| MN469520 | .............................V............................................................ |
| RM6719.d0616.B06 \| MN469521 | .............................V............................................................ |
| RM6719.d0616.B07 \| MN469522 | ..............................K........................................................... |
| RM6719.d0616.B09 \| MN469523 | .............................V............................................................ |
| RM6719.d0616.B11 \| MN469524 | ..............................K........................................................... |
| RM6719.d0616.B13 \| MN469525 | ..............................K........................................................... |
| RM6719.d0616.B14 \| MN469526 | ..............................G........................................................... |
| RM6719.d0616.B15 \| MN469527 | .............................V............................................................ |
| RM6719.d0616.B16 \| MN469528 | .............................V............................................................ |
| RM6719.d0616.B17 \| MN469529 | .............................V............................................................ |
| RM6719.d0616.B18 \| MN469530 | ..............................K........................................................... |
| RM6719.d0616.B19 \| MN469531 | ..............................K........................................................... |

. identity    A substitution    A insertion    − deletion    − gap at insertion column    X unresolved residue

* in numbering: insertion column without a supplied HXB2 position.

Domain and position annotations follow HXB2 numbering. SP, signal peptide; Fusion, fusion peptide.

MPER, membrane−proximal external region. Each sequence is displayed separately; no identical sequences are collapsed.

### BG505 Env amino−acid alignment

6 macaques | 74 sampled sequences | reference coverage: HXB2 1−709

Segment 2/8 | HXB2 92−184e | alignment columns 91−180

| Sequence identifier \| GenBank accession | |
|---|---|
| Env region | C1 (HXB2 92–130) / V1 (HXB2 131–157) / V2 (HXB2 158–184e) |
| HXB2 numbering | 92 93 94 95 96 97 98 99 100 101 102 103 104 105 106 107 108 109 110 111 112 113 114 115 116 117 118 119 120 121 122 123 124 125 126 127 128 129 130 131 132 133 134 135 136 137 138 139 140 141 142 151 152 153 154 155 156 157 158 159 160 161 162 163 164 165 166 167 168 169 170 171 172 173 174 175 176 177 178 179 180 181 182 183 184 184a 184b 184c 184d 184e |
| **SHIV.A.BG505.T332N.375Y \| WNK77136** | EFNMWKNNMVEQMHTDIISLWDQSLKPCVKLTPLCVTLQCTNVTNNITDDMRGELKNCSFNMTTELRDKKQKVYSLFYRLDVVQINENQG |
| **RM6708 \| n = 3 \| 88 WPI** | |
| RM6708.d0616.A03 \| MN468749 | .................T........................................................................ |
| RM6708.d0616.A07 \| MN468750 | .................T........................................................................ |
| RM6708.d0616.A08 \| MN468751 | ..............A........................................................................... |
| **RMT283 \| n = 2 \| 88 WPI** | |
| RMT283.d0616.A01 \| MN469591 | .............................E............................................................ |
| RMT283.d0616.A02 \| MN469592 | .............................E............................................................ |
| **RM6706 \| n = 10 \| 88 WPI** | |
| RM6706.d0616.A03 \| MN468576 | ..............A..............R............................................................ |
| RM6706.d0616.A04 \| MN468577 | ..............A..............R............................................................ |
| RM6706.d0616.A05 \| MN468578 | .................T.......M................................................................ |
| RM6706.d0616.A06 \| MN468579 | ..............A..............R............................................................ |
| RM6706.d0616.A10 \| MN468580 | ..............A..............R............................................................ |
| RM6706.d0616.A12 \| MN468581 | ..............A..............R............................................................ |
| RM6706.d0616.A13 \| MN468582 | ..............A..............R............................................................ |
| RM6706.d0616.A15 \| MN468583 | ..............A..............R............................................................ |
| RM6706.d0616.A16 \| MN468584 | .................T.......M................................................................ |
| RM6706.d0616.A18 \| MN468585 | .................T.......M................................................................ |
| **RM6715 \| n = 16 \| 88 WPI** | |
| RM6715.d0616.A01 \| MN468914 | ........................................A.....S.N......................................... |
| RM6715.d0616.A02 \| MN468915 | .............................E.....................GE..................................... |
| RM6715.d0616.A05 \| MN468917 | .............................E..................N..GE............................I........ |
| RM6715.d0616.A06 \| MN468918 | ................................................N..G...................................... |
| RM6715.d0616.A07 \| MN468919 | ........................................A.....S.N......................................... |
| RM6715.d0616.A09 \| MN468920 | .............................E.....................GE..................................... |
| RM6715.d0616.A10 \| MN468921 | .............................E..................N...E..................................... |
| RM6715.d0616.A11 \| MN468922 | .............................E................T.N......................................... |
| RM6715.d0616.A12 \| MN468923 | ................................................N..G...................................... |
| RM6715.d0616.A14 \| MN468924 | ................................................N..G...................................... |
| RM6715.d0616.A15 \| MN468925 | ................................................N..G...................................... |
| RM6715.d0616.B02 \| MN468926 | .............................E....................TG...................................... |
| RM6715.d0616.B03 \| MN468927 | .............................E...............D......E..................................... |
| RM6715.d0616.B04 \| MN468928 | ........................................A.....S.N......................................... |
| RM6715.d0616.B06 \| MN468929 | ................................................N..G...................................... |
| RM6715.d0616.B07 \| MN468930 | .............................E.....................GE..................................... |
| **RM6718 \| n = 27 \| 88 WPI** | |
| RM6718.d0616.A01 \| MN469235 | .................................................N........................................ |
| RM6718.d0616.A02 \| MN469236 | .......................................................................................... |
| RM6718.d0616.A04 \| MN469237 | .......................................................................................... |
| RM6718.d0616.A05 \| MN469238 | .......................................................................................... |
| RM6718.d0616.A08 \| MN469240 | .................................................G........................................ |
| RM6718.d0616.A09 \| MN469241 | .......................................................................................... |
| RM6718.d0616.A11 \| MN469242 | .......................................................................................... |
| RM6718.d0616.A13 \| MN469243 | ................................................N......................................... |
| RM6718.d0616.A14 \| MN469244 | .......................................................................................... |
| RM6718.d0616.A17 \| MN469246 | .....................................................................R.................... |
| RM6718.d0616.A18 \| MN469247 | .......................................................................................... |
| RM6718.d0616.A19 \| MN469248 | .......................................................................................... |
| RM6718.d0616.A21 \| MN469249 | ................................................N....................R.................... |
| RM6718.d0616.A22 \| MN469250 | .................................................G........................................ |
| RM6718.d0616.A24 \| MN469251 | .......................................................................................... |
| RM6718.d0616.A26 \| MN469252 | .......................................................................................... |
| RM6718.d0616.A28 \| MN469254 | .................................................G........................................ |
| RM6718.d0616.A29 \| MN469255 | ................................................N......................................... |
| RM6718.d0616.A30 \| MN469256 | ................................................N....................R.................... |
| RM6718.d0616.A31 \| MN469257 | ................................................N....................R.................... |
| RM6718.d0616.B01 \| MN469259 | .......................................................................................... |
| RM6718.d0616.B04 \| MN469260 | .......................................................................................... |
| RM6718.d0616.B05 \| MN469261 | .....................................................................R.................... |
| RM6718.d0616.B09 \| MN469262 | .................................................G........................................ |
| RM6718.d0616.B10 \| MN469263 | .......................................................................................... |
| RM6718.d0616.B11 \| MN469264 | .......................................................................................... |
| RM6718.d0616.B13 \| MN469265 | .................................................G........................................ |
| **RM6719 \| n = 16 \| 88 WPI** | |
| RM6719.d0616.A01 \| MN469516 | .......................................................................................... |
| RM6719.d0616.B01 \| MN469517 | .......................................................................................... |
| RM6719.d0616.B03 \| MN469518 | .......................................................................................... |
| RM6719.d0616.B04 \| MN469519 | .......................................................................................... |
| RM6719.d0616.B05 \| MN469520 | .......................................................................................... |
| RM6719.d0616.B06 \| MN469521 | .......................................................................................... |
| RM6719.d0616.B07 \| MN469522 | .......................................................................................... |
| RM6719.d0616.B09 \| MN469523 | .......................................................................................... |
| RM6719.d0616.B11 \| MN469524 | .......................................................................................... |
| RM6719.d0616.B13 \| MN469525 | .......................................................................................... |
| RM6719.d0616.B14 \| MN469526 | .......................................................................................... |
| RM6719.d0616.B15 \| MN469527 | .......................................................................................... |
| RM6719.d0616.B16 \| MN469528 | .......................................................................................... |
| RM6719.d0616.B17 \| MN469529 | .......................................................................................... |
| RM6719.d0616.B18 \| MN469530 | .......................................................................................... |
| RM6719.d0616.B19 \| MN469531 | .......................................................................................... |

. identity    A substitution    A insertion    − deletion    − gap at insertion column    X unresolved residue

* in numbering: insertion column without a supplied HXB2 position.

Domain and position annotations follow HXB2 numbering. SP, signal peptide; Fusion, fusion peptide.

MPER, membrane−proximal external region. Each sequence is displayed separately; no identical sequences are collapsed.

### BG505 Env amino−acid alignment

6 macaques | 74 sampled sequences | reference coverage: HXB2 1−709

Segment 3/8 | HXB2 184f−271 | alignment columns 181−270

**Sequence identifier | GenBank accession**

Env region: V2 | C2

HXB2 numbering: 184f 184g 184h 185 186 187 188 189 190 191 192 193 194 195 196 197 198 199 200 201 202 203 204 205 206 207 208 209 210 211 212 213 214 215 216 217 218 219 220 221 222 223 224 225 226 227 228 229 230 231 232 233 234 235 236 237 238 239 240 241 242 243 244 245 246 247 248 249 250 251 252 253 254 255 256 257 258 259 260 261 262 263 264 265 266 267 268 269 270 271

```
SHIV.A.BG505.T332N.375Y | WNK77136   NRSNNSNKEYRLINCNTSAITQACPKVSFEPIPIHYCAPAGFAILKCKDKKFNGTGPCPSVSTVQCTHGIKPVVSTQLLLNGSLAEEEVM

RM6708 | n = 3 | 88 WPI
RM6708.d0616.A03 | MN468749   ...............................................E...........NA.............................
RM6708.d0616.A07 | MN468750   ...............................................E...........N..............................
RM6708.d0616.A08 | MN468751   ...............................................E...........N..........................G...

RMT283 | n = 2 | 88 WPI
RMT283.d0616.A01 | MN469591   ..........................................................................................
RMT283.d0616.A02 | MN469592   ..........................................................................................

RM6706 | n = 10 | 88 WPI
RM6706.d0616.A03 | MN468576   ..........................................................................................
RM6706.d0616.A04 | MN468577   ..........................................................................................
RM6706.d0616.A05 | MN468578   ..........................................................................................
RM6706.d0616.A06 | MN468579   ..........................................................................................
RM6706.d0616.A10 | MN468580   ..........................................................................................
RM6706.d0616.A12 | MN468581   ..........................................................................................
RM6706.d0616.A13 | MN468582   ..........................................................................................
RM6706.d0616.A15 | MN468583   ..........................................................................................
RM6706.d0616.A16 | MN468584   ..........................................................S...............................
RM6706.d0616.A18 | MN468585   ..........................................................S...........................G...

RM6715 | n = 16 | 88 WPI
RM6715.d0616.A01 | MN468914   ..........................................................................................
RM6715.d0616.A02 | MN468915   ...............................................T..........................................
RM6715.d0616.A05 | MN468917   ..........................................................................................
RM6715.d0616.A06 | MN468918   ...............................................T..........................................
RM6715.d0616.A07 | MN468919   ..........................................................................................
RM6715.d0616.A09 | MN468920   ......................................................................................G...
RM6715.d0616.A10 | MN468921   ..........................................................................................
RM6715.d0616.A11 | MN468922   ......................................................................................G...
RM6715.d0616.A12 | MN468923   .......E.......................................T..........................................
RM6715.d0616.A14 | MN468924   ...............................................T......................................K...
RM6715.d0616.A15 | MN468925   ...............................................T..........................................
RM6715.d0616.B02 | MN468926   ......................................................................................G...
RM6715.d0616.B03 | MN468927   ..................................................M....................................K..
RM6715.d0616.B04 | MN468928   ..........................................................................................
RM6715.d0616.B06 | MN468929   ...............................................T..........................................
RM6715.d0616.B07 | MN468930   ..........................................................................................

RM6718 | n = 27 | 88 WPI
RM6718.d0616.A01 | MN469235   ....S.............................................E........N..........................G...
RM6718.d0616.A02 | MN469236   ..................................................E........N..........................G...
RM6718.d0616.A04 | MN469237   ..................................................E........N..........................G...
RM6718.d0616.A05 | MN469238   ..................................................E........N..........................G...
RM6718.d0616.A08 | MN469240   ..................................................E........N..........................G...
RM6718.d0616.A09 | MN469241   ..................................................E........N..........................G...
RM6718.d0616.A11 | MN469242   ..................................................E........N..........................G...
RM6718.d0616.A13 | MN469243   ..................................................E........N..........................G...
RM6718.d0616.A14 | MN469244   ..................................................E........N..........................G...
RM6718.d0616.A17 | MN469246   ....S.............................................E........N..........................G...
RM6718.d0616.A18 | MN469247   ..................................................E........N..........................G...
RM6718.d0616.A19 | MN469248   ..................................................E........N..........................G...
RM6718.d0616.A21 | MN469249   ....S.............................................E........N..........................G...
RM6718.d0616.A22 | MN469250   ..................................................E........N..........................G...
RM6718.d0616.A24 | MN469251   ....S.............................................E........N..........................G...
RM6718.d0616.A26 | MN469252   ..................................................E........N..........................G...
RM6718.d0616.A28 | MN469254   .................................................E.........N..........................G...
RM6718.d0616.A29 | MN469255   ..................................................E........N..........................G...
RM6718.d0616.A30 | MN469256   ....S.............................................E........N..........................G...
RM6718.d0616.A31 | MN469257   ....S.............................................E........N..........................G...
RM6718.d0616.B01 | MN469259   ..................................................E........N..........................G...
RM6718.d0616.B04 | MN469260   ....S............................................E.........N..........................G...
RM6718.d0616.B05 | MN469261   ....S.....................................................QN..........................G...
RM6718.d0616.B09 | MN469262   ..................................................E........N..........................G...
RM6718.d0616.B10 | MN469263   ..................................................E........N..........................R...
RM6718.d0616.B11 | MN469264   ..................................................E........N..........................G...
RM6718.d0616.B13 | MN469265   ..................................................E........N..........................G...

RM6719 | n = 16 | 88 WPI
RM6719.d0616.A01 | MN469516   ..................................................E...................................G...
RM6719.d0616.B01 | MN469517   ..................................................E...................................G...
RM6719.d0616.B03 | MN469518   .....................................................................................QG...
RM6719.d0616.B04 | MN469519   ................A....................................................................QG...
RM6719.d0616.B05 | MN469520   .....................................................................................QG...
RM6719.d0616.B06 | MN469521   .....................................................................................QG...
RM6719.d0616.B07 | MN469522   ..................................................E...................................G...
RM6719.d0616.B09 | MN469523   .....................................................................................QG...
RM6719.d0616.B11 | MN469524   ......................................................................................G...
RM6719.d0616.B13 | MN469525   ..................................................E...................................GK..
RM6719.d0616.B14 | MN469526   ......................................................................................G...
RM6719.d0616.B15 | MN469527   .....................................................................................QG...
RM6719.d0616.B16 | MN469528   .....................................................................................QG...
RM6719.d0616.B17 | MN469529   .....................................................................................QG...
RM6719.d0616.B18 | MN469530   ..................................................E...................................G...
RM6719.d0616.B19 | MN469531   ..................................................E...................................G...
```

. identity A substitution A insertion − deletion − gap at insertion column X unresolved residue

* in numbering: insertion column without a supplied HXB2 position.

Domain and position annotations follow HXB2 numbering. SP, signal peptide; Fusion, fusion peptide.

MPER, membrane−proximal external region. Each sequence is displayed separately; no identical sequences are collapsed.

### BG505 Env amino−acid alignment

6 macaques | 74 sampled sequences | reference coverage: HXB2 1−709

Segment 4/8 | HXB2 272−362 | alignment columns 271−360

**Sequence identifier | GenBank accession**

Env region: C2 (HXB2 272–295) | V3 (HXB2 296–330) | C3 (HXB2 331–362)

HXB2 numbering: 272 273 274 275 276 277 278 279 280 281 282 283 284 285 286 287 288 289 290 291 292 293 294 295 296 297 298 299 300 301 302 303 304 305 306 307 308 309 312 313 314 315 316 317 318 319 320 321 322 322a 323 324 325 326 327 328 329 330 331 332 333 334 335 336 337 338 339 340 341 342 343 344 345 346 347 348 349 350 351 352 353 354 355 356 357 358 359 360 361 362

| Sequence identifier | GenBank accession | Alignment (HXB2 272–362) |
|---|---|---|
| **SHIV.A.BG505.T332N.375Y** | **WNK77136** | IRSENITNNAKNILVQFNTPVQINCTRPNNNTRKSIRIGPGQAFYATGDIIGDIRQAHCNVSKATWNETLGKVVKQLRKHFGNNTIIRFA |
| **RM6708 \| n = 3 \| 88 WPI** | | |
| RM6708.d0616.A03 | MN468749 | ....................................H..............................K..E..A.............K.. |
| RM6708.d0616.A07 | MN468750 | ....................................H..............................K..E..A.............K.. |
| RM6708.d0616.A08 | MN468751 | .................D..................H..............................K..E................K.. |
| **RMT283 \| n = 2 \| 88 WPI** | | |
| RMT283.d0616.A01 | MN469591 | ..........................................................................E......R.....G.. |
| RMT283.d0616.A02 | MN469592 | ..........................................................................E......R.....G.. |
| **RM6706 \| n = 10 \| 88 WPI** | | |
| RM6706.d0616.A03 | MN468576 | .................D........................................................E.......S..T.... |
| RM6706.d0616.A04 | MN468577 | .................D........................................................E.......S..T.... |
| RM6706.d0616.A05 | MN468578 | ..........................................................................E..........T.... |
| RM6706.d0616.A06 | MN468579 | .................D........................................................E.......S..T.... |
| RM6706.d0616.A10 | MN468580 | .................D........................................................E.......S..T.... |
| RM6706.d0616.A12 | MN468581 | .................D........................................................E.......S..T.... |
| RM6706.d0616.A13 | MN468582 | .................D........................................................E.......S..T.... |
| RM6706.d0616.A15 | MN468583 | .................D........................................................E.......S..T.... |
| RM6706.d0616.A16 | MN468584 | .......................................................................E.............T.... |
| RM6706.d0616.A18 | MN468585 | .......................................................................R.............T.... |
| **RM6715 \| n = 16 \| 88 WPI** | | |
| RM6715.d0616.A01 | MN468914 | .........................I......................................I.........ER.........T.... |
| RM6715.d0616.A02 | MN468915 | .................H.......I................................................E..........T.... |
| RM6715.d0616.A05 | MN468917 | .................K.......I...........................V.............K......E.........−T.... |
| RM6715.d0616.A06 | MN468918 | .................H.......I................................................E..........T.... |
| RM6715.d0616.A07 | MN468919 | .........................I......................................I.........ER.........T.... |
| RM6715.d0616.A09 | MN468920 | .................D.......I..........H..................................R..E.........−T.... |
| RM6715.d0616.A10 | MN468921 | .................K.......I.........................................K..R...E.........−T.... |
| RM6715.d0616.A11 | MN468922 | .........................I..........H.....................................E...E.....−T.... |
| RM6715.d0616.A12 | MN468923 | .................H.......I................................................E..........T.... |
| RM6715.d0616.A14 | MN468924 | .................H.......I.........................................K......E..........T.... |
| RM6715.d0616.A15 | MN468925 | .................H.......I................................................E..........T.... |
| RM6715.d0616.B02 | MN468926 | .................D.......I..........H.................................R...E.........−T.... |
| RM6715.d0616.B03 | MN468927 | .........................I..........H...E.................................E..........T.... |
| RM6715.d0616.B04 | MN468928 | .........................I......................................I.........ER.........T.... |
| RM6715.d0616.B06 | MN468929 | .................H.......I................................................E..........T.... |
| RM6715.d0616.B07 | MN468930 | .................K.......I...........................V.............K......E...E.....−T.... |
| **RM6718 \| n = 27 \| 88 WPI** | | |
| RM6718.d0616.A01 | MN469235 | .................D.......I..........H.............................IK......E.......S..T.K.. |
| RM6718.d0616.A02 | MN469236 | .................D.......I..........H.........................E....K......Q.......S..T.K.. |
| RM6718.d0616.A04 | MN469237 | .................D..................H.........................E....K......E.......S..T.... |
| RM6718.d0616.A05 | MN469238 | .................D.......I..........H.........................E....K......Q.......S..T.K.. |
| RM6718.d0616.A08 | MN469240 | .................D.......I..........H..........................T......R...E.......S..T.... |
| RM6718.d0616.A09 | MN469241 | .................D.......I..........H.........................E...YK......Q.......S..T.K.. |
| RM6718.d0616.A11 | MN469242 | .................D..................H.........................E.......R...E.......S..T.... |
| RM6718.d0616.A13 | MN469243 | .................D.......I..........H.........................E....K......Q.......S..T.... |
| RM6718.d0616.A14 | MN469244 | .................D..................H..........................T......R...E.......S..T.... |
| RM6718.d0616.A17 | MN469246 | .................D.......I..........H.........................E....K......Q.......S..T.K.. |
| RM6718.d0616.A18 | MN469247 | .................D.......I..........H.........................E....K......Q.......S..T.K.. |
| RM6718.d0616.A19 | MN469248 | .................D..................H.........................E....K......E.......S..T.... |
| RM6718.d0616.A21 | MN469249 | .................D..................H.........................R...D...R...E.......S..T.... |
| RM6718.d0616.A22 | MN469250 | .................D..................H.............................DK......E.......S..T.... |
| RM6718.d0616.A24 | MN469251 | .................D..................H.........................R....K......E.......S..T.K.. |
| RM6718.d0616.A26 | MN469252 | .................D..................H.................................R...E.......S..T.... |
| RM6718.d0616.A28 | MN469254 | .................D.......I..........H..........................T......R...E.......S..T.... |
| RM6718.d0616.A29 | MN469255 | .................D..................H.............................Y.......E........T.T.... |
| RM6718.d0616.A30 | MN469256 | .................D.......I..........H.....................................N.......S..T.... |
| RM6718.d0616.A31 | MN469257 | .................D..................H.........................R...D...R...E.......S..T.... |
| RM6718.d0616.B01 | MN469259 | .................D.......I..........H.........................E....K......Q.......S..T.K.. |
| RM6718.d0616.B04 | MN469260 | .................D..................H.........................E....K......E.......S..T.K.. |
| RM6718.d0616.B05 | MN469261 | .................D..................H.........................E....K......E.......S..T.... |
| RM6718.d0616.B09 | MN469262 | .................D.......I..........H.........................R...........Q......RS..T.K.. |
| RM6718.d0616.B10 | MN469263 | .................D.......I..........H.........................E....K......Q.......S..T.K.. |
| RM6718.d0616.B11 | MN469264 | .................D.......I..........H..............................K......Q.......S..T.... |
| RM6718.d0616.B13 | MN469265 | .................D.......I..........H..........................T......R...E.......S..T.... |
| **RM6719 \| n = 16 \| 88 WPI** | | |
| RM6719.d0616.A01 | MN469516 | ....................................H....................................AE......RD....... |
| RM6719.d0616.B01 | MN469517 | .................D..................H..............................K.....AE...R..R.D...... |
| RM6719.d0616.B03 | MN469518 | ...................S................H.....................................E.......D..T.... |
| RM6719.d0616.B04 | MN469519 | ...................S................H.....................................E.......D..T.... |
| RM6719.d0616.B05 | MN469520 | ...................S................H.....................................E.......D..T.... |
| RM6719.d0616.B06 | MN469521 | ...................S................H.....................................E.......D..T.... |
| RM6719.d0616.B07 | MN469522 | ....................................H..................................E.AE......R........ |
| RM6719.d0616.B09 | MN469523 | ...................S................H.....................................E.......D..T.... |
| RM6719.d0616.B11 | MN469524 | ....................................H....................................AE...R..R........ |
| RM6719.d0616.B13 | MN469525 | ....................................H....................................AE......R........ |
| RM6719.d0616.B14 | MN469526 | ....................................H...........................A........AE...R..R........ |
| RM6719.d0616.B15 | MN469527 | ...................S................H.....................................E.......D..T.... |
| RM6719.d0616.B16 | MN469528 | ...................S................H.....................................E.......D..T.... |
| RM6719.d0616.B17 | MN469529 | ...................S................H.....................................E.......D..T.... |
| RM6719.d0616.B18 | MN469530 | ....................................H....................................AE......R........ |
| RM6719.d0616.B19 | MN469531 | ....................................H....................................AE......RD....... |

. identity | A substitution | A insertion | − deletion | − gap at insertion column | X unresolved residue

* in numbering: insertion column without a supplied HXB2 position.

Domain and position annotations follow HXB2 numbering. SP, signal peptide; Fusion, fusion peptide.

MPER, membrane−proximal external region. Each sequence is displayed separately; no identical sequences are collapsed.

### BG505 Env amino−acid alignment

6 macaques | 74 sampled sequences | reference coverage: HXB2 1−709

Segment 5/8 | HXB2 363−453 | alignment columns 361−450

**Sequence identifier | GenBank accession**

Env region: C3 (HXB2 363–384) | V4 (HXB2 385–418) | C4 (HXB2 419–453)

HXB2 numbering: 363 364 365 366 367 368 369 370 371 372 373 374 375 376 377 378 379 380 381 382 383 384 385 386 387 388 389 390 391 392 393 394 395 396 397 398 399 401 402 403 404 405 406 407 408 409 410 411 412 413 414 415 416 417 418 419 420 421 422 423 424 425 426 427 428 429 430 431 432 433 434 435 436 437 438 439 440 441 442 443 444 445 446 447 448 449 450 451 452 453

```
SHIV.A.BG505.T332N.375Y | WNK77136   NSSGGDLEVTTHYFNCGGEFFYCNTSGLFNSTWISNTSVQGSNSTGSNDSITLPCRIKQIINMWQRIGQAMYAPPIQGVIRCVSNITGLI
```

**RM6708 | n = 3 | 88 WPI**

```
RM6708.d0616.A03 | MN468749   S.................................T......E.................................................
RM6708.d0616.A07 | MN468750   S.................................T......E.................................................
RM6708.d0616.A08 | MN468751   .................................TN.......................................................
```

**RMT283 | n = 2 | 88 WPI**

```
RMT283.d0616.A01 | MN469591   S................................T..........................................R.............
RMT283.d0616.A02 | MN469592   S................................T..........................................R.............
```

**RM6706 | n = 10 | 88 WPI**

```
RM6706.d0616.A03 | MN468576   .................................N........................................................
RM6706.d0616.A04 | MN468577   .................................N........................................................
RM6706.d0616.A05 | MN468578   .................................N........................................................
RM6706.d0616.A06 | MN468579   .................................N........................................................
RM6706.d0616.A10 | MN468580   .................................N........................................................
RM6706.d0616.A12 | MN468581   .................................N........................................................
RM6706.d0616.A13 | MN468582   .................................N........................................................
RM6706.d0616.A15 | MN468583   .................................N........................................................
RM6706.d0616.A16 | MN468584   ............H....................N...............N........................................
RM6706.d0616.A18 | MN468585   ............H....................N........................................................
```

**RM6715 | n = 16 | 88 WPI**

```
RM6715.d0616.A01 | MN468914   .................................T........................................................
RM6715.d0616.A02 | MN468915   .................................T........................................................
RM6715.d0616.A05 | MN468917   .................................T...............N........................................
RM6715.d0616.A06 | MN468918   .................................T........................................................
RM6715.d0616.A07 | MN468919   .................................T........................................................
RM6715.d0616.A09 | MN468920   S................................T.............S..........................................
RM6715.d0616.A10 | MN468921   .................................T...............N........................................
RM6715.d0616.A11 | MN468922   S................................T........D...............................................
RM6715.d0616.A12 | MN468923   .................................T..........A.P...........................................
RM6715.d0616.A14 | MN468924   ....E............................T.........N..............................................
RM6715.d0616.A15 | MN468925   .................................T........................................................
RM6715.d0616.B02 | MN468926   .................................T...............N........................................
RM6715.d0616.B03 | MN468927   .................................T...........R............................................
RM6715.d0616.B04 | MN468928   .................................T........................................................
RM6715.d0616.B06 | MN468929   .................................T........................................................
RM6715.d0616.B07 | MN468930   .................................T...............N........................................
```

**RM6718 | n = 27 | 88 WPI**

```
RM6718.d0616.A01 | MN469235   ..................................G..........E............................................
RM6718.d0616.A02 | MN469236   ..................................G............D.N........................................
RM6718.d0616.A04 | MN469237   .............................................E..N.................V.......................
RM6718.d0616.A05 | MN469238   ..................................G..........E............................................
RM6718.d0616.A08 | MN469240   ..................................G..........E...N................V.......................
RM6718.d0616.A09 | MN469241   ..................................G............D.N........................................
RM6718.d0616.A11 | MN469242   ..................................G............SN.................V.......................
RM6718.d0616.A13 | MN469243   ..................................G............D.N........................................
RM6718.d0616.A14 | MN469244   ..................................G..........EY--.................V.......................
RM6718.d0616.A17 | MN469246   ..................................G..........E....................V.......................
RM6718.d0616.A18 | MN469247   .................................VG.........................................R.............
RM6718.d0616.A19 | MN469248   .............................................E..N.................V.......................
RM6718.d0616.A21 | MN469249   ..................................G...........LS..................V.......................
RM6718.d0616.A22 | MN469250   .................................TG...A......E...N........................................
RM6718.d0616.A24 | MN469251   ..................................G...........P..N........................................
RM6718.d0616.A26 | MN469252   ..................................G...........L--.................V.......................
RM6718.d0616.A28 | MN469254   ..................................G..........E...N................V.......................
RM6718.d0616.A29 | MN469255   ..................................G...........P.............................R.............
RM6718.d0616.A30 | MN469256   .............................................E..G.........................................
RM6718.d0616.A31 | MN469257   ..................................G...........LS..................V.......................
RM6718.d0616.B01 | MN469259   ..................................G.......K..E....................V.......................
RM6718.d0616.B04 | MN469260   ..................................G..........EP.G.........................................
RM6718.d0616.B05 | MN469261   .............................................E..GN..........................R.............
RM6718.d0616.B09 | MN469262   ..................................G..........E...N........................................
RM6718.d0616.B10 | MN469263   ..................................G...........LS..................V.......................
RM6718.d0616.B11 | MN469264   ..................................G...........P.G.........................................
RM6718.d0616.B13 | MN469265   ..................................G..........E...N................V.......................
```

**RM6719 | n = 16 | 88 WPI**

```
RM6719.d0616.A01 | MN469516   .................................V......R..N..............................................
RM6719.d0616.B01 | MN469517   ..........................................K...............................................
RM6719.d0616.B03 | MN469518   S.......................................R..N..............................................
RM6719.d0616.B04 | MN469519   S.......................................RP.N..............................................
RM6719.d0616.B05 | MN469520   S.......................................R..N..............................................
RM6719.d0616.B06 | MN469521   S.......................................R..N..............................................
RM6719.d0616.B07 | MN469522   .................................V......R..N..............................................
RM6719.d0616.B09 | MN469523   S.......................................R..N..............................................
RM6719.d0616.B11 | MN469524   .......................................RR.................................................
RM6719.d0616.B13 | MN469525   .................................V......R..N..............................................
RM6719.d0616.B14 | MN469526   .........................................L................................................
RM6719.d0616.B15 | MN469527   S.......................................R..NM.............................................
RM6719.d0616.B16 | MN469528   S.......................................R..N..............................................
RM6719.d0616.B17 | MN469529   S.......................................R..N..............................................
RM6719.d0616.B18 | MN469530   ..................K..............V....M.R..N..............................................
RM6719.d0616.B19 | MN469531   .................................V......R..N..............................................
```

. identity A substitution A insertion − deletion − gap at insertion column X unresolved residue

* in numbering: insertion column without a supplied HXB2 position.

Domain and position annotations follow HXB2 numbering. SP, signal peptide; Fusion, fusion peptide.

MPER, membrane−proximal external region. Each sequence is displayed separately; no identical sequences are collapsed.

### BG505 Env amino−acid alignment

6 macaques | 74 sampled sequences | reference coverage: HXB2 1−709

Segment 6/8 | HXB2 454−535 | alignment columns 451−540

**Sequence identifier | GenBank accession**

Env region: C4 | V5 | C5 | Fusion

HXB2 numbering: 454 455 456 457 458 459 460 * * * * * * 461 462 463 464 * * 465 466 467 468 469 470 471 472 473 474 475 476 477 478 479 480 481 482 483 484 485 486 487 488 489 490 491 492 493 494 495 496 497 498 499 500 501 502 503 504 505 506 507 508 509 510 511 512 513 514 515 516 517 518 519 520 521 522 523 524 525 526 527 528 529 530 531 532 533 534 535

```
SHIV.A.BG505.T332N.375Y | WNK77136   LTRDGGS------TNST--TETFRPGGGDMRDNWRSELYKYKVVKIEPLGVAPTRAKRRVVGREKRAVGIGAVFLGFLGAAGSTMGAASM
```

**RM6708 | n = 3 | 88 WPI**

```
RM6708.d0616.A03 | MN468749   .......------..D.--..........................................E............................
RM6708.d0616.A07 | MN468750   .......------..D.--..........................................E............................
RM6708.d0616.A08 | MN468751   .......------..NA--..........................................E............................
```

**RMT283 | n = 2 | 88 WPI**

```
RMT283.d0616.A01 | MN469591   .......------....--.......................................................................
RMT283.d0616.A02 | MN469592   .......------....--.......................................................................
```

**RM6706 | n = 10 | 88 WPI**

```
RM6706.d0616.A03 | MN468576   .......------....--.......................................................................
RM6706.d0616.A04 | MN468577   .......------....--.......................................................................
RM6706.d0616.A05 | MN468578   .......------..N.--.......................................................................
RM6706.d0616.A06 | MN468579   .......------....--.......................................................................
RM6706.d0616.A10 | MN468580   .......------....--.......................................................................
RM6706.d0616.A12 | MN468581   .......------....--.......................................................................
RM6706.d0616.A13 | MN468582   .......------....--.......................................................................
RM6706.d0616.A15 | MN468583   .......------....--.......................................................................
RM6706.d0616.A16 | MN468584   .......------..N.--..I....................................................................
RM6706.d0616.A18 | MN468585   .......------..N.--.......................................................................
```

**RM6715 | n = 16 | 88 WPI**

```
RM6715.d0616.A01 | MN468914   .......------.DN.--..........................................E............................
RM6715.d0616.A02 | MN468915   ......N------.DN.--..........................................E...........................I
RM6715.d0616.A05 | MN468917   ......N------.DN.--..........................................E............................
RM6715.d0616.A06 | MN468918   ......N------.DN.--..........................................E............................
RM6715.d0616.A07 | MN468919   .......------.DN.--..........................................E............................
RM6715.d0616.A09 | MN468920   ......N------.DN.--..........................................E............................
RM6715.d0616.A10 | MN468921   ......N------.DN.--..........................................E............................
RM6715.d0616.A11 | MN468922   ......N------.DN.--..........................................E............................
RM6715.d0616.A12 | MN468923   ......N------.DN.--..........................................E............................
RM6715.d0616.A14 | MN468924   ......N------.DN.--..........................................E.............R..............
RM6715.d0616.A15 | MN468925   ......N------.DN.--..........................................E............................
RM6715.d0616.B02 | MN468926   ......N------.DN.--..........................................E............................
RM6715.d0616.B03 | MN468927   .......------....TE..........................................E............................
RM6715.d0616.B04 | MN468928   .......------.DN.--..........................................E............................
RM6715.d0616.B06 | MN468929   ......N------.DN.--..........................................E............................
RM6715.d0616.B07 | MN468930   ......N------.DN.--..........................................E............................
```

**RM6718 | n = 27 | 88 WPI**

```
RM6718.d0616.A01 | MN469235   .......TNS---..G.--.....................................R.................................
RM6718.d0616.A02 | MN469236   .......TNS---....--.......................................................................
RM6718.d0616.A04 | MN469237   .......TNS---....--.......................................................................
RM6718.d0616.A05 | MN469238   .......------..N.--.....................................R.................................
RM6718.d0616.A08 | MN469240   ..K....TNS---....--.......................................................................
RM6718.d0616.A09 | MN469241   .......TNS---....--.......................................................................
RM6718.d0616.A11 | MN469242   .......TNS---....--.......................................................................
RM6718.d0616.A13 | MN469243   .......TNS---....--.......................................................................
RM6718.d0616.A14 | MN469244   .......TNS---....--.......................................................................
RM6718.d0616.A17 | MN469246   .......TNS---....--.......................................................................
RM6718.d0616.A18 | MN469247   .......TNS---..G.--.....................................R.................................
RM6718.d0616.A19 | MN469248   .......TNS---....--.......................................................................
RM6718.d0616.A21 | MN469249   .......TNS---....--..........................................E............................
RM6718.d0616.A22 | MN469250   .......TDSTTE....--.......................................................................
RM6718.d0616.A24 | MN469251   .......TNS---....--.......................................................................
RM6718.d0616.A26 | MN469252   .......TNS---....--..........................................E............................
RM6718.d0616.A28 | MN469254   .......TNS---....--.......................................................................
RM6718.d0616.A29 | MN469255   .......TNS---..G.--.....................................R.................................
RM6718.d0616.A30 | MN469256   .......TNS---..G.--.....................................R.................................
RM6718.d0616.A31 | MN469257   .......TNS---....--..........................................E............................
RM6718.d0616.B01 | MN469259   .......TNS---....--.......................................................................
RM6718.d0616.B04 | MN469260   .......TNS---....--.......................................................................
RM6718.d0616.B05 | MN469261   ......G------NTN.--.......................................................................
RM6718.d0616.B09 | MN469262   .......TNS---....--...............................I.....R.................................
RM6718.d0616.B10 | MN469263   .......TNS---....--.......................................................................
RM6718.d0616.B11 | MN469264   .......TDSTTE....--I......................................................................
RM6718.d0616.B13 | MN469265   .......TNS---....--.......................................................................
```

**RM6719 | n = 16 | 88 WPI**

```
RM6719.d0616.A01 | MN469516   .......------P.N.--......................................................................I
RM6719.d0616.B01 | MN469517   .......------P.N.--......................................................................I
RM6719.d0616.B03 | MN469518   .......------P.N.--......................................................................I
RM6719.d0616.B04 | MN469519   .......------P.N.--......................................................................I
RM6719.d0616.B05 | MN469520   .......------P.N.--......................................................................I
RM6719.d0616.B06 | MN469521   .......------P.N.--......................................................................I
RM6719.d0616.B07 | MN469522   .......------P.N.--......................................................................I
RM6719.d0616.B09 | MN469523   .......------P.N.--......................................................................I
RM6719.d0616.B11 | MN469524   .......------P.N.--......................................................................I
RM6719.d0616.B13 | MN469525   .......------P.N.--......................................................................I
RM6719.d0616.B14 | MN469526   .......------..N.--......................................................................I
RM6719.d0616.B15 | MN469527   .......------P.N.--......................................................................I
RM6719.d0616.B16 | MN469528   .......------P.N.--......................................................................I
RM6719.d0616.B17 | MN469529   .......------P.N.--......................................................................I
RM6719.d0616.B18 | MN469530   .......------P.N.--.................K...............................E....................I
RM6719.d0616.B19 | MN469531   .......------P.N.--......................................................................I
```

. identity    A substitution    A insertion    − deletion    − gap at insertion column    X unresolved residue

* in numbering: insertion column without a supplied HXB2 position.

Domain and position annotations follow HXB2 numbering. SP, signal peptide; Fusion, fusion peptide.

MPER, membrane−proximal external region. Each sequence is displayed separately; no identical sequences are collapsed.

### BG505 Env amino−acid alignment

6 macaques | 74 sampled sequences | reference coverage: HXB2 1−709

Segment 7/8 | HXB2 536−625 | alignment columns 541−630

**Sequence identifier | GenBank accession**

Env region

HXB2 numbering: 536–625

| Sequence identifier \| GenBank accession | HXB2 536–625 |
|---|---|
| **SHIV.A.BG505.T332N.375Y \| WNK77136** | TLTVQARNLLSGIVQQQSNLLRAIEAQQHLLKLTVWGIKQLQARVLAVERYLRDQQLLGIWGCSGKLICTTNVPWNSSWSNRNLSEIWDN |
| **RM6708 \| n = 3 \| 88 WPI** | |
| RM6708.d0616.A03 \| MN468749 | .......................................................................................... |
| RM6708.d0616.A07 \| MN468750 | .......................................................................................... |
| RM6708.d0616.A08 \| MN468751 | .......................................................................................... |
| **RMT283 \| n = 2 \| 88 WPI** | |
| RMT283.d0616.A01 \| MN469591 | ....................................................................................N..... |
| RMT283.d0616.A02 \| MN469592 | ....................................................................................N..... |
| **RM6706 \| n = 10 \| 88 WPI** | |
| RM6706.d0616.A03 \| MN468576 | .......................................................................................... |
| RM6706.d0616.A04 \| MN468577 | .......................................................................................... |
| RM6706.d0616.A05 \| MN468578 | .......................................................................................... |
| RM6706.d0616.A06 \| MN468579 | .......................................................................................... |
| RM6706.d0616.A10 \| MN468580 | .......................................................................................... |
| RM6706.d0616.A12 \| MN468581 | .....................................V.................................................... |
| RM6706.d0616.A13 \| MN468582 | .......................................................................................... |
| RM6706.d0616.A15 \| MN468583 | .......................................................................................... |
| RM6706.d0616.A16 \| MN468584 | .......................................................................................... |
| RM6706.d0616.A18 \| MN468585 | .......................................................................................... |
| **RM6715 \| n = 16 \| 88 WPI** | |
| RM6715.d0616.A01 \| MN468914 | .......................................................................................... |
| RM6715.d0616.A02 \| MN468915 | .......................................................................................... |
| RM6715.d0616.A05 \| MN468917 | .......................................................................................... |
| RM6715.d0616.A06 \| MN468918 | .......................................................................................... |
| RM6715.d0616.A07 \| MN468919 | .......................................................................................... |
| RM6715.d0616.A09 \| MN468920 | .......................................................................................... |
| RM6715.d0616.A10 \| MN468921 | .......................................................................................... |
| RM6715.d0616.A11 \| MN468922 | .......................................................................................... |
| RM6715.d0616.A12 \| MN468923 | .......................................................................................... |
| RM6715.d0616.A14 \| MN468924 | ....................................................K..................................... |
| RM6715.d0616.A15 \| MN468925 | .......................................................................................... |
| RM6715.d0616.B02 \| MN468926 | .......................................................................................... |
| RM6715.d0616.B03 \| MN468927 | .......................................................................................... |
| RM6715.d0616.B04 \| MN468928 | .......................................................................................... |
| RM6715.d0616.B06 \| MN468929 | .......................................................................................... |
| RM6715.d0616.B07 \| MN468930 | .......................................................................................... |
| **RM6718 \| n = 27 \| 88 WPI** | |
| RM6718.d0616.A01 \| MN469235 | .................D........................................................................ |
| RM6718.d0616.A02 \| MN469236 | .................D........................................................................ |
| RM6718.d0616.A04 \| MN469237 | .................D........................................................................ |
| RM6718.d0616.A05 \| MN469238 | .................D........................................................................ |
| RM6718.d0616.A08 \| MN469240 | .................D........................................................................ |
| RM6718.d0616.A09 \| MN469241 | .................D........................................................................ |
| RM6718.d0616.A11 \| MN469242 | .................N........................................................................ |
| RM6718.d0616.A13 \| MN469243 | .................D........................................................................ |
| RM6718.d0616.A14 \| MN469244 | .................N........................................................................ |
| RM6718.d0616.A17 \| MN469246 | .................D........................................................................ |
| RM6718.d0616.A18 \| MN469247 | .................D........................................................................ |
| RM6718.d0616.A19 \| MN469248 | .................D........................................................................ |
| RM6718.d0616.A21 \| MN469249 | .................D........................................................................ |
| RM6718.d0616.A22 \| MN469250 | .................D........................................................................ |
| RM6718.d0616.A24 \| MN469251 | .................N........................................................................ |
| RM6718.d0616.A26 \| MN469252 | .................N........................................................................ |
| RM6718.d0616.A28 \| MN469254 | .................D........................................................................ |
| RM6718.d0616.A29 \| MN469255 | .................D..........Q............................................................. |
| RM6718.d0616.A30 \| MN469256 | .................D........................................................................ |
| RM6718.d0616.A31 \| MN469257 | .................D........................................................................ |
| RM6718.d0616.B01 \| MN469259 | .................D........................................................................ |
| RM6718.d0616.B04 \| MN469260 | .................N........................................................................ |
| RM6718.d0616.B05 \| MN469261 | .................D........................................................................ |
| RM6718.d0616.B09 \| MN469262 | .................D........................................................................ |
| RM6718.d0616.B10 \| MN469263 | .................D........................................................................ |
| RM6718.d0616.B11 \| MN469264 | .................N........................................................................ |
| RM6718.d0616.B13 \| MN469265 | .................D........................................................................ |
| **RM6719 \| n = 16 \| 88 WPI** | |
| RM6719.d0616.A01 \| MN469516 | .................N...K.................................................................... |
| RM6719.d0616.B01 \| MN469517 | .................N...K.................................................................... |
| RM6719.d0616.B03 \| MN469518 | .................N...K.................................................................... |
| RM6719.d0616.B04 \| MN469519 | .................N...K.................................................................... |
| RM6719.d0616.B05 \| MN469520 | .................N...K.................................................................... |
| RM6719.d0616.B06 \| MN469521 | .................N...K.................................................................... |
| RM6719.d0616.B07 \| MN469522 | .................N...K.................................................................... |
| RM6719.d0616.B09 \| MN469523 | .................N...K.................................................................... |
| RM6719.d0616.B11 \| MN469524 | .....................K.................................................................... |
| RM6719.d0616.B13 \| MN469525 | .................N...K.................................................................... |
| RM6719.d0616.B14 \| MN469526 | .....................K.................................................................... |
| RM6719.d0616.B15 \| MN469527 | .................N...K.................................................................... |
| RM6719.d0616.B16 \| MN469528 | .................N...K.................................................................... |
| RM6719.d0616.B17 \| MN469529 | .................N...K.................................................................... |
| RM6719.d0616.B18 \| MN469530 | .................N...K.................................................................... |
| RM6719.d0616.B19 \| MN469531 | .................N...K.................................................................... |

. identity    A substitution    A insertion    − deletion    − gap at insertion column    X unresolved residue

* in numbering: insertion column without a supplied HXB2 position.

Domain and position annotations follow HXB2 numbering. SP, signal peptide; Fusion, fusion peptide.

MPER, membrane−proximal external region. Each sequence is displayed separately; no identical sequences are collapsed.

### BG505 Env amino−acid alignment

6 macaques | 74 sampled sequences | reference coverage: HXB2 1−709

Segment 8/8 | HXB2 626−709 | alignment columns 631−714

**Sequence identifier | GenBank accession**

Env region: MPER (HXB2 662–683)

HXB2 numbering: 626 627 628 629 630 631 632 633 634 635 636 637 638 639 640 641 642 643 644 645 646 647 648 649 650 651 652 653 654 655 656 657 658 659 660 661 662 663 664 665 666 667 668 669 670 671 672 673 674 675 676 677 678 679 680 681 682 683 684 685 686 687 688 689 690 691 692 693 694 695 696 697 698 699 700 701 702 703 704 705 706 707 708 709

```
SHIV.A.BG505.T332N.375Y | WNK77136   MTWLQWDKEISNYTQIIYGLLEESQNQQEKNEQDLLALDKWASLWNWFDISNWLWYIKIFIMIVGGLIGLRIVFAVLSVIHRVR
```

**RM6708 | n = 3 | 88 WPI**

```
RM6708.d0616.A03 | MN468749   ........................................................T...........................
RM6708.d0616.A07 | MN468750   ........................................................T...........................
RM6708.d0616.A08 | MN468751   ........................................................T...........................
```

**RMT283 | n = 2 | 88 WPI**

```
RMT283.d0616.A01 | MN469591   ...............................................................................V....
RMT283.d0616.A02 | MN469592   ...............................................................................V....
```

**RM6706 | n = 10 | 88 WPI**

```
RM6706.d0616.A03 | MN468576   ....................................................................................
RM6706.d0616.A04 | MN468577   ...........................................................................A........
RM6706.d0616.A05 | MN468578   ...............................................................................V....
RM6706.d0616.A06 | MN468579   ....................................................................................
RM6706.d0616.A10 | MN468580   ....................................................................................
RM6706.d0616.A12 | MN468581   ....................................................................................
RM6706.d0616.A13 | MN468582   ....................................................................................
RM6706.d0616.A15 | MN468583   ..........................................G.........................................
RM6706.d0616.A16 | MN468584   ....................................................................................
RM6706.d0616.A18 | MN468585   .......................................R............................................
```

**RM6715 | n = 16 | 88 WPI**

```
RM6715.d0616.A01 | MN468914   .......................................................................V..T.........
RM6715.d0616.A02 | MN468915   ..........................................G...............................T.......A.
RM6715.d0616.A05 | MN468917   .........................................TG...............................T.........
RM6715.d0616.A06 | MN468918   ..........................................................................T.......A.
RM6715.d0616.A07 | MN468919   .......................................................................V..T.........
RM6715.d0616.A09 | MN468920   ..........................................G...............................T.........
RM6715.d0616.A10 | MN468921   ..........................................G...............................T.........
RM6715.d0616.A11 | MN468922   ..........................................G...............................T.........
RM6715.d0616.A12 | MN468923   ..........................................................................T.......A.
RM6715.d0616.A14 | MN468924   ....................................................................E.....T.......A.
RM6715.d0616.A15 | MN468925   ..........................................................................T.......A.
RM6715.d0616.B02 | MN468926   ..........................................G...............................T.........
RM6715.d0616.B03 | MN468927   ..........................................G...............................T.........
RM6715.d0616.B04 | MN468928   .......................................................................V..T.........
RM6715.d0616.B06 | MN468929   ..........................................................................T.......A.
RM6715.d0616.B07 | MN468930   ..........................................................................T.........
```

**RM6718 | n = 27 | 88 WPI**

```
RM6718.d0616.A01 | MN469235   ....................................................................................
RM6718.d0616.A02 | MN469236   ....................................................................................
RM6718.d0616.A04 | MN469237   ....................................................................................
RM6718.d0616.A05 | MN469238   ....................................................................................
RM6718.d0616.A08 | MN469240   ..........................................N.........................................
RM6718.d0616.A09 | MN469241   ....................................................................................
RM6718.d0616.A11 | MN469242   ....................................................................................
RM6718.d0616.A13 | MN469243   ....................................................................................
RM6718.d0616.A14 | MN469244   ....................................................................................
RM6718.d0616.A17 | MN469246   ....................................................................................
RM6718.d0616.A18 | MN469247   ....................................................................................
RM6718.d0616.A19 | MN469248   ....................................................................................
RM6718.d0616.A21 | MN469249   ....................................................................................
RM6718.d0616.A22 | MN469250   ....................................................................................
RM6718.d0616.A24 | MN469251   ..................E..........................S......................................
RM6718.d0616.A26 | MN469252   ..........................................G.........................................
RM6718.d0616.A28 | MN469254   ....................................................................................
RM6718.d0616.A29 | MN469255   ................................................E...................................
RM6718.d0616.A30 | MN469256   ....................................................................................
RM6718.d0616.A31 | MN469257   ....................................................................................
RM6718.d0616.B01 | MN469259   ....................................................................................
RM6718.d0616.B04 | MN469260   ....................................................................................
RM6718.d0616.B05 | MN469261   ....................................................................................
RM6718.d0616.B09 | MN469262   .................................................................................K..
RM6718.d0616.B10 | MN469263   ....................................................................................
RM6718.d0616.B11 | MN469264   ....................................................................................
RM6718.d0616.B13 | MN469265   ....................................................................................
```

**RM6719 | n = 16 | 88 WPI**

```
RM6719.d0616.A01 | MN469516   ....................................................................................
RM6719.d0616.B01 | MN469517   ....................................................................................
RM6719.d0616.B03 | MN469518   .......................................R............................................
RM6719.d0616.B04 | MN469519   ....................................................................................
RM6719.d0616.B05 | MN469520   ....................................................................................
RM6719.d0616.B06 | MN469521   ....................................................................................
RM6719.d0616.B07 | MN469522   ....................................................................................
RM6719.d0616.B09 | MN469523   ....................................................................................
RM6719.d0616.B11 | MN469524   ....................................................................................
RM6719.d0616.B13 | MN469525   ....................................................................................
RM6719.d0616.B14 | MN469526   ....................................................................................
RM6719.d0616.B15 | MN469527   ....................................................................................
RM6719.d0616.B16 | MN469528   ....................................................................................
RM6719.d0616.B17 | MN469529   ....................................................................................
RM6719.d0616.B18 | MN469530   ....................................................................................
RM6719.d0616.B19 | MN469531   ....................................................................................
```

. identity  A substitution  A insertion  − deletion  − gap at insertion column  X unresolved residue

* in numbering: insertion column without a supplied HXB2 position.

Domain and position annotations follow HXB2 numbering. SP, signal peptide; Fusion, fusion peptide.

MPER, membrane−proximal external region. Each sequence is displayed separately; no identical sequences are collapsed.
