## Supplementary material for "Functional Convergence of Genetically Diverse B-Cell Receptors in Simian-HIV Infected Rhesus Macaques": Document S2

### CH505 Env amino−acid alignment

6 macaques | 105 sampled sequences | reference coverage: HXB2 1−709

Segment 1/8 | HXB2 1−94 | alignment columns 1−90

**Sequence identifier | GenBank accession**

Env region: SP (HXB2 1–30) | C1 (HXB2 34–94)

HXB2 numbering: 1 2 3 4 5 6 7 8 9 10 11 12 13 15 16 17 18 19 20 21 22 23 24 25 26 27 28 29 30 34 35 36 37 38 39 40 41 42 43 44 45 46 47 48 49 50 51 52 53 54 55 56 57 58 59 60 61 62 63 64 65 66 67 68 69 70 71 72 73 74 75 76 77 78 79 80 81 82 83 84 85 86 87 88 89 90 91 92 93 94

| Sequence identifier \| GenBank accession | Alignment |
|---|---|
| **SHIV.CH505.375H \| QSQ65056** | MRVMGIQRNYPQWWIWSMLGFWMLMICNGMWVTVYYGVPVWKEAKTTLFCASDAKAYEKEVHNVWATHACVPTDPNPQEMVLKNVTENFN |
| **RM6697 \| n = 38 \| 56 WPI** | |
| RM6697_w056_A02 \| MT486140 | ..........L..........L.................................................................D.. |
| RM6697_w056_A03 \| MT486141 | ..........L..........L..........................................................I......... |
| RM6697_w056_A04 \| MT486142 | ..........L..........L.................................................................... |
| RM6697_w056_A05 \| MT486143 | ..........L..........L.................................................................... |
| RM6697_w056_A06 \| MT486144 | ..........L..........L.................................................................... |
| RM6697_w056_A07 \| MT486145 | ..........L..........L.................................................................... |
| RM6697_w056_A09 \| MT486147 | ..........L..........L.................................................................... |
| RM6697_w056_A11 \| MT486148 | ..........L..........L.................................................................... |
| RM6697_w056_A12 \| MT486149 | ..........L..........L.................................................................... |
| RM6697_w056_A13 \| MT486150 | ..........L..........L.................................................................... |
| RM6697_w056_A14 \| MT486151 | ..........L..........L.................................................................... |
| RM6697_w056_A15 \| MT486152 | ..........L..........L.................................................................... |
| RM6697_w056_A16 \| MT486153 | ..........L..........L.................................................................... |
| RM6697_w056_A17 \| MT486154 | ..........L..........L.................................................................... |
| RM6697_w056_A19 \| MT486155 | ..........L..........L.................................................................... |
| RM6697_w056_A22 \| MT486156 | ..........L............................................................................... |
| RM6697_w056_A23 \| MT486157 | ..........L............................................................................... |
| RM6697_w056_A24 \| MT486158 | ..........L..........L.................................................................... |
| RM6697_w056_A25 \| MT486159 | ..........L..........L.................................................................... |
| RM6697_w056_A27 \| MT486160 | ..........L..........L.................................................................D.. |
| RM6697_w056_A30 \| MT486161 | ..........L..........L.................................................................... |
| RM6697_w056_A31 \| MT486162 | ..........L..........L.................................................................... |
| RM6697_w056_B01 \| MT486164 | ..........L..........L.................................................................... |
| RM6697_w056_B03 \| MT486165 | ..........L..........L.................................................................... |
| RM6697_w056_B04 \| MT486166 | ..........L..........L.................................................................... |
| RM6697_w056_B06 \| MT486167 | ..........L..........L.................................................................... |
| RM6697_w056_B07 \| MT486168 | ..........L...........R................................................................... |
| RM6697_w056_B08 \| MT486169 | ..........L............................................................................... |
| RM6697_w056_B09 \| MT486170 | ..........L..........L.........I.......................................................... |
| RM6697_w056_B11 \| MT486171 | ..........L..........L.................................................................... |
| RM6697_w056_B12 \| MT486172 | ..........L..........L.................................................................... |
| RM6697_w056_B17 \| MT486174 | ..........L..........L.................................................................D.. |
| RM6697_w056_B19 \| MT486175 | ..........L..........L.................................................................... |
| RM6697_w056_B20 \| MT486176 | ..........L..........L.................................................................... |
| RM6697_w056_B21 \| MT486177 | ..........L..........L.................................................................... |
| RM6697_w056_B22 \| MT486178 | ..........L..........L.................................................................... |
| RM6697_w056_T01 \| MT486179 | ..........L..........L.................................................................... |
| RM6697_w056_T04 \| MT486180 | ..........L..........L...N................................................................ |
| **RM6698 \| n = 3 \| 56 WPI** | |
| RM6698_w056_B01 \| MT486279 | ...T.................L.................................................................... |
| RM6698_w056_B03 \| MT486281 | ...T.................L.................................................................... |
| RM6698_w056_T01 \| MT486282 | .......................................................................................... |
| **RM6699 \| n = 9 \| 56 WPI** | |
| RM6699_w056_B01 \| MT486349 | .......................................................................................... |
| RM6699_w056_B04 \| MT486351 | .......................................................................................... |
| RM6699_w056_B05 \| MT486352 | .......................................................................................... |
| RM6699_w056_B06 \| MT486353 | .......................................................................................... |
| RM6699_w056_B07 \| MT486354 | .......................................................................................... |
| RM6699_w056_T03 \| MT486355 | .......................................................................................... |
| RM6699_w056_T04 \| MT486356 | .......................................................................................... |
| RM6699_w056_T05 \| MT486357 | .......................................................................................... |
| RM6699_w056_T06 \| MT486358 | .......................................................................................... |
| **RM6701 \| n = 18 \| 56 WPI** | |
| RM6701_w056_A04 \| MT486603 | ..................................H....................................................... |
| RM6701_w056_A06 \| MT486604 | .......................................................................................... |
| RM6701_w056_A08 \| MT486605 | .......................................................................................... |
| RM6701_w056_A10 \| MT486607 | .......................................................................................... |
| RM6701_w056_A12 \| MT486608 | .......................................................................................... |
| RM6701_w056_A15 \| MT486609 | ..................................C...................................................K... |
| RM6701_w056_A16 \| MT486610 | .......................................................................................... |
| RM6701_w056_A19 \| MT486612 | .......................................................................................... |
| RM6701_w056_A20 \| MT486613 | .......................................................................................... |
| RM6701_w056_A22 \| MT486615 | .......................................................................................... |
| RM6701_w056_B01 \| MT486616 | ..M....................................................................................... |
| RM6701_w056_B03 \| MT486618 | .......................................................................................... |
| RM6701_w056_B04 \| MT486619 | .......................................................................................... |
| RM6701_w056_B05 \| MT486620 | .......................................................................................... |
| RM6701_w056_B06 \| MT486621 | .......................................................................................... |
| RM6701_w056_T01 \| MT486622 | .......................................................................................... |
| RM6701_w056_T02 \| MT486623 | .......................................................................................... |
| RM6701_w056_T04 \| MT486624 | .......................................................................................... |
| **RM6702 \| n = 17 \| 24 WPI** | |
| RM6702_w024_A01 \| MT486654 | .......................................................................................... |
| RM6702_w024_A02 \| MT486655 | .......................................................................................... |
| RM6702_w024_A03 \| MT486656 | .......................................................................................... |
| RM6702_w024_A04 \| MT486657 | .......................................................................................... |
| RM6702_w024_A05 \| MT486658 | .......................................................................................... |
| RM6702_w024_A08 \| MT486660 | .......................................................................................... |
| RM6702_w024_A10 \| MT486661 | .......................................................................................... |
| RM6702_w024_A13 \| MT486663 | .......................................................................................... |
| RM6702_w024_A14 \| MT486664 | .......................................................................................... |
| RM6702_w024_A15 \| MT486665 | .......................................................................................... |
| RM6702_w024_A16 \| MT486666 | .......................................................................................... |
| RM6702_w024_A17 \| MT486667 | .......................................................................................... |
| RM6702_w024_A18 \| MT486668 | .......................................................................................... |
| RM6702_w024_A19 \| MT486669 | .......................................................................................... |
| RM6702_w024_A20 \| MT486670 | .......................................................................................... |
| RM6702_w024_T01 \| MT486671 | .......................................................................................... |
| RM6702_w024_T02 \| MT486672 | .......................................................................................... |
| **RM6703 \| n = 20 \| 48 WPI** | |
| RM6703_w048_A03 \| MT486741 | .....................................................T.................................... |
| RM6703_w048_A04 \| MT486742 | .......................................................................................... |
| RM6703_w048_A05 \| MT486743 | .......................................................................................... |
| RM6703_w048_A06 \| MT486744 | .......................................................................................... |
| RM6703_w048_A08 \| MT486746 | .......................................................T.................................. |
| RM6703_w048_A11 \| MT486747 | .......................................................................................... |
| RM6703_w048_A13 \| MT486748 | .......................................................................................... |
| RM6703_w048_A15 \| MT486749 | .......................................................................................... |
| RM6703_w048_B01 \| MT486750 | ..............T........................................................................... |
| RM6703_w048_B04 \| MT486752 | .......................................................................................... |
| RM6703_w048_B06 \| MT486753 | .......................................................................................... |
| RM6703_w048_B07 \| MT486754 | .......................................................................................... |
| RM6703_w048_B08 \| MT486755 | .......................................................................................... |
| RM6703_w048_T01 \| MT486756 | .......................................................................................... |
| RM6703_w048_T02 \| MT486757 | .......................................................................................... |
| RM6703_w048_T03 \| MT486758 | .......................................................................................... |
| RM6703_w048_T06 \| MT486759 | .......................................................................................... |
| RM6703_w048_T07 \| MT486760 | .......................................................................................... |
| RM6703_w048_T08 \| MT486761 | .......................................................................................... |
| RM6703_w048_T09 \| MT486762 | .......................................................................................... |

. identity    A substitution    A insertion    − deletion    − gap at insertion column    X unresolved residue

* in numbering: insertion column without a supplied HXB2 position.

Domain and position annotations follow HXB2 numbering. SP, signal peptide; Fusion, fusion peptide.

MPER, membrane−proximal external region. Each sequence is displayed separately; no identical sequences are collapsed.

### CH505 Env amino−acid alignment

6 macaques | 105 sampled sequences | reference coverage: HXB2 1−709

Segment 2/8 | HXB2 95−182 | alignment columns 91−180

**Sequence identifier | GenBank accession**

Env region: C1 (HXB2 95–132) | V1 (HXB2 136–156) | V2 (HXB2 157–182)

HXB2 numbering: 95 96 97 98 99 100 101 102 103 104 105 106 107 108 109 110 111 112 113 114 115 116 117 118 119 120 121 122 123 124 125 126 127 128 129 130 131 132 136 137 138 * * * * * * * * * 139 140 * * 141 142 143 148 149 150 153 154 155 156 157 158 159 160 161 162 163 164 165 166 167 168 169 170 171 172 173 174 175 176 177 178 179 180 181 182

```
SHIV.CH505.375H | QSQ65056   MWKNDMVDQMHEDVISLWDQSLKPCVKLTPLCVTLNCTNAT---------AS--NSSIIEGMKNCSFNITTELRDKREKKNALFYKLDIV
```

**RM6697 | n = 38 | 56 WPI**

```
RM6697_w056_A02 | MT486140   ...................................D.....---------.G--....................................
RM6697_w056_A03 | MT486141   .....................................A...---------.G--....................................
RM6697_w056_A04 | MT486142   .........................................-ASSSTNAT..--....................................
RM6697_w056_A05 | MT486143   .........................................-ASSSTNAT..--................................F...
RM6697_w056_A06 | MT486144   .........................................-ASSSTNAT..--....................................
RM6697_w056_A07 | MT486145   .........................................-ASSSTNAT..--....................................
RM6697_w056_A09 | MT486147   .....................................A...-TSSGTNAT..--....................................
RM6697_w056_A11 | MT486148   ......................R..................-ASNSTNDTV.--....................................
RM6697_w056_A12 | MT486149   .........................................-ASSSTNAT..--....................................
RM6697_w056_A13 | MT486150   .........................................-ASNSTNAT..--....................................
RM6697_w056_A14 | MT486151   .........................................-ASSSTNAT..--....................................
RM6697_w056_A15 | MT486152   .........................................-ASSSTNAT..--....................................
RM6697_w056_A16 | MT486153   ...................................D.....-ASSSTNAT..--.....................R..............
RM6697_w056_A17 | MT486154   .........................................-ASNSTNAT..--....................................
RM6697_w056_A19 | MT486155   .........................................-ASSSTNAT..--....................................
RM6697_w056_A22 | MT486156   ...................................D.....-ASSSTNAT..--....................................
RM6697_w056_A23 | MT486157   ...................................D.....-ASSSTNAT..--....................................
RM6697_w056_A24 | MT486158   .........................................-ASSSTNAT.G--....................................
RM6697_w056_A25 | MT486159   .........................................-AGNSTNAT.G--....................................
RM6697_w056_A27 | MT486160   ...................................D.....---------.G--....................................
RM6697_w056_A30 | MT486161   ...................................D.....-ASSSTNAT..--....................................
RM6697_w056_A31 | MT486162   .........................................-ASSSTNAT..--....................................
RM6697_w056_B01 | MT486164   .........................................-ASNSTNAT..--....................................
RM6697_w056_B03 | MT486165   .........................................-ASSSTNAT..--....................................
RM6697_w056_B04 | MT486166   .........................................-ASNSTNAT..--....................................
RM6697_w056_B06 | MT486167   ...................................D.....-ASSSTNATV.--....................................
RM6697_w056_B07 | MT486168   ...................................D.....-ASSSTNAT..--....................................
RM6697_w056_B08 | MT486169   ...................................D.....-ASSSTNAT..--....................................
RM6697_w056_B09 | MT486170   ......M..................................-ASSSTNAT..--....................................
RM6697_w056_B11 | MT486171   .........................................-ASSSTNAT..--....................................
RM6697_w056_B12 | MT486172   .....................................I...-ASSSTNAT..--....................................
RM6697_w056_B17 | MT486174   ...................................D.....---------.G--....................................
RM6697_w056_B19 | MT486175   .........................................-ASSSTNAT..--....................................
RM6697_w056_B20 | MT486176   ...................................T.....-ASSSTNAT..--....................................
RM6697_w056_B21 | MT486177   ...................................D.....-ASSSTNAT..--....................................
RM6697_w056_B22 | MT486178   ...................................D.....-ASSSTNATV.--....................................
RM6697_w056_T01 | MT486179   ...................................D.....---------.N--.N...................R..............
RM6697_w056_T04 | MT486180   ......................R..................-ASNSTNAT..--....................................
```

**RM6698 | n = 3 | 56 WPI**

```
RM6698_w056_B01 | MT486279   .........................................DSNSSIIEGMK--....................................
RM6698_w056_B03 | MT486281   .........................................DSNSSIIEGMK--....................................
RM6698_w056_T01 | MT486282   .........................................--ASNSSIT..--....................................
```

**RM6699 | n = 9 | 56 WPI**

```
RM6699_w056_B01 | MT486349   .........................................---------VN--....................................
RM6699_w056_B04 | MT486351   .........................................---------T.--........R...........................
RM6699_w056_B05 | MT486352   .........................................---------T.--........R...........................
RM6699_w056_B06 | MT486353   .........................................---------T.--........R...........................
RM6699_w056_B07 | MT486354   ..........................R..............---------.R--....................................
RM6699_w056_T03 | MT486355   ..........................R..............---------.R--....................................
RM6699_w056_T04 | MT486356   .........................................---------.N--....M...............................
RM6699_w056_T05 | MT486357   .........................................---------T.--........R...........................
RM6699_w056_T06 | MT486358   ..........................R..............---------.R--....................................
```

**RM6701 | n = 18 | 56 WPI**

```
RM6701_w056_A04 | MT486603   ...............G.........................---------..--.....................R..............
RM6701_w056_A06 | MT486604   .........................................---------..--....................................
RM6701_w056_A08 | MT486605   ...............G.........................---------..--.....................R..............
RM6701_w056_A10 | MT486607   ...............G.........................---------..--.....................R..............
RM6701_w056_A12 | MT486608   ...............G.........................---------..--....................................
RM6701_w056_A15 | MT486609   ...............G.........................---------..--.....................R..............
RM6701_w056_A16 | MT486610   ...............G.........................---------..--.....................R..............
RM6701_w056_A19 | MT486612   ...............G.........................---------..--.....................R..............
RM6701_w056_A20 | MT486613   ...............G.........................---------..--.....................R..............
RM6701_w056_A22 | MT486615   ...............G.........................---------..--.....................R..............
RM6701_w056_B01 | MT486616   ...............G.........................---------..--.....................R..............
RM6701_w056_B03 | MT486618   ...............G.........................---------..--.....................R..............
RM6701_w056_B04 | MT486619   ...............G.........................---------..--.....................R..............
RM6701_w056_B05 | MT486620   ...............G.........................---------..--....................................
RM6701_w056_B06 | MT486621   ...............G.........................---------..--.....................R.K............
RM6701_w056_T01 | MT486622   ...............G.........................---------..--.....................R..............
RM6701_w056_T02 | MT486623   ...............G.........................---------..--.....................R..............
RM6701_w056_T04 | MT486624   .........................................---------..--....................................
```

**RM6702 | n = 17 | 24 WPI**

```
RM6702_w024_A01 | MT486654   .........................................---------..--...-................................
RM6702_w024_A02 | MT486655   .........................................---------..--...-................................
RM6702_w024_A03 | MT486656   .........................................---------..--...-................................
RM6702_w024_A04 | MT486657   .........................................---------..--...-................................
RM6702_w024_A05 | MT486658   .........................................---------..--...-................................
RM6702_w024_A08 | MT486660   .........................................---------..--...-................................
RM6702_w024_A10 | MT486661   .........................................---------..--...-................................
RM6702_w024_A13 | MT486663   .........................................---------..--...-................................
RM6702_w024_A14 | MT486664   .........................................---------..--...-................................
RM6702_w024_A15 | MT486665   .........................................---------..--...-................................
RM6702_w024_A16 | MT486666   .........................................---------..--...-................................
RM6702_w024_A17 | MT486667   .........................................---------..--..G-................................
RM6702_w024_A18 | MT486668   .........................................---------..--...-.........D......................
RM6702_w024_A19 | MT486669   .........................................---------..--...-................................
RM6702_w024_A20 | MT486670   ..............................................................-A.K................................
RM6702_w024_T01 | MT486671   .........................................---------..--...-................................
RM6702_w024_T02 | MT486672   .........................................---------..--...-................................
```

**RM6703 | n = 20 | 48 WPI**

```
RM6703_w048_A03 | MT486741   .......................................D.---------..--...-................................
RM6703_w048_A04 | MT486742   .........................................-----ANAT..--....................................
RM6703_w048_A05 | MT486743   .........................................---------..--...-.........D......................
RM6703_w048_A06 | MT486744   .........................................-----ANAT..--....................................
RM6703_w048_A08 | MT486746   .........................................---ASTNAT..--...-................................
RM6703_w048_A11 | MT486747   ...................................D.....---------V.--...-................................
RM6703_w048_A13 | MT486748   .........................................-----ANAT..--.......I............................
RM6703_w048_A15 | MT486749   ...................................D.....-----A--SN.--S..........................T........
RM6703_w048_B01 | MT486750   ...................................D.....---------V.--...-................................
RM6703_w048_B04 | MT486752   .........................................-----ANAT..--....................................
RM6703_w048_B06 | MT486753   .........................................---------V.--...-................................
RM6703_w048_B07 | MT486754   .........................................-----ANAT..--....................................
RM6703_w048_B08 | MT486755   .........................................---------V.--...-V...............................
RM6703_w048_T01 | MT486756   .........................................---------..--...-.............K..................
RM6703_w048_T02 | MT486757   ...................................D.....---------..--...-................................
RM6703_w048_T03 | MT486758   .........................................-----ANAT..--....................................
RM6703_w048_T06 | MT486759   .........................................---------..--.---................................
RM6703_w048_T07 | MT486760   .........................................---------V.--...-V...............................
RM6703_w048_T08 | MT486761   .....................................A...ASNSSLIEGMKNS....................................
RM6703_w048_T09 | MT486762   .........................................---------..--...-................................
```

. identity | A substitution | A insertion | − deletion | − gap at insertion column | X unresolved residue

* in numbering: insertion column without a supplied HXB2 position.

Domain and position annotations follow HXB2 numbering. SP, signal peptide; Fusion, fusion peptide.

MPER, membrane−proximal external region. Each sequence is displayed separately; no identical sequences are collapsed.

### CH505 Env amino−acid alignment

6 macaques | 105 sampled sequences | reference coverage: HXB2 1−709

Segment 3/8 | HXB2 183−272 | alignment columns 181−270

**Sequence identifier | GenBank accession**

Env region: V2 | C2

HXB2 numbering: 183 184 185 186 187 188 189 190 191 192 193 194 195 196 197 198 199 200 201 202 203 204 205 206 207 208 209 210 211 212 213 214 215 216 217 218 219 220 221 222 223 224 225 226 227 228 229 230 231 232 233 234 235 236 237 238 239 240 241 242 243 244 245 246 247 248 249 250 251 252 253 254 255 256 257 258 259 260 261 262 263 264 265 266 267 268 269 270 271 272

| Sequence identifier | GenBank accession | Alignment |
|---|---|---|
| **SHIV.CH505.375H** | **QSQ65056** | QLDGNSSQYRLINCNTSVITQACPKVSFDPIPIHYCAPAGYAILKCNNKTFTGTGPCNNVSTVQCTHGIKPVVSTQLLLNGSLAEGEIII |
| **RM6697 \| n = 38 \| 56 WPI** | | |
| RM6697_w056_A02 | MT486140 | .................I........................................................................ |
| RM6697_w056_A03 | MT486141 | .................I.............................D...N...................................... |
| RM6697_w056_A04 | MT486142 | .................I..............................E......................................... |
| RM6697_w056_A05 | MT486143 | .................I........................................................................ |
| RM6697_w056_A06 | MT486144 | .................I........................................................................ |
| RM6697_w056_A07 | MT486145 | .................I.............................D...N....................I................. |
| RM6697_w056_A09 | MT486147 | .................I........................................................................ |
| RM6697_w056_A11 | MT486148 | .................I.............................D.......................................... |
| RM6697_w056_A12 | MT486149 | .................I...................................I.................................... |
| RM6697_w056_A13 | MT486150 | .................I........................................................................ |
| RM6697_w056_A14 | MT486151 | .................I.............................D...N...................................... |
| RM6697_w056_A15 | MT486152 | .................I........................................................................ |
| RM6697_w056_A16 | MT486153 | .................I........................................................................ |
| RM6697_w056_A17 | MT486154 | .................I........................................................................ |
| RM6697_w056_A19 | MT486155 | ..............D..I........................................................................ |
| RM6697_w056_A22 | MT486156 | .................I........................................................................ |
| RM6697_w056_A23 | MT486157 | .................I........................................................................ |
| RM6697_w056_A24 | MT486158 | .................I........................................................................ |
| RM6697_w056_A25 | MT486159 | .................I........................................................................ |
| RM6697_w056_A27 | MT486160 | .................I........................................................................ |
| RM6697_w056_A30 | MT486161 | .................I........................................................................ |
| RM6697_w056_A31 | MT486162 | .................I........................................................................ |
| RM6697_w056_B01 | MT486164 | .................I.............................D...N...................................... |
| RM6697_w056_B03 | MT486165 | .................I........................................................................ |
| RM6697_w056_B04 | MT486166 | .................I........................................................................ |
| RM6697_w056_B06 | MT486167 | .................I........................................................................ |
| RM6697_w056_B07 | MT486168 | .................I........................................................................ |
| RM6697_w056_B08 | MT486169 | .................I........................................................................ |
| RM6697_w056_B09 | MT486170 | .................I.............................D...N...................................... |
| RM6697_w056_B11 | MT486171 | .................I........................................................................ |
| RM6697_w056_B12 | MT486172 | .................I........................................................................ |
| RM6697_w056_B17 | MT486174 | .................I........................................................................ |
| RM6697_w056_B19 | MT486175 | .................I........................................................................ |
| RM6697_w056_B20 | MT486176 | .................I.................................N....................I................. |
| RM6697_w056_B21 | MT486177 | .................I........................................................................ |
| RM6697_w056_B22 | MT486178 | .................I........................................................................ |
| RM6697_w056_T01 | MT486179 | .................I........................................................................ |
| RM6697_w056_T04 | MT486180 | .................I.............................D...N...................................... |
| **RM6698 \| n = 3 \| 56 WPI** | | |
| RM6698_w056_B01 | MT486279 | .....................................................................................E.... |
| RM6698_w056_B03 | MT486281 | .....................................................................................E.... |
| RM6698_w056_T01 | MT486282 | .......................................................................................... |
| **RM6699 \| n = 9 \| 56 WPI** | | |
| RM6699_w056_B01 | MT486349 | ..................M............................D.......................................... |
| RM6699_w056_B04 | MT486351 | ...............................................D.....................R.................... |
| RM6699_w056_B05 | MT486352 | ...............................................D.....................R.................... |
| RM6699_w056_B06 | MT486353 | ...............................................D.....................R.................... |
| RM6699_w056_B07 | MT486354 | ...S...........................................D.....................R.................... |
| RM6699_w056_T03 | MT486355 | ...............................................D.......................................... |
| RM6699_w056_T04 | MT486356 | ...............................................S.......................................... |
| RM6699_w056_T05 | MT486357 | ...............................................D.....................R.................... |
| RM6699_w056_T06 | MT486358 | ...............................................D.......................................... |
| **RM6701 \| n = 18 \| 56 WPI** | | |
| RM6701_w056_A04 | MT486603 | .......................................................................................... |
| RM6701_w056_A06 | MT486604 | .....................................................I.................................... |
| RM6701_w056_A08 | MT486605 | .......................................................................................... |
| RM6701_w056_A10 | MT486607 | .......................................................................................... |
| RM6701_w056_A12 | MT486608 | .......................................................................................... |
| RM6701_w056_A15 | MT486609 | .......................................................................................... |
| RM6701_w056_A16 | MT486610 | .......................................................................................... |
| RM6701_w056_A19 | MT486612 | .......................................................................................... |
| RM6701_w056_A20 | MT486613 | .......................................................................................... |
| RM6701_w056_A22 | MT486615 | .......................................................................................... |
| RM6701_w056_B01 | MT486616 | .......................................................................................... |
| RM6701_w056_B03 | MT486618 | .......................................................................................... |
| RM6701_w056_B04 | MT486619 | .......................................................................................... |
| RM6701_w056_B05 | MT486620 | .....................................................I.................................... |
| RM6701_w056_B06 | MT486621 | .......................................................................................... |
| RM6701_w056_T01 | MT486622 | .......................................................................................... |
| RM6701_w056_T02 | MT486623 | .......................................................................................... |
| RM6701_w056_T04 | MT486624 | .....................................................I.................................... |
| **RM6702 \| n = 17 \| 24 WPI** | | |
| RM6702_w024_A01 | MT486654 | .......................................................................................... |
| RM6702_w024_A02 | MT486655 | .......................................................................................... |
| RM6702_w024_A03 | MT486656 | .......................................................................................... |
| RM6702_w024_A04 | MT486657 | .......................................................................................... |
| RM6702_w024_A05 | MT486658 | .......................................................................................... |
| RM6702_w024_A08 | MT486660 | ...D...................................................................................... |
| RM6702_w024_A10 | MT486661 | .......................................................................................... |
| RM6702_w024_A13 | MT486663 | .......................................................................................... |
| RM6702_w024_A14 | MT486664 | ...D...................................................................................... |
| RM6702_w024_A15 | MT486665 | .......................................................................................... |
| RM6702_w024_A16 | MT486666 | ...D...................................................................................... |
| RM6702_w024_A17 | MT486667 | .......................................................................................... |
| RM6702_w024_A18 | MT486668 | .......................................................................................... |
| RM6702_w024_A19 | MT486669 | .......................................................................................... |
| RM6702_w024_A20 | MT486670 | .......................................................................................... |
| RM6702_w024_T01 | MT486671 | .......................................................................................... |
| RM6702_w024_T02 | MT486672 | .......................................................................................... |
| **RM6703 \| n = 20 \| 48 WPI** | | |
| RM6703_w048_A03 | MT486741 | .......................................................................................... |
| RM6703_w048_A04 | MT486742 | .......................................................................................... |
| RM6703_w048_A05 | MT486743 | ..................V....................................................................... |
| RM6703_w048_A06 | MT486744 | ............................................................I............................. |
| RM6703_w048_A08 | MT486746 | .......................................................................................... |
| RM6703_w048_A11 | MT486747 | .......................................................................................... |
| RM6703_w048_A13 | MT486748 | .......................................................................................... |
| RM6703_w048_A15 | MT486749 | .......................................................................................... |
| RM6703_w048_B01 | MT486750 | .......................................................................................... |
| RM6703_w048_B04 | MT486752 | .......................................................................................... |
| RM6703_w048_B06 | MT486753 | .......................................................................................... |
| RM6703_w048_B07 | MT486754 | .......................................................................................... |
| RM6703_w048_B08 | MT486755 | .......................................................................................... |
| RM6703_w048_T01 | MT486756 | .......................................................................................... |
| RM6703_w048_T02 | MT486757 | .......................................................................................... |
| RM6703_w048_T03 | MT486758 | .......................................................................................... |
| RM6703_w048_T06 | MT486759 | .......................................................................................... |
| RM6703_w048_T07 | MT486760 | ....................................................................................K..... |
| RM6703_w048_T08 | MT486761 | .......................................................................................... |
| RM6703_w048_T09 | MT486762 | .......................................................................................... |

. identity    A substitution    A insertion    − deletion    − gap at insertion column    X unresolved residue

* in numbering: insertion column without a supplied HXB2 position.

Domain and position annotations follow HXB2 numbering. SP, signal peptide; Fusion, fusion peptide.

MPER, membrane−proximal external region. Each sequence is displayed separately; no identical sequences are collapsed.

### CH505 Env amino−acid alignment

6 macaques | 105 sampled sequences | reference coverage: HXB2 1−709

Segment 4/8 | HXB2 273−364 | alignment columns 271−360

**Sequence identifier | GenBank accession**

Env region: C2 (HXB2 273–295) | V3 (HXB2 296–330) | C3 (HXB2 331–364)

HXB2 numbering: 273 274 275 276 277 278 279 280 281 282 283 284 285 286 287 288 289 290 291 292 293 294 295 296 297 298 299 300 301 302 303 304 305 306 307 308 309 312 313 314 315 316 317 318 319 320 321 322 322a 323 324 325 326 327 328 329 330 331 332 333 334 335 336 337 338 339 340 341 342 343 344 345 346 347 348 349 350 351 352 353 355 356 357 358 359 360 361 362 363 364

| Sequence identifier | GenBank accession | Alignment |
|---|---|---|
| **SHIV.CH505.375H** | **QSQ65056** | RSENITNNVKTIIVHLNESVKIECTRPNNKTRTSIRIGPGQAFYATGQVIGDIREAYCNINESKWNETLQRVSKKLKEYFPHKNITFQPS |
| **RM6697 \| n = 38 \| 56 WPI** | | |
| RM6697_w056_A02 | MT486140 | .............................N..........................H...S..........................N........ |
| RM6697_w056_A03 | MT486141 | .............................N...........T..............H...S...............E.....N........ |
| RM6697_w056_A04 | MT486142 | .............................N..........................H...S.................H..N........ |
| RM6697_w056_A05 | MT486143 | .............................N..........................H...S...............E.....N........ |
| RM6697_w056_A06 | MT486144 | .............................N..........................H...S..........................N........ |
| RM6697_w056_A07 | MT486145 | .............................N..........................H...S......I........EG............ |
| RM6697_w056_A09 | MT486147 | .............................N..........................H...S.................H..N........ |
| RM6697_w056_A11 | MT486148 | .............................N..........................H...S.................H.L......... |
| RM6697_w056_A12 | MT486149 | .............................N..........................H...S................G...N........ |
| RM6697_w056_A13 | MT486150 | .............................N..........................H...S..........................N........ |
| RM6697_w056_A14 | MT486151 | .............................N..........................H...S...............E.....Y........ |
| RM6697_w056_A15 | MT486152 | .............................N..........................H...S...............E.....N........ |
| RM6697_w056_A16 | MT486153 | .............................N..........................H...S................G...N........ |
| RM6697_w056_A17 | MT486154 | .............................N...............I..........H...S..........................N........ |
| RM6697_w056_A19 | MT486155 | .............................N..........................H...S................K...N........ |
| RM6697_w056_A22 | MT486156 | .............................N..........................H...S...............E.....N........ |
| RM6697_w056_A23 | MT486157 | .............................N..........................H...S................G...N........ |
| RM6697_w056_A24 | MT486158 | .............................N..........................H...S..........................N........ |
| RM6697_w056_A25 | MT486159 | .............................N..........................H...S...............EK............ |
| RM6697_w056_A27 | MT486160 | .............................N..........................H...S................G...N........ |
| RM6697_w056_A30 | MT486161 | .............................N..........................H...S..........................N........ |
| RM6697_w056_A31 | MT486162 | .............................N..........................H...S..........................N........ |
| RM6697_w056_B01 | MT486164 | .............................N..........................H...S.................H.L......... |
| RM6697_w056_B03 | MT486165 | .............................N..........................H...S..........................N........ |
| RM6697_w056_B04 | MT486166 | .............................N..........................H...S..........................N........ |
| RM6697_w056_B06 | MT486167 | .............................N..........................H...S..........................N........ |
| RM6697_w056_B07 | MT486168 | .............................N..........................H...S................G...N........ |
| RM6697_w056_B08 | MT486169 | .............................N..........................H...S................G...N........ |
| RM6697_w056_B09 | MT486170 | ......D......................N..........................H...S................G...N........ |
| RM6697_w056_B11 | MT486171 | .............................N..........................H...S..........................N........ |
| RM6697_w056_B12 | MT486172 | .............................N..........................H...S...............E.....N........ |
| RM6697_w056_B17 | MT486174 | .............................N..........................H...S................G...N........ |
| RM6697_w056_B19 | MT486175 | .............................N..........................H...S..........................N........ |
| RM6697_w056_B20 | MT486176 | .............................N..........................H...S................G...N........ |
| RM6697_w056_B21 | MT486177 | .............................N..........................H...S...............E.....Y........ |
| RM6697_w056_B22 | MT486178 | .............................N..........................H...S..........................N........ |
| RM6697_w056_T01 | MT486179 | .............................N..........................H...S................G...N........ |
| RM6697_w056_T04 | MT486180 | .............................N..........................H...S...............EA............ |
| **RM6698 \| n = 3 \| 56 WPI** | | |
| RM6698_w056_B01 | MT486279 | ............................................................S............................. |
| RM6698_w056_B03 | MT486281 | ............................................................S............................A |
| RM6698_w056_T01 | MT486282 | .......D....................................................S....................R........ |
| **RM6699 \| n = 9 \| 56 WPI** | | |
| RM6699_w056_B01 | MT486349 | ......D.....................................................S....................Y........ |
| RM6699_w056_B04 | MT486351 | ......D.I.............V........................R............S............................. |
| RM6699_w056_B05 | MT486352 | ......D.I......................................R............S............................. |
| RM6699_w056_B06 | MT486353 | ......D.I......................................R............S............................. |
| RM6699_w056_B07 | MT486354 | ......D.I...................................................S............................. |
| RM6699_w056_T03 | MT486355 | ......D.....................................................S............................. |
| RM6699_w056_T04 | MT486356 | ......DSI...................................................S...............R....Y........ |
| RM6699_w056_T05 | MT486357 | ......D.I......................................R............S............................. |
| RM6699_w056_T06 | MT486358 | ......D.I...................................................S............................. |
| **RM6701 \| n = 18 \| 56 WPI** | | |
| RM6701_w056_A04 | MT486603 | ........................................................H...S............................. |
| RM6701_w056_A06 | MT486604 | ........................................................H...S............................. |
| RM6701_w056_A08 | MT486605 | ........................................................H...S............................. |
| RM6701_w056_A10 | MT486607 | ........................................................H...S............................. |
| RM6701_w056_A12 | MT486608 | ........................................................H...S............................. |
| RM6701_w056_A15 | MT486609 | ........................................................H...S............................. |
| RM6701_w056_A16 | MT486610 | ........................................................H...S............................. |
| RM6701_w056_A19 | MT486612 | ........................................................H...S............................. |
| RM6701_w056_A20 | MT486613 | ........................................................H...S............................. |
| RM6701_w056_A22 | MT486615 | ........................................................H...S............................. |
| RM6701_w056_B01 | MT486616 | ........................................................H...S............................. |
| RM6701_w056_B03 | MT486618 | ........................................................H...S............................. |
| RM6701_w056_B04 | MT486619 | ........................................................H...S............................. |
| RM6701_w056_B05 | MT486620 | ........................................................H...S............................. |
| RM6701_w056_B06 | MT486621 | ........................................................H...S............................. |
| RM6701_w056_T01 | MT486622 | ........................................................H...S............................. |
| RM6701_w056_T02 | MT486623 | ........................................................H...S............................. |
| RM6701_w056_T04 | MT486624 | ........................................................H...S............................. |
| **RM6702 \| n = 17 \| 24 WPI** | | |
| RM6702_w024_A01 | MT486654 | ......................................................K.....S............................. |
| RM6702_w024_A02 | MT486655 | ......................................................K.....S............................. |
| RM6702_w024_A03 | MT486656 | ......................................................K.....S............................. |
| RM6702_w024_A04 | MT486657 | ......................................................K.....S............................. |
| RM6702_w024_A05 | MT486658 | ......................................................K.....S............................. |
| RM6702_w024_A08 | MT486660 | ......................................................K.....S............................. |
| RM6702_w024_A10 | MT486661 | ......................................................K.....S............................. |
| RM6702_w024_A13 | MT486663 | ......................................................K.....S............................. |
| RM6702_w024_A14 | MT486664 | ......................................................K.....S............................. |
| RM6702_w024_A15 | MT486665 | ......................................................K.....S............................. |
| RM6702_w024_A16 | MT486666 | ......................................................K.....S............................. |
| RM6702_w024_A17 | MT486667 | ......................................................K.....S............................. |
| RM6702_w024_A18 | MT486668 | .......D..............................................K.....S............................. |
| RM6702_w024_A19 | MT486669 | ......................................................K.....S........................A.... |
| RM6702_w024_A20 | MT486670 | .......D..............................................K.....S............................. |
| RM6702_w024_T01 | MT486671 | ......................................................K.....S............................. |
| RM6702_w024_T02 | MT486672 | ................................I.....................K.....S........................A.... |
| **RM6703 \| n = 20 \| 48 WPI** | | |
| RM6703_w048_A03 | MT486741 | ............................................................S......A...................... |
| RM6703_w048_A04 | MT486742 | ............................................................S....S........................ |
| RM6703_w048_A05 | MT486743 | ............................................................S......A...................... |
| RM6703_w048_A06 | MT486744 | ............................................................S....S........................ |
| RM6703_w048_A08 | MT486746 | ............................................................S......A...................... |
| RM6703_w048_A11 | MT486747 | ......................................S.....................S......A...................... |
| RM6703_w048_A13 | MT486748 | .................K..........................................S....S...........K............ |
| RM6703_w048_A15 | MT486749 | ............................................................S....S...............Q........ |
| RM6703_w048_B01 | MT486750 | ............................................................S......A...................... |
| RM6703_w048_B04 | MT486752 | ............................................................S....S........................ |
| RM6703_w048_B06 | MT486753 | .......................S....................................S....S........................ |
| RM6703_w048_B07 | MT486754 | ............................................................S....S........................ |
| RM6703_w048_B08 | MT486755 | ............................................................S......A...................... |
| RM6703_w048_T01 | MT486756 | ........................................................H...S......A...................... |
| RM6703_w048_T02 | MT486757 | ............................................................S......A...................... |
| RM6703_w048_T03 | MT486758 | ............................................................S....S........................ |
| RM6703_w048_T06 | MT486759 | ............................................................S......A...................... |
| RM6703_w048_T07 | MT486760 | ............................................................S......A...................... |
| RM6703_w048_T08 | MT486761 | ............................................................S......A...................... |
| RM6703_w048_T09 | MT486762 | ............................................................S......A...................... |

. identity   A substitution   A insertion   − deletion   − gap at insertion column   X unresolved residue

* in numbering: insertion column without a supplied HXB2 position.

Domain and position annotations follow HXB2 numbering. SP, signal peptide; Fusion, fusion peptide.

MPER, membrane−proximal external region. Each sequence is displayed separately; no identical sequences are collapsed.

### CH505 Env amino−acid alignment

6 macaques | 105 sampled sequences | reference coverage: HXB2 1−709

Segment 5/8 | HXB2 365−454 | alignment columns 361−450

**Sequence identifier | GenBank accession**

Env region: C3 | V4 | C4

HXB2 numbering: 365 366 367 368 369 370 371 372 373 374 375 376 377 378 379 380 381 382 383 384 385 386 387 388 389 390 391 392 393 394 395 396 397 398 399 400 401 402 403 404 405 406 407 408 409 410 411 412 413 414 415 416 417 418 419 420 421 422 423 424 425 426 427 428 429 430 431 432 433 434 435 436 437 438 439 440 441 442 443 444 445 446 447 448 449 450 451 452 453 454

| Sequence identifier | GenBank accession | HXB2 365−454 |
|---|---|---|
| **SHIV.CH505.375H** | **QSQ65056** | SGGDLEITTHHFNCGGEFFYCNTSSLFNRTYMANSTDMANSTETNSTRTITIHCRIKQIINMWQEVGRAMYAPPIAGNITCISNITGLLL |
| **RM6697 \| n = 38 \| 56 WPI** | | |
| RM6697_w056_A02 | MT486140 | .......................................................................................... |
| RM6697_w056_A03 | MT486141 | .......................................................................................... |
| RM6697_w056_A04 | MT486142 | .......................................................................................... |
| RM6697_w056_A05 | MT486143 | ....................................................R..................................... |
| RM6697_w056_A06 | MT486144 | ................................D...................R..................................... |
| RM6697_w056_A07 | MT486145 | ....................................................R..................................... |
| RM6697_w056_A09 | MT486147 | ....................................................R..................................... |
| RM6697_w056_A11 | MT486148 | ................................D...................R..................................... |
| RM6697_w056_A12 | MT486149 | .......................................................................................... |
| RM6697_w056_A13 | MT486150 | ................................D...................R..................................... |
| RM6697_w056_A14 | MT486151 | ....................................................R..................................... |
| RM6697_w056_A15 | MT486152 | ...........................................................V.............................. |
| RM6697_w056_A16 | MT486153 | ................................D...................R..................................... |
| RM6697_w056_A17 | MT486154 | ................................D...................R..................................... |
| RM6697_w056_A19 | MT486155 | ....................................................R..................................... |
| RM6697_w056_A22 | MT486156 | .......................................................................................... |
| RM6697_w056_A23 | MT486157 | .......................................................................................... |
| RM6697_w056_A24 | MT486158 | ................................D...................R..................................... |
| RM6697_w056_A25 | MT486159 | .......................................................................................... |
| RM6697_w056_A27 | MT486160 | .......................................................................................... |
| RM6697_w056_A30 | MT486161 | ................................D...................R..................................... |
| RM6697_w056_A31 | MT486162 | ................................D...................R..................................... |
| RM6697_w056_B01 | MT486164 | ....................................................R..................................... |
| RM6697_w056_B03 | MT486165 | ................................D...................R..................................... |
| RM6697_w056_B04 | MT486166 | ................................D...................R..................................... |
| RM6697_w056_B06 | MT486167 | ................................D...................R..................................... |
| RM6697_w056_B07 | MT486168 | .......................................................................................... |
| RM6697_w056_B08 | MT486169 | .......................................................................................... |
| RM6697_w056_B09 | MT486170 | ....................................................R..................................... |
| RM6697_w056_B11 | MT486171 | ....................................................R..................................... |
| RM6697_w056_B12 | MT486172 | .......................................................................................... |
| RM6697_w056_B17 | MT486174 | .......................................................................................... |
| RM6697_w056_B19 | MT486175 | ....................................................R..................................... |
| RM6697_w056_B20 | MT486176 | ....................................................R.................H................... |
| RM6697_w056_B21 | MT486177 | .....................D........................I.....R..................................... |
| RM6697_w056_B22 | MT486178 | ................................D...................R..................................... |
| RM6697_w056_T01 | MT486179 | ................................D...................R..................................... |
| RM6697_w056_T04 | MT486180 | ....................................................R..................................... |
| **RM6698 \| n = 3 \| 56 WPI** | | |
| RM6698_w056_B01 | MT486279 | .......................................................................................... |
| RM6698_w056_B03 | MT486281 | .......................................................................................... |
| RM6698_w056_T01 | MT486282 | .......................................................................................... |
| **RM6699 \| n = 9 \| 56 WPI** | | |
| RM6699_w056_B01 | MT486349 | ................................................I...R...................................I. |
| RM6699_w056_B04 | MT486351 | ....................................................R...................................I. |
| RM6699_w056_B05 | MT486352 | ....................................................R...................................I. |
| RM6699_w056_B06 | MT486353 | ....................................................R...................................I. |
| RM6699_w056_B07 | MT486354 | ....................................................R..................................... |
| RM6699_w056_T03 | MT486355 | ....................................................R..................................... |
| RM6699_w056_T04 | MT486356 | ....................................................R..................................... |
| RM6699_w056_T05 | MT486357 | ....................................................R...................................I. |
| RM6699_w056_T06 | MT486358 | ....................................................R..................................... |
| **RM6701 \| n = 18 \| 56 WPI** | | |
| RM6701_w056_A04 | MT486603 | ........................G................................................................. |
| RM6701_w056_A06 | MT486604 | ........................G.......V......................................................... |
| RM6701_w056_A08 | MT486605 | ........................G................................................................. |
| RM6701_w056_A10 | MT486607 | ........................G................................................................. |
| RM6701_w056_A12 | MT486608 | ........................G................................................................. |
| RM6701_w056_A15 | MT486609 | ........................G................................................................. |
| RM6701_w056_A16 | MT486610 | ........................G................................................................. |
| RM6701_w056_A19 | MT486612 | ........................G................................................................. |
| RM6701_w056_A20 | MT486613 | ........................G................................................................. |
| RM6701_w056_A22 | MT486615 | ........................G................................................................. |
| RM6701_w056_B01 | MT486616 | ........................G................................................................. |
| RM6701_w056_B03 | MT486618 | ........................G................................................................. |
| RM6701_w056_B04 | MT486619 | ........................G................................................................. |
| RM6701_w056_B05 | MT486620 | ........................G.......V.........K............................................... |
| RM6701_w056_B06 | MT486621 | ........................G................................................................. |
| RM6701_w056_T01 | MT486622 | ........................G................................................................. |
| RM6701_w056_T02 | MT486623 | ........................G................................................................. |
| RM6701_w056_T04 | MT486624 | ........................G.......V......................................................... |
| **RM6702 \| n = 17 \| 24 WPI** | | |
| RM6702_w024_A01 | MT486654 | .......................................................................................... |
| RM6702_w024_A02 | MT486655 | .......................................................................................... |
| RM6702_w024_A03 | MT486656 | .......................................................................................... |
| RM6702_w024_A04 | MT486657 | .......................................................................................... |
| RM6702_w024_A05 | MT486658 | .......................................................................................... |
| RM6702_w024_A08 | MT486660 | .......................................................................................... |
| RM6702_w024_A10 | MT486661 | .......................................................................................... |
| RM6702_w024_A13 | MT486663 | .......................................................................................... |
| RM6702_w024_A14 | MT486664 | .......................................................................................... |
| RM6702_w024_A15 | MT486665 | .......................................................................................... |
| RM6702_w024_A16 | MT486666 | .......................................................................................... |
| RM6702_w024_A17 | MT486667 | .......................................................................................... |
| RM6702_w024_A18 | MT486668 | .......................................................................................... |
| RM6702_w024_A19 | MT486669 | ......V.........................................N......................................... |
| RM6702_w024_A20 | MT486670 | .......................................................................................... |
| RM6702_w024_T01 | MT486671 | .......................................................................................... |
| RM6702_w024_T02 | MT486672 | ................................................N......................................... |
| **RM6703 \| n = 20 \| 48 WPI** | | |
| RM6703_w048_A03 | MT486741 | ................................................I...R..................................... |
| RM6703_w048_A04 | MT486742 | ................................................I...R..................................... |
| RM6703_w048_A05 | MT486743 | ................................................I...R..................................... |
| RM6703_w048_A06 | MT486744 | ................................................I...R..................................... |
| RM6703_w048_A08 | MT486746 | ................................................I...R..................................... |
| RM6703_w048_A11 | MT486747 | ................................................I...R..................................... |
| RM6703_w048_A13 | MT486748 | ................................................I...R.............E....................... |
| RM6703_w048_A15 | MT486749 | ................................................I...R..................................... |
| RM6703_w048_B01 | MT486750 | ................................................I...R..................................... |
| RM6703_w048_B04 | MT486752 | ................................................I...R..................................... |
| RM6703_w048_B06 | MT486753 | ................................................I...R..................................... |
| RM6703_w048_B07 | MT486754 | ................................................I...R..................................... |
| RM6703_w048_B08 | MT486755 | ................................................I...R..................................... |
| RM6703_w048_T01 | MT486756 | ................................................I...R..................................... |
| RM6703_w048_T02 | MT486757 | ...........................................A.............................................. |
| RM6703_w048_T03 | MT486758 | ................................................I...R..................................... |
| RM6703_w048_T06 | MT486759 | ................................................I...R..................................... |
| RM6703_w048_T07 | MT486760 | ................................................I...R..................................... |
| RM6703_w048_T08 | MT486761 | ....................................................R..................................... |
| RM6703_w048_T09 | MT486762 | ................................................I...R..................................... |

. identity     A substitution     A insertion     − deletion     − gap at insertion column     X unresolved residue

* in numbering: insertion column without a supplied HXB2 position.

Domain and position annotations follow HXB2 numbering. SP, signal peptide; Fusion, fusion peptide.

MPER, membrane−proximal external region. Each sequence is displayed separately; no identical sequences are collapsed.

### CH505 Env amino−acid alignment

6 macaques | 105 sampled sequences | reference coverage: HXB2 1−709

Segment 6/8 | HXB2 455−546 | alignment columns 451−540

**Sequence identifier | GenBank accession**

Env region: C4 | V5 | C5 | Fusion

HXB2 numbering: 455 456 457 458 459 460 461 462 463 464 465 468 469 470 471 472 473 474 475 476 477 478 479 480 481 482 483 484 485 486 487 488 489 490 491 492 493 494 495 496 497 498 499 500 501 502 503 504 505 506 507 508 509 510 511 512 513 514 515 516 517 518 519 520 521 522 523 524 525 526 527 528 529 530 531 532 533 534 535 536 537 538 539 540 541 542 543 544 545 546

| Sequence identifier | GenBank accession | Alignment |
|---|---|---|
| **SHIV.CH505.375H** | **QSQ65056** | TRDGGKNNTETFRPGGGNMKDNWRSELYKYKVVEVKPLGVAPTNARRRVVEREKRAVGMGAVFLGFLGAAGSTMGAASITLTVQARQLLS |
| **RM6697 \| n = 38 \| 56 WPI** | | |
| RM6697_w056_A02 | MT486140 | .....E............L....................................................................... |
| RM6697_w056_A03 | MT486141 | .....E.................................................................................... |
| RM6697_w056_A04 | MT486142 | .....E.................................................................................... |
| RM6697_w056_A05 | MT486143 | .....E..M................................................................................. |
| RM6697_w056_A06 | MT486144 | .....E.................................................................................... |
| RM6697_w056_A07 | MT486145 | .....E.................................................................................... |
| RM6697_w056_A09 | MT486147 | .....E.................................................................................... |
| RM6697_w056_A11 | MT486148 | .....E.................................................................................... |
| RM6697_w056_A12 | MT486149 | .....E.D.................................................................................. |
| RM6697_w056_A13 | MT486150 | .....E.................................................................................... |
| RM6697_w056_A14 | MT486151 | .....E.................................................................................... |
| RM6697_w056_A15 | MT486152 | .....E.................................................................................... |
| RM6697_w056_A16 | MT486153 | .....E.................................................................................... |
| RM6697_w056_A17 | MT486154 | .....E.................................................................................... |
| RM6697_w056_A19 | MT486155 | .....E.................................................................................... |
| RM6697_w056_A22 | MT486156 | .....E.................................................................................... |
| RM6697_w056_A23 | MT486157 | .....E...................................................E................................ |
| RM6697_w056_A24 | MT486158 | .....E.................................................................................... |
| RM6697_w056_A25 | MT486159 | .....E.................................................................................... |
| RM6697_w056_A27 | MT486160 | .....E............L....................................................................... |
| RM6697_w056_A30 | MT486161 | .....E.................................................................................... |
| RM6697_w056_A31 | MT486162 | .....E.................................................................................... |
| RM6697_w056_B01 | MT486164 | .....E.................................................................................... |
| RM6697_w056_B03 | MT486165 | .....E.................................................................................... |
| RM6697_w056_B04 | MT486166 | .....E.................................................................................... |
| RM6697_w056_B06 | MT486167 | .....E.................................................................................... |
| RM6697_w056_B07 | MT486168 | .....E.................................................................................... |
| RM6697_w056_B08 | MT486169 | .....E.................................................................................... |
| RM6697_w056_B09 | MT486170 | .....E.................................................................................... |
| RM6697_w056_B11 | MT486171 | .....E.................................I............G..................................... |
| RM6697_w056_B12 | MT486172 | .....E.................................................................................... |
| RM6697_w056_B17 | MT486174 | .....E............L....................................................................... |
| RM6697_w056_B19 | MT486175 | .....E.D.................................................................................. |
| RM6697_w056_B20 | MT486176 | .....E.................................................................................... |
| RM6697_w056_B21 | MT486177 | .....E.DM................................................................................. |
| RM6697_w056_B22 | MT486178 | .....E.................................................................................... |
| RM6697_w056_T01 | MT486179 | .....E.................................................................................... |
| RM6697_w056_T04 | MT486180 | .....E.................................................................................... |
| **RM6698 \| n = 3 \| 56 WPI** | | |
| RM6698_w056_B01 | MT486279 | .....R.................................................................................... |
| RM6698_w056_B03 | MT486281 | .....R.................................................................................... |
| RM6698_w056_T01 | MT486282 | .................D......N................................................................. |
| **RM6699 \| n = 9 \| 56 WPI** | | |
| RM6699_w056_B01 | MT486349 | .....E....I............................................................................... |
| RM6699_w056_B04 | MT486351 | .....E....I............................................................................... |
| RM6699_w056_B05 | MT486352 | .....E....I............................................................................... |
| RM6699_w056_B06 | MT486353 | .....E....I............................................................................... |
| RM6699_w056_B07 | MT486354 | ....E..D.................................................................................. |
| RM6699_w056_T03 | MT486355 | ....E..D.................................................................................. |
| RM6699_w056_T04 | MT486356 | ....E..............................................................................R...... |
| RM6699_w056_T05 | MT486357 | .....E....I............................................................................... |
| RM6699_w056_T06 | MT486358 | ....E..D.................................................................................. |
| **RM6701 \| n = 18 \| 56 WPI** | | |
| RM6701_w056_A04 | MT486603 | .....E..A................................................................................. |
| RM6701_w056_A06 | MT486604 | .....E..A................................................................................. |
| RM6701_w056_A08 | MT486605 | .....E..A................................................................................. |
| RM6701_w056_A10 | MT486607 | .....E..A................................................................................. |
| RM6701_w056_A12 | MT486608 | .....E..A................................................................................. |
| RM6701_w056_A15 | MT486609 | .....E..A................................................................................. |
| RM6701_w056_A16 | MT486610 | .....E..A................................................................................. |
| RM6701_w056_A19 | MT486612 | .....E..A................................................................................. |
| RM6701_w056_A20 | MT486613 | .....E..A................................................................................. |
| RM6701_w056_A22 | MT486615 | .....E..A................................................................................. |
| RM6701_w056_B01 | MT486616 | .....E..A................................................................................. |
| RM6701_w056_B03 | MT486618 | .....E..A................................................................................. |
| RM6701_w056_B04 | MT486619 | .......DA................................................................................. |
| RM6701_w056_B05 | MT486620 | .....E..A...................................T............................................. |
| RM6701_w056_B06 | MT486621 | .....E..A................................................................................. |
| RM6701_w056_T01 | MT486622 | .....E..A................................................................................. |
| RM6701_w056_T02 | MT486623 | .....E..A................................................................................. |
| RM6701_w056_T04 | MT486624 | .....E..A................................................................................. |
| **RM6702 \| n = 17 \| 24 WPI** | | |
| RM6702_w024_A01 | MT486654 | .....E.D.................................................................................. |
| RM6702_w024_A02 | MT486655 | ....E.D................................................................................... |
| RM6702_w024_A03 | MT486656 | .....E.................................................................................... |
| RM6702_w024_A04 | MT486657 | .....N.D.................................................................................. |
| RM6702_w024_A05 | MT486658 | .....E.................................................................................... |
| RM6702_w024_A08 | MT486660 | .....E.................................................................................... |
| RM6702_w024_A10 | MT486661 | .....E.................................................................................... |
| RM6702_w024_A13 | MT486663 | .....E.................................................................................... |
| RM6702_w024_A14 | MT486664 | .....E.................................................................................... |
| RM6702_w024_A15 | MT486665 | .....E.................................................................................... |
| RM6702_w024_A16 | MT486666 | .....E.................................................................................... |
| RM6702_w024_A17 | MT486667 | ....E.D.........E........K................................................................ |
| RM6702_w024_A18 | MT486668 | .......D.................................................................................. |
| RM6702_w024_A19 | MT486669 | .....E.................................................................................... |
| RM6702_w024_A20 | MT486670 | ......TD.................................................................................. |
| RM6702_w024_T01 | MT486671 | .....E.S...............................I.................................................. |
| RM6702_w024_T02 | MT486672 | .....E.................................................................................... |
| **RM6703 \| n = 20 \| 48 WPI** | | |
| RM6703_w048_A03 | MT486741 | .......................................................................................... |
| RM6703_w048_A04 | MT486742 | ...................................................K...................................... |
| RM6703_w048_A05 | MT486743 | .......................................................................................... |
| RM6703_w048_A06 | MT486744 | ...................................................K...................................... |
| RM6703_w048_A08 | MT486746 | .......................................I.................................................. |
| RM6703_w048_A11 | MT486747 | .......................................................................................... |
| RM6703_w048_A13 | MT486748 | ..N......................................................E................................ |
| RM6703_w048_A15 | MT486749 | .......................................................................................... |
| RM6703_w048_B01 | MT486750 | .......................................................................................... |
| RM6703_w048_B04 | MT486752 | .......................................................................................... |
| RM6703_w048_B06 | MT486753 | .......................................................................................... |
| RM6703_w048_B07 | MT486754 | ...................................................K..................................R... |
| RM6703_w048_B08 | MT486755 | ......................................................K................................... |
| RM6703_w048_T01 | MT486756 | .......................................................................................... |
| RM6703_w048_T02 | MT486757 | .......................................................................................... |
| RM6703_w048_T03 | MT486758 | .......................................................................................... |
| RM6703_w048_T06 | MT486759 | .......................................................................................... |
| RM6703_w048_T07 | MT486760 | .......................................................................................... |
| RM6703_w048_T08 | MT486761 | .......................................................................................... |
| RM6703_w048_T09 | MT486762 | .......................................................................................... |

. identity    A substitution    A insertion    − deletion    − gap at insertion column    X unresolved residue

* in numbering: insertion column without a supplied HXB2 position.

Domain and position annotations follow HXB2 numbering. SP, signal peptide; Fusion, fusion peptide.

MPER, membrane−proximal external region. Each sequence is displayed separately; no identical sequences are collapsed.

### CH505 Env amino−acid alignment

6 macaques | 105 sampled sequences | reference coverage: HXB2 1−709

Segment 7/8 | HXB2 547−636 | alignment columns 541−630

**Sequence identifier | GenBank accession**

Env region

| HXB2 numbering | 547–636 |
|---|---|
| **SHIV.CH505.375H \| QSQ65056** | GIVQQQSNLLKAIEAQQHMLKLTVWGIKQLQARVLALERYLKDQQLLGMWGCSGKLICTTNVYWNSSWSNKTYGDIWDNMTWMQWEREIS |
| **RM6697 \| n = 38 \| 56 WPI** | |
| RM6697_w056_A02 \| MT486140 | .......................................................................................... |
| RM6697_w056_A03 \| MT486141 | .......................................................................................... |
| RM6697_w056_A04 \| MT486142 | .......................................................................................... |
| RM6697_w056_A05 \| MT486143 | ....................R..................................................................... |
| RM6697_w056_A06 \| MT486144 | ....................R..................................................................... |
| RM6697_w056_A07 \| MT486145 | .......................................................................................... |
| RM6697_w056_A09 \| MT486147 | ....................R..................................................................... |
| RM6697_w056_A11 \| MT486148 | .........................S................................................................ |
| RM6697_w056_A12 \| MT486149 | ....................R..................................................................... |
| RM6697_w056_A13 \| MT486150 | ....................R..................................................................... |
| RM6697_w056_A14 \| MT486151 | .......................................................................................... |
| RM6697_w056_A15 \| MT486152 | ....................R..................................................................... |
| RM6697_w056_A16 \| MT486153 | ....................R..................................................................... |
| RM6697_w056_A17 \| MT486154 | .......................................................................................... |
| RM6697_w056_A19 \| MT486155 | ....................R..................................................................... |
| RM6697_w056_A22 \| MT486156 | ....................R..................................................................... |
| RM6697_w056_A23 \| MT486157 | ....................R..................................................................... |
| RM6697_w056_A24 \| MT486158 | ....................R..................................................................... |
| RM6697_w056_A25 \| MT486159 | ....................R..................................................................... |
| RM6697_w056_A27 \| MT486160 | .......................................................................................... |
| RM6697_w056_A30 \| MT486161 | ....................R..................................................................... |
| RM6697_w056_A31 \| MT486162 | ....................R..................................................................... |
| RM6697_w056_B01 \| MT486164 | ....................R..................................................................... |
| RM6697_w056_B03 \| MT486165 | ....................R..................................................................... |
| RM6697_w056_B04 \| MT486166 | ....................R...........................I......................................... |
| RM6697_w056_B06 \| MT486167 | ....................R..................................................................... |
| RM6697_w056_B07 \| MT486168 | ....................R..................................................................... |
| RM6697_w056_B08 \| MT486169 | ....................R..................................................................... |
| RM6697_w056_B09 \| MT486170 | ....................R..................................................................... |
| RM6697_w056_B11 \| MT486171 | ....................R..................................................................... |
| RM6697_w056_B12 \| MT486172 | .....................................................................S................K... |
| RM6697_w056_B17 \| MT486174 | .......................................................................................... |
| RM6697_w056_B19 \| MT486175 | ....................R..................................................................... |
| RM6697_w056_B20 \| MT486176 | ....................R..................................................................... |
| RM6697_w056_B21 \| MT486177 | ....................R..................................................................... |
| RM6697_w056_B22 \| MT486178 | ....................R..................................................................... |
| RM6697_w056_T01 \| MT486179 | ....................R..................................................................... |
| RM6697_w056_T04 \| MT486180 | ....................R..................................................................... |
| **RM6698 \| n = 3 \| 56 WPI** | |
| RM6698_w056_B01 \| MT486279 | .......................................................................................... |
| RM6698_w056_B03 \| MT486281 | .......................................................................................... |
| RM6698_w056_T01 \| MT486282 | .......................................................................................... |
| **RM6699 \| n = 9 \| 56 WPI** | |
| RM6699_w056_B01 \| MT486349 | .......................................................................................... |
| RM6699_w056_B04 \| MT486351 | .......................................................................................... |
| RM6699_w056_B05 \| MT486352 | .......................................................................................... |
| RM6699_w056_B06 \| MT486353 | .......................................................................................... |
| RM6699_w056_B07 \| MT486354 | ....................R..................................................................... |
| RM6699_w056_T03 \| MT486355 | ....................R..................................................................... |
| RM6699_w056_T04 \| MT486356 | .......................................................................................... |
| RM6699_w056_T05 \| MT486357 | .......................................................................................... |
| RM6699_w056_T06 \| MT486358 | ....................R..................................................................... |
| **RM6701 \| n = 18 \| 56 WPI** | |
| RM6701_w056_A04 \| MT486603 | .......................................................................................... |
| RM6701_w056_A06 \| MT486604 | .......................................................................................... |
| RM6701_w056_A08 \| MT486605 | .......................................................................................... |
| RM6701_w056_A10 \| MT486607 | .......................................................................................... |
| RM6701_w056_A12 \| MT486608 | .................................................................F........................ |
| RM6701_w056_A15 \| MT486609 | .......................................................................................... |
| RM6701_w056_A16 \| MT486610 | ....................R................................................S.................... |
| RM6701_w056_A19 \| MT486612 | .......................................................................................... |
| RM6701_w056_A20 \| MT486613 | .............................................................I............................ |
| RM6701_w056_A22 \| MT486615 | .......................................................................................... |
| RM6701_w056_B01 \| MT486616 | .......................................................................................... |
| RM6701_w056_B03 \| MT486618 | .......................................................................................... |
| RM6701_w056_B04 \| MT486619 | .......................................................................................... |
| RM6701_w056_B05 \| MT486620 | .......................................................................................... |
| RM6701_w056_B06 \| MT486621 | .......................................................................................... |
| RM6701_w056_T01 \| MT486622 | .......................................................................................... |
| RM6701_w056_T02 \| MT486623 | .......................................................................................... |
| RM6701_w056_T04 \| MT486624 | .......................................................................................... |
| **RM6702 \| n = 17 \| 24 WPI** | |
| RM6702_w024_A01 \| MT486654 | .......................................................................................... |
| RM6702_w024_A02 \| MT486655 | .......................................................................................... |
| RM6702_w024_A03 \| MT486656 | .......................................................................................... |
| RM6702_w024_A04 \| MT486657 | .......................................................................................... |
| RM6702_w024_A05 \| MT486658 | .......................................................................................... |
| RM6702_w024_A08 \| MT486660 | .......................................................................................... |
| RM6702_w024_A10 \| MT486661 | ................................................................D......................... |
| RM6702_w024_A13 \| MT486663 | .......................................................................................... |
| RM6702_w024_A14 \| MT486664 | .......................................................................................... |
| RM6702_w024_A15 \| MT486665 | .......................................................................................... |
| RM6702_w024_A16 \| MT486666 | .......................................................................................... |
| RM6702_w024_A17 \| MT486667 | .......................................................................................... |
| RM6702_w024_A18 \| MT486668 | .......................................................................................... |
| RM6702_w024_A19 \| MT486669 | ................................................................D......................... |
| RM6702_w024_A20 \| MT486670 | ..............................................................................S........... |
| RM6702_w024_T01 \| MT486671 | .......................................................................................... |
| RM6702_w024_T02 \| MT486672 | .......................................................................................... |
| **RM6703 \| n = 20 \| 48 WPI** | |
| RM6703_w048_A03 \| MT486741 | .......................................................................................... |
| RM6703_w048_A04 \| MT486742 | .......................................................................................... |
| RM6703_w048_A05 \| MT486743 | .......................................................................................... |
| RM6703_w048_A06 \| MT486744 | .......................................................................................... |
| RM6703_w048_A08 \| MT486746 | .......................................................................................... |
| RM6703_w048_A11 \| MT486747 | .......................................................................................... |
| RM6703_w048_A13 \| MT486748 | .......................................................................................... |
| RM6703_w048_A15 \| MT486749 | .......................................................................................... |
| RM6703_w048_B01 \| MT486750 | .......................................................................................... |
| RM6703_w048_B04 \| MT486752 | .......................................................................................... |
| RM6703_w048_B06 \| MT486753 | .......................................................................................... |
| RM6703_w048_B07 \| MT486754 | .......................................................................................... |
| RM6703_w048_B08 \| MT486755 | .......................................................................................... |
| RM6703_w048_T01 \| MT486756 | .......................................................................................... |
| RM6703_w048_T02 \| MT486757 | ......................................................................................K... |
| RM6703_w048_T03 \| MT486758 | .......................................................................................... |
| RM6703_w048_T06 \| MT486759 | .......................................................................................... |
| RM6703_w048_T07 \| MT486760 | .......................................................................................... |
| RM6703_w048_T08 \| MT486761 | .......................................................................................... |
| RM6703_w048_T09 \| MT486762 | .......................................................................................... |

. identity    A substitution    A insertion    − deletion    − gap at insertion column    X unresolved residue

* in numbering: insertion column without a supplied HXB2 position.

Domain and position annotations follow HXB2 numbering. SP, signal peptide; Fusion, fusion peptide.

MPER, membrane−proximal external region. Each sequence is displayed separately; no identical sequences are collapsed.

### CH505 Env amino−acid alignment

6 macaques | 105 sampled sequences | reference coverage: HXB2 1−709

Segment 8/8 | HXB2 637−709 | alignment columns 631−703

**Sequence identifier | GenBank accession**

Env region: MPER (HXB2 662−683)

HXB2 numbering: 637 638 639 640 641 642 643 644 645 646 647 648 649 650 651 652 653 654 655 656 657 658 659 660 661 662 663 664 665 666 667 668 669 670 671 672 673 674 675 676 677 678 679 680 681 682 683 684 685 686 687 688 689 690 691 692 693 694 695 696 697 698 699 700 701 702 703 704 705 706 707 708 709

| Sequence identifier | GenBank accession | HXB2 637−709 |
|---|---|---|
| **SHIV.CH505.375H** | **QSQ65056** | NYTEIIYELLEESQNQQEKNEQDLLALDRWNSLWNWFNITNWLWYIKIFIMIVGGLIGLRIIFAVLSLVNRVR |

##### RM6697 | n = 38 | 56 WPI

| Sequence identifier | GenBank accession | HXB2 637−709 |
|---|---|---|
| RM6697_w056_A02 | MT486140 | .......................................A................................. |
| RM6697_w056_A03 | MT486141 | .............................................M.............K............. |
| RM6697_w056_A04 | MT486142 | ......................................................................... |
| RM6697_w056_A05 | MT486143 | .............................................M........................... |
| RM6697_w056_A06 | MT486144 | .............................................M........................... |
| RM6697_w056_A07 | MT486145 | ....T........................................M........................... |
| RM6697_w056_A09 | MT486147 | ......................................................................... |
| RM6697_w056_A11 | MT486148 | .............................................M........................... |
| RM6697_w056_A12 | MT486149 | .............................................M........................... |
| RM6697_w056_A13 | MT486150 | .......................................A................................. |
| RM6697_w056_A14 | MT486151 | ......................................................................... |
| RM6697_w056_A15 | MT486152 | .............................................M........................... |
| RM6697_w056_A16 | MT486153 | .......................................A................................. |
| RM6697_w056_A17 | MT486154 | .............................................M........................... |
| RM6697_w056_A19 | MT486155 | .............................................M........................... |
| RM6697_w056_A22 | MT486156 | .......................................A................................. |
| RM6697_w056_A23 | MT486157 | .............................................M........................... |
| RM6697_w056_A24 | MT486158 | .............................................M........................... |
| RM6697_w056_A25 | MT486159 | .............................................M........................... |
| RM6697_w056_A27 | MT486160 | .......................................A................................. |
| RM6697_w056_A30 | MT486161 | .............................................M........................... |
| RM6697_w056_A31 | MT486162 | .............................................M........................... |
| RM6697_w056_B01 | MT486164 | ......................................................................... |
| RM6697_w056_B03 | MT486165 | .............................................M........................... |
| RM6697_w056_B04 | MT486166 | .............................................M........................... |
| RM6697_w056_B06 | MT486167 | .......................................A................................. |
| RM6697_w056_B07 | MT486168 | .............................................M........................... |
| RM6697_w056_B08 | MT486169 | .............................................M........................... |
| RM6697_w056_B09 | MT486170 | .............................................M........................... |
| RM6697_w056_B11 | MT486171 | .............................................M........................... |
| RM6697_w056_B12 | MT486172 | .......................................A................................. |
| RM6697_w056_B17 | MT486174 | .......................................A................................. |
| RM6697_w056_B19 | MT486175 | ......................................................................... |
| RM6697_w056_B20 | MT486176 | .......................................A................................. |
| RM6697_w056_B21 | MT486177 | ......................................................................... |
| RM6697_w056_B22 | MT486178 | .......................................A................................. |
| RM6697_w056_T01 | MT486179 | .......................................A................................. |
| RM6697_w056_T04 | MT486180 | .......................................A................................. |

##### RM6698 | n = 3 | 56 WPI

| Sequence identifier | GenBank accession | HXB2 637−709 |
|---|---|---|
| RM6698_w056_B01 | MT486279 | ......................................................................... |
| RM6698_w056_B03 | MT486281 | ..........K.............................................................. |
| RM6698_w056_T01 | MT486282 | .......................................A................................. |

##### RM6699 | n = 9 | 56 WPI

| Sequence identifier | GenBank accession | HXB2 637−709 |
|---|---|---|
| RM6699_w056_B01 | MT486349 | ......................................................................... |
| RM6699_w056_B04 | MT486351 | ......................................................................... |
| RM6699_w056_B05 | MT486352 | ......................................................................... |
| RM6699_w056_B06 | MT486353 | ......................................................................... |
| RM6699_w056_B07 | MT486354 | .......D................................................................. |
| RM6699_w056_T03 | MT486355 | ......................................................................... |
| RM6699_w056_T04 | MT486356 | ......................................................................... |
| RM6699_w056_T05 | MT486357 | ......................................................................... |
| RM6699_w056_T06 | MT486358 | ......................................................................... |

##### RM6701 | n = 18 | 56 WPI

| Sequence identifier | GenBank accession | HXB2 637−709 |
|---|---|---|
| RM6701_w056_A04 | MT486603 | ......................................................................... |
| RM6701_w056_A06 | MT486604 | ......................................................................... |
| RM6701_w056_A08 | MT486605 | ......................................................................... |
| RM6701_w056_A10 | MT486607 | ......................................................................... |
| RM6701_w056_A12 | MT486608 | ......................................................................... |
| RM6701_w056_A15 | MT486609 | ......................................................................... |
| RM6701_w056_A16 | MT486610 | ........................................S................................ |
| RM6701_w056_A19 | MT486612 | ......................................................................... |
| RM6701_w056_A20 | MT486613 | ......................................................................... |
| RM6701_w056_A22 | MT486615 | ......................................................................... |
| RM6701_w056_B01 | MT486616 | ......................................................................... |
| RM6701_w056_B03 | MT486618 | ......................................................................... |
| RM6701_w056_B04 | MT486619 | ......................................................................... |
| RM6701_w056_B05 | MT486620 | ......................................................................... |
| RM6701_w056_B06 | MT486621 | ......................................................................... |
| RM6701_w056_T01 | MT486622 | ......................................................................... |
| RM6701_w056_T02 | MT486623 | ......................................................................... |
| RM6701_w056_T04 | MT486624 | .....X................................................................... |

##### RM6702 | n = 17 | 24 WPI

| Sequence identifier | GenBank accession | HXB2 637−709 |
|---|---|---|
| RM6702_w024_A01 | MT486654 | ......................................................................S.. |
| RM6702_w024_A02 | MT486655 | ......................................................................S.. |
| RM6702_w024_A03 | MT486656 | ......................................................................... |
| RM6702_w024_A04 | MT486657 | ......................................................................S.. |
| RM6702_w024_A05 | MT486658 | ......................................................................... |
| RM6702_w024_A08 | MT486660 | ......................................................................... |
| RM6702_w024_A10 | MT486661 | ......................................................................... |
| RM6702_w024_A13 | MT486663 | ......................................................................... |
| RM6702_w024_A14 | MT486664 | ......................................................................... |
| RM6702_w024_A15 | MT486665 | ......................................................................... |
| RM6702_w024_A16 | MT486666 | ......................................................................... |
| RM6702_w024_A17 | MT486667 | ......................................................................... |
| RM6702_w024_A18 | MT486668 | ......................................................................... |
| RM6702_w024_A19 | MT486669 | ......................................................................... |
| RM6702_w024_A20 | MT486670 | ..................................................I...................... |
| RM6702_w024_T01 | MT486671 | ......................................................................... |
| RM6702_w024_T02 | MT486672 | ......................................................................... |

##### RM6703 | n = 20 | 48 WPI

| Sequence identifier | GenBank accession | HXB2 637−709 |
|---|---|---|
| RM6703_w048_A03 | MT486741 | ......................................................................... |
| RM6703_w048_A04 | MT486742 | ......................................................................... |
| RM6703_w048_A05 | MT486743 | ......................................................................... |
| RM6703_w048_A06 | MT486744 | ......................................................................... |
| RM6703_w048_A08 | MT486746 | ......................................................................... |
| RM6703_w048_A11 | MT486747 | ......................................................................... |
| RM6703_w048_A13 | MT486748 | ...........................................................K............. |
| RM6703_w048_A15 | MT486749 | ......................................................................... |
| RM6703_w048_B01 | MT486750 | ......................................................................... |
| RM6703_w048_B04 | MT486752 | ......................................................................... |
| RM6703_w048_B06 | MT486753 | ......................................................................... |
| RM6703_w048_B07 | MT486754 | ........................................H................................ |
| RM6703_w048_B08 | MT486755 | ......................................................................... |
| RM6703_w048_T01 | MT486756 | ...........................................R............................. |
| RM6703_w048_T02 | MT486757 | ......................................................................... |
| RM6703_w048_T03 | MT486758 | ......................................................................... |
| RM6703_w048_T06 | MT486759 | ......................................................................... |
| RM6703_w048_T07 | MT486760 | ......................................................................... |
| RM6703_w048_T08 | MT486761 | ......................................................................... |
| RM6703_w048_T09 | MT486762 | ......................................................................... |

. identity    A substitution    A insertion    − deletion    − gap at insertion column    X unresolved residue

* in numbering: insertion column without a supplied HXB2 position.

Domain and position annotations follow HXB2 numbering. SP, signal peptide; Fusion, fusion peptide.

MPER, membrane−proximal external region. Each sequence is displayed separately; no identical sequences are collapsed.
