## Supplementary material for "Functional Convergence of Genetically Diverse B-Cell Receptors in Simian-HIV Infected Rhesus Macaques": Document S3: Document_S3_Phylogenetic_Trees.html

Phylogenetic Trees of Identified Lineages


Phylogenetic Trees of Identified Lineages  
  
- RM6708
- RMT283
- RM6706
- RM6697
- RM6698
- RM6699
  
  

- RM6708\_C009
- RM6708\_C012
- RM6708\_C013
- RM6708\_C016
- RM6708\_C023
- RM6708\_C026
- RM6708\_C030
- RM6708\_C033
- RM6708\_C035
- RM6708\_C039

- RMT283\_C005
- RMT283\_C011
- RMT283\_C014
- RMT283\_C025
- RMT283\_C037
- RMT283\_C047
- RMT283\_C057
- RMT283\_C058
- RMT283\_C062
- RMT283\_C069
- RMT283\_C070
- RMT283\_C078
- RMT283\_C080
- RMT283\_C081
- RMT283\_C086
- RMT283\_C087
- RMT283\_C090
- RMT283\_C092
- RMT283\_C095
- RMT283\_C097
- RMT283\_C105
- RMT283\_C106
- RMT283\_C107
- RMT283\_C108
- RMT283\_C110
- RMT283\_C113
- RMT283\_C114
- RMT283\_C118
- RMT283\_C121
- RMT283\_C130
- RMT283\_C135
- RMT283\_C136
- RMT283\_C139
- RMT283\_C146
- RMT283\_C155
- RMT283\_C166
- RMT283\_C172
- RMT283\_C175
- RMT283\_C176
- RMT283\_C178
- RMT283\_C183
- RMT283\_C190
- RMT283\_C191

- RM6706\_C002
- RM6706\_C004
- RM6706\_C015
- RM6706\_C028
- RM6706\_C036
- RM6706\_C043
- RM6706\_C045
- RM6706\_C048
- RM6706\_C049
- RM6706\_C055
- RM6706\_C057
- RM6706\_C058

- RM6697\_C010
- RM6697\_C017
- RM6697\_C018
- RM6697\_C019
- RM6697\_C020
- RM6697\_C030
- RM6697\_C034
- RM6697\_C037
- RM6697\_C040
- RM6697\_C043
- RM6697\_C044
- RM6697\_C051
- RM6697\_C053
- RM6697\_C058
- RM6697\_C065
- RM6697\_C071
- RM6697\_C075
- RM6697\_C079

- RM6698\_C003
- RM6698\_C005
- RM6698\_C007
- RM6698\_C018
- RM6698\_C020
- RM6698\_C023
- RM6698\_C026
- RM6698\_C027
- RM6698\_C031
- RM6698\_C035
- RM6698\_C038
- RM6698\_C042
- RM6698\_C044
- RM6698\_C046
- RM6698\_C047
- RM6698\_C049
- RM6698\_C053
- RM6698\_C055
- RM6698\_C056
- RM6698\_C061
- RM6698\_C065
- RM6698\_C066
- RM6698\_C072
- RM6698\_C080
- RM6698\_C090
- RM6698\_C091
- RM6698\_C093
- RM6698\_C094
- RM6698\_C099
- RM6698\_C101
- RM6698\_C115

- RM6699\_C001
- RM6699\_C005
- RM6699\_C007
- RM6699\_C009
- RM6699\_C016
- RM6699\_C017
- RM6699\_C022
- RM6699\_C023
- RM6699\_C027
- RM6699\_C028
- RM6699\_C029
- RM6699\_C030
- RM6699\_C032
- RM6699\_C034
- RM6699\_C039
- RM6699\_C040
- RM6699\_C042
- RM6699\_C043
- RM6699\_C044
- RM6699\_C047
- RM6699\_C050
- RM6699\_C051
- RM6699\_C056
- RM6699\_C058
- RM6699\_C061
- RM6699\_C065
- RM6699\_C069
- RM6699\_C073
- RM6699\_C075
- RM6699\_C078
- RM6699\_C080
- RM6699\_C082
- RM6699\_C085
- RM6699\_C086
- RM6699\_C087
- RM6699\_C091
- RM6699\_C097
- RM6699\_C099
- RM6699\_C100
- RM6699\_C104
- RM6699\_C106
- RM6699\_C108
- RM6699\_C111
- RM6699\_C113
- RM6699\_C114
