## Supplementary figures and images for "Functional Convergence of Genetically Diverse B-Cell Receptors in Simian-HIV Infected Rhesus Macaques"

### Document S4

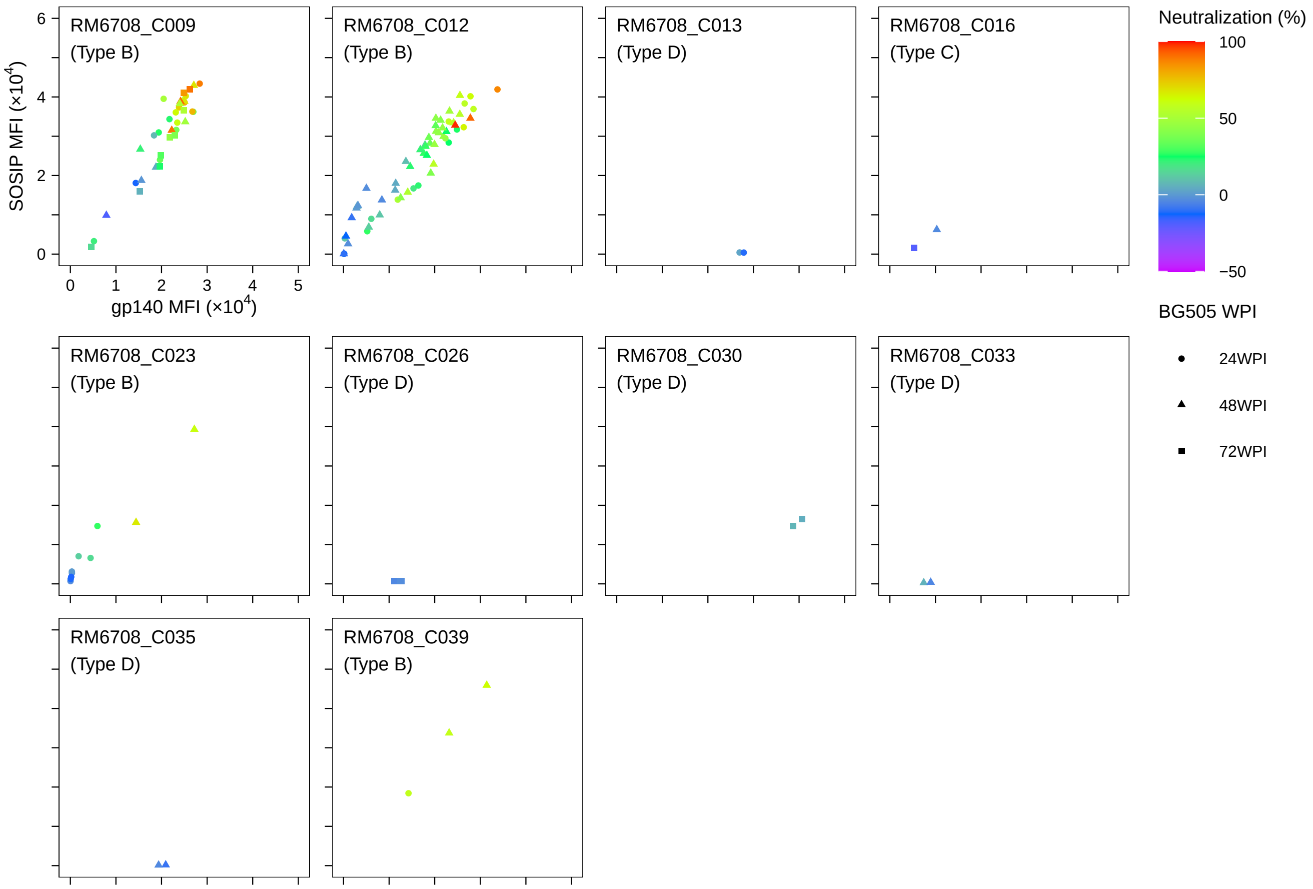

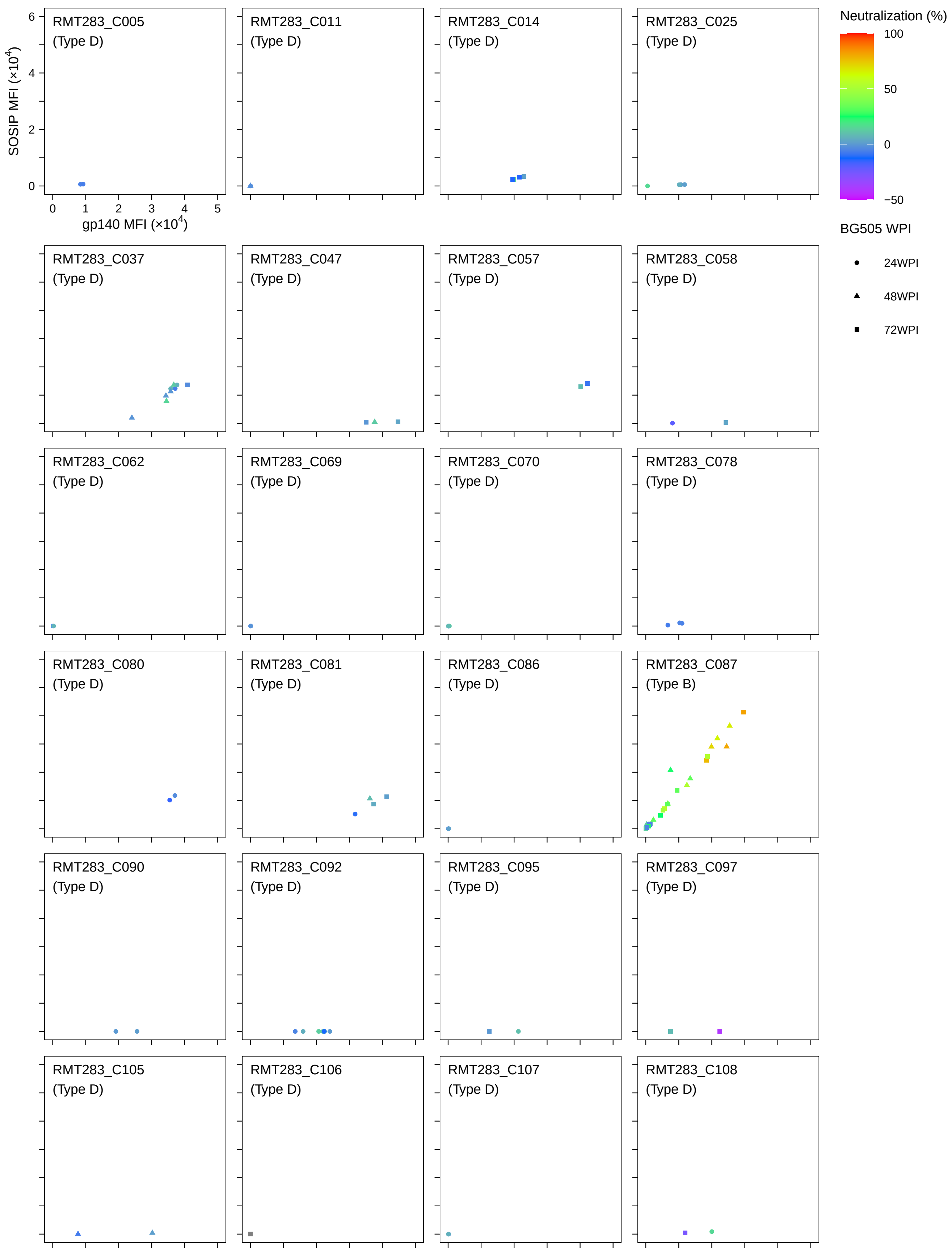

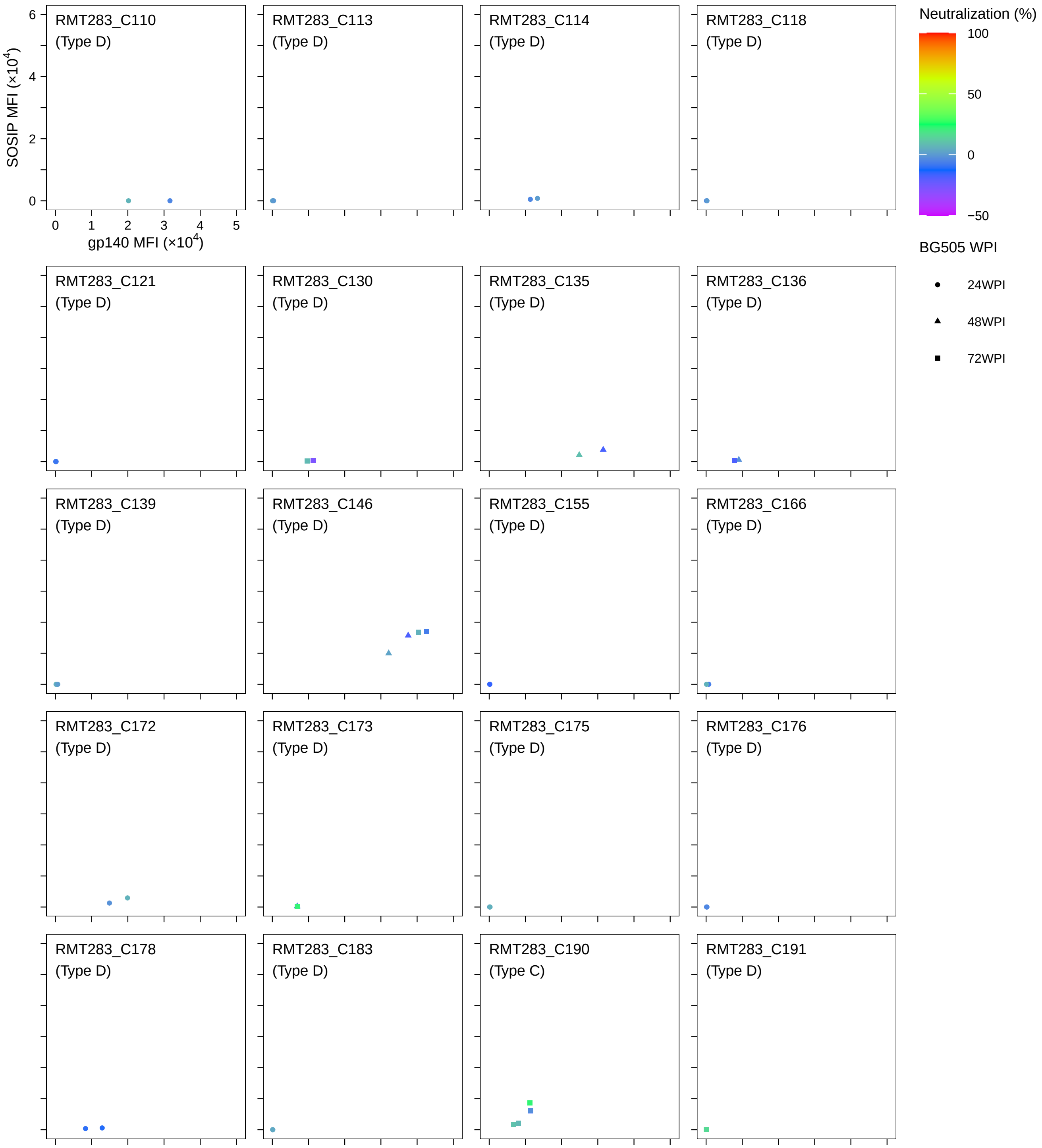

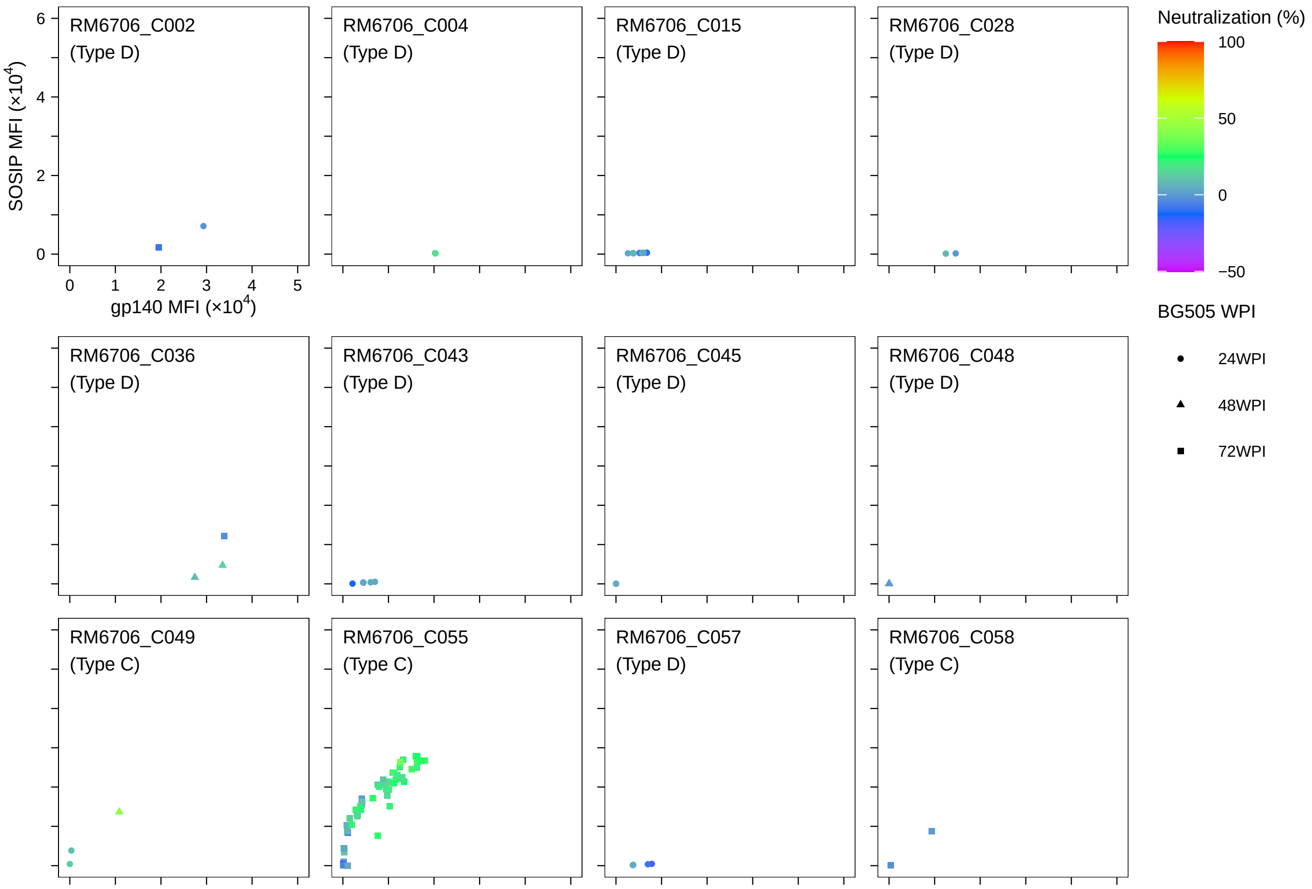

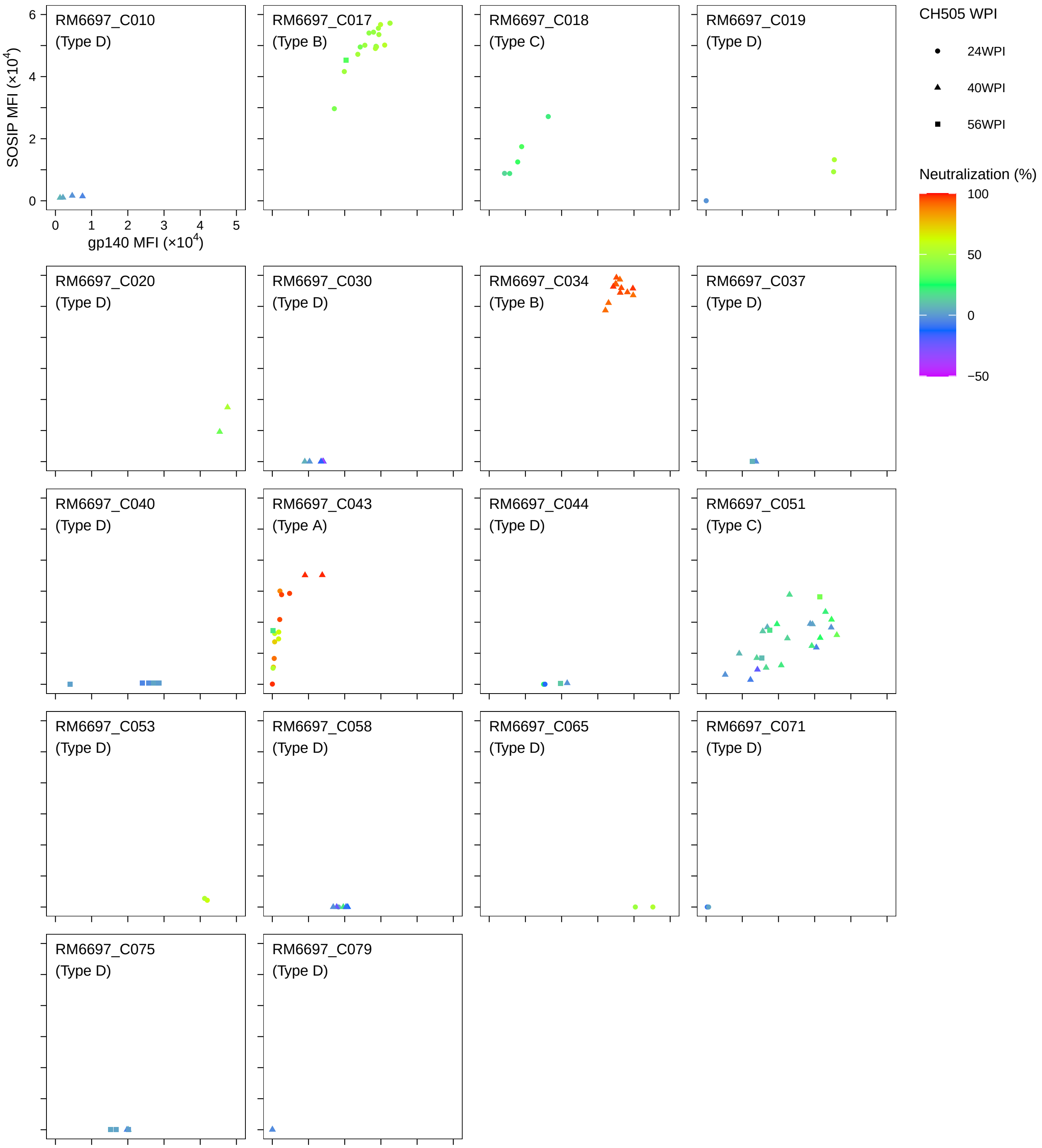

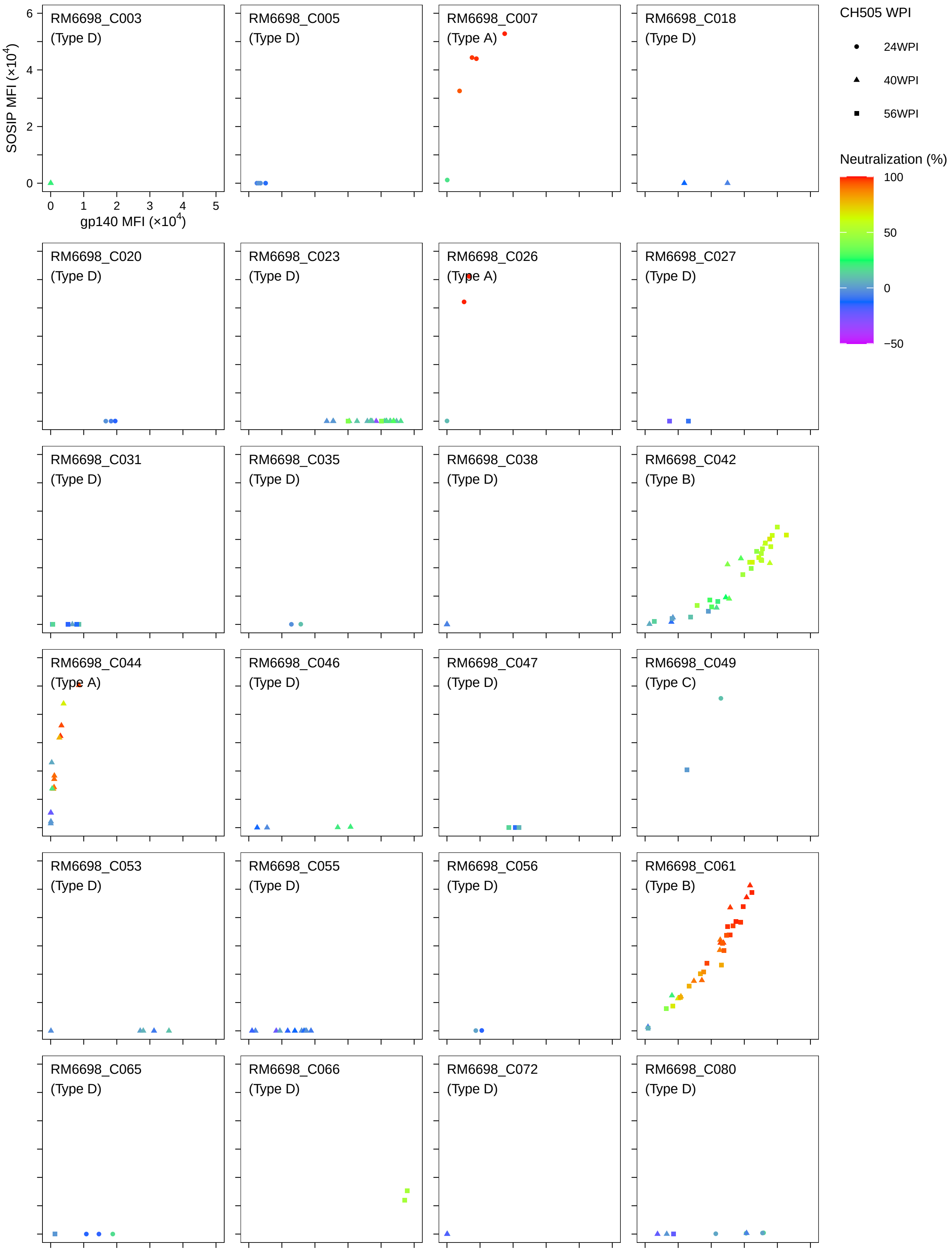

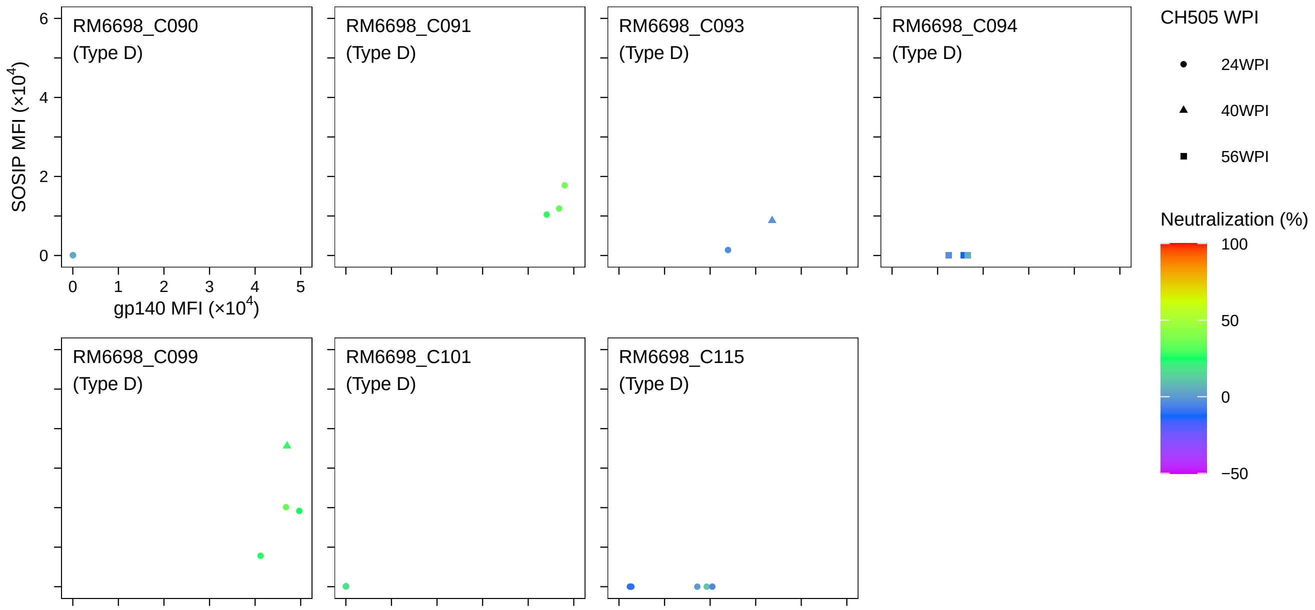

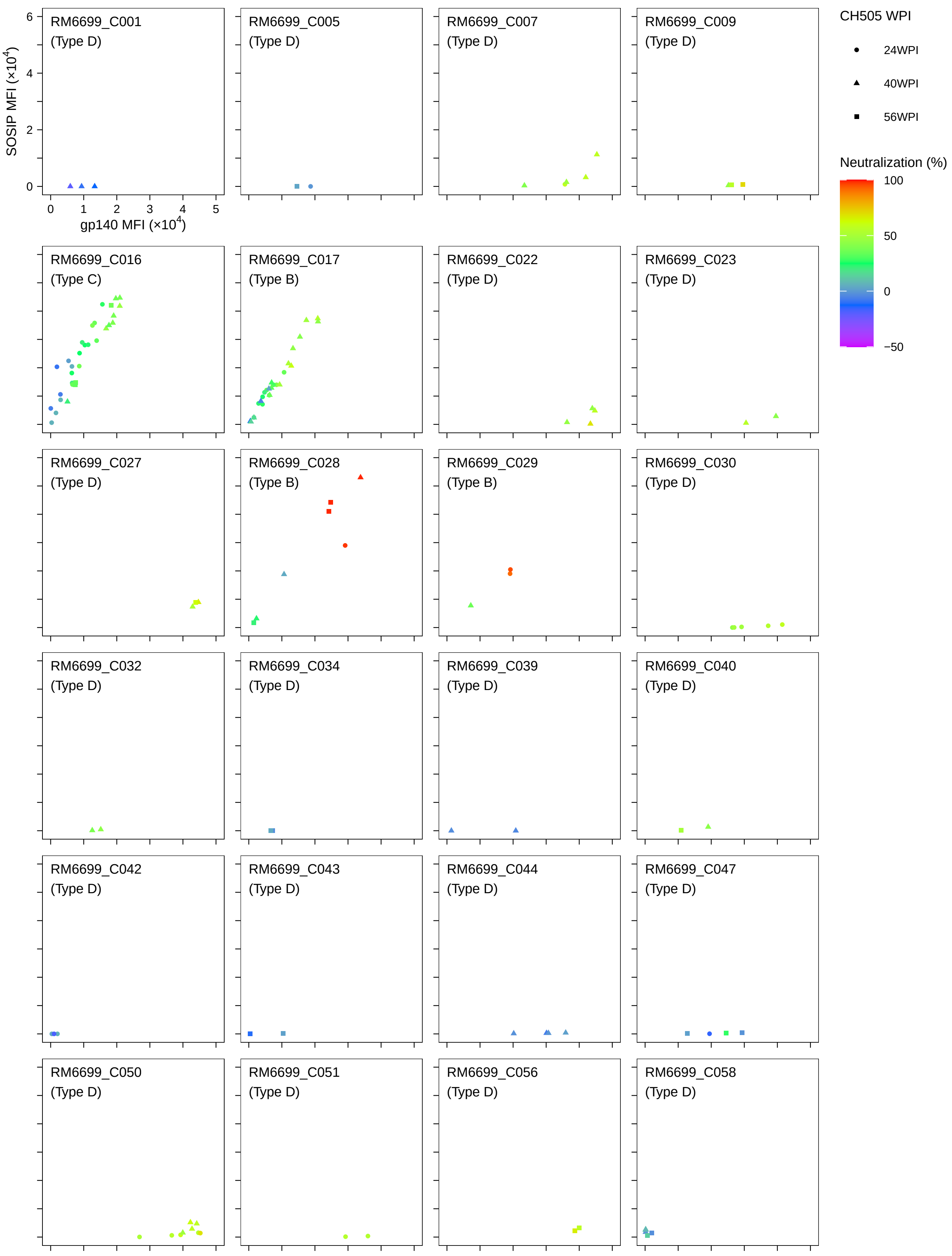

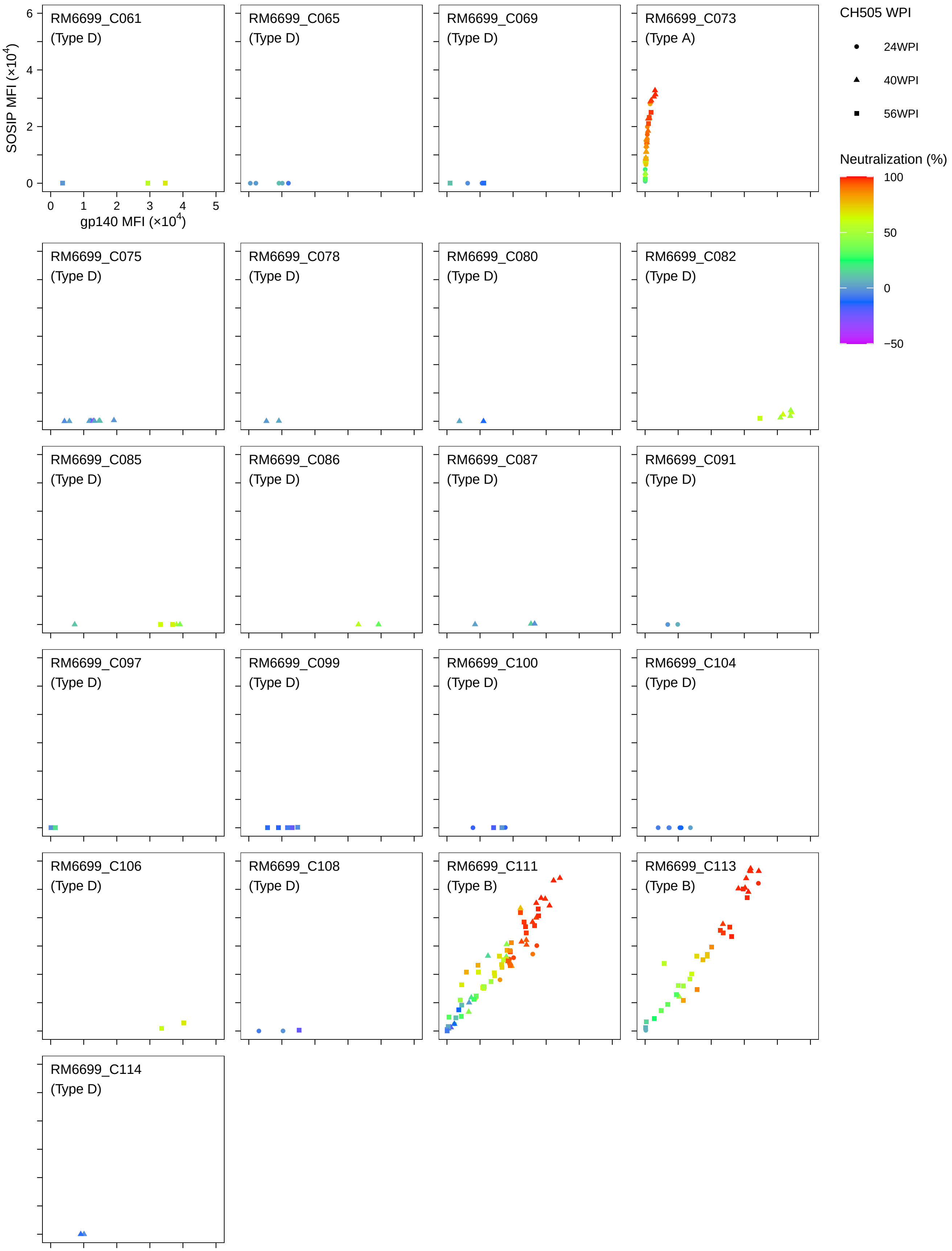

### Document S5

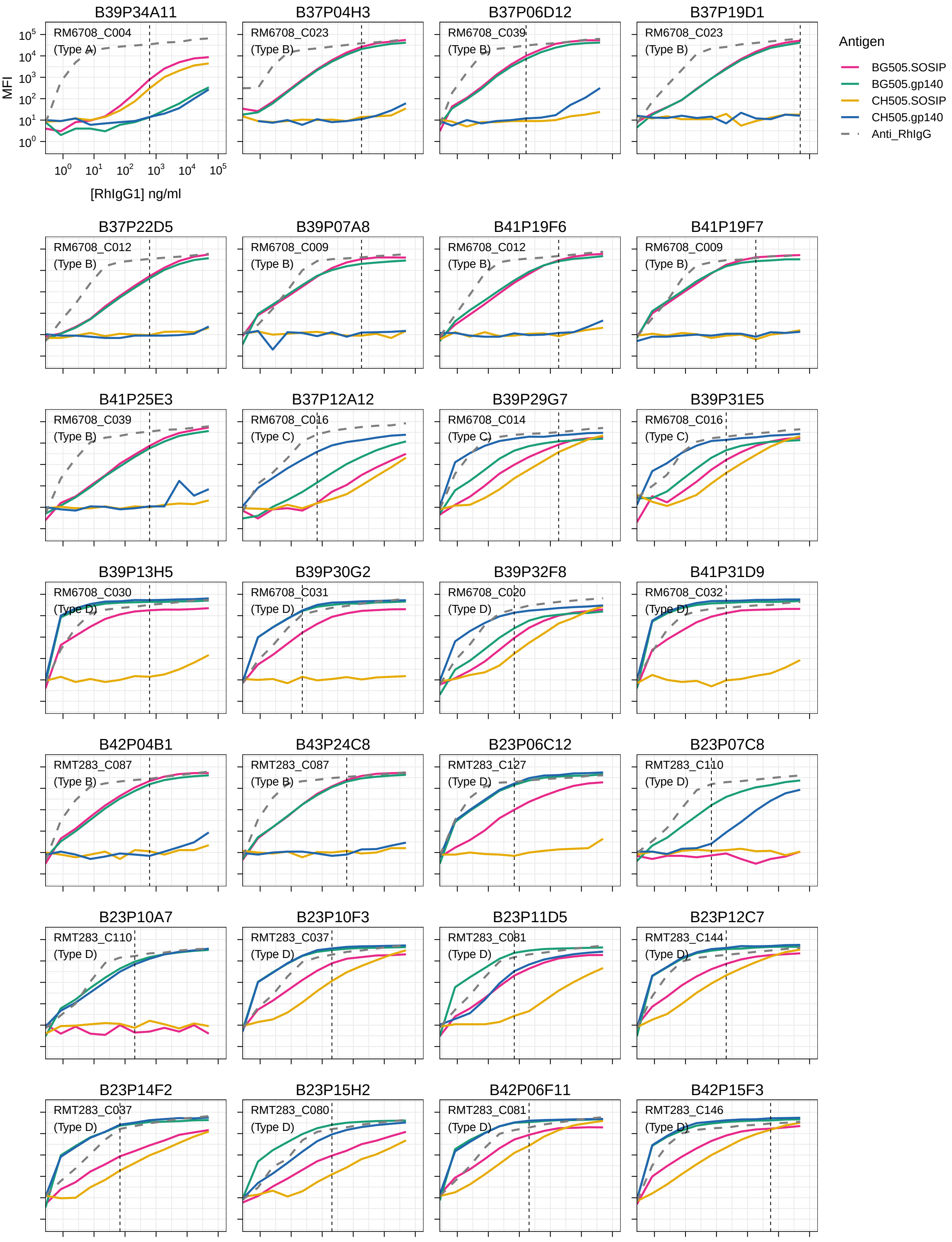

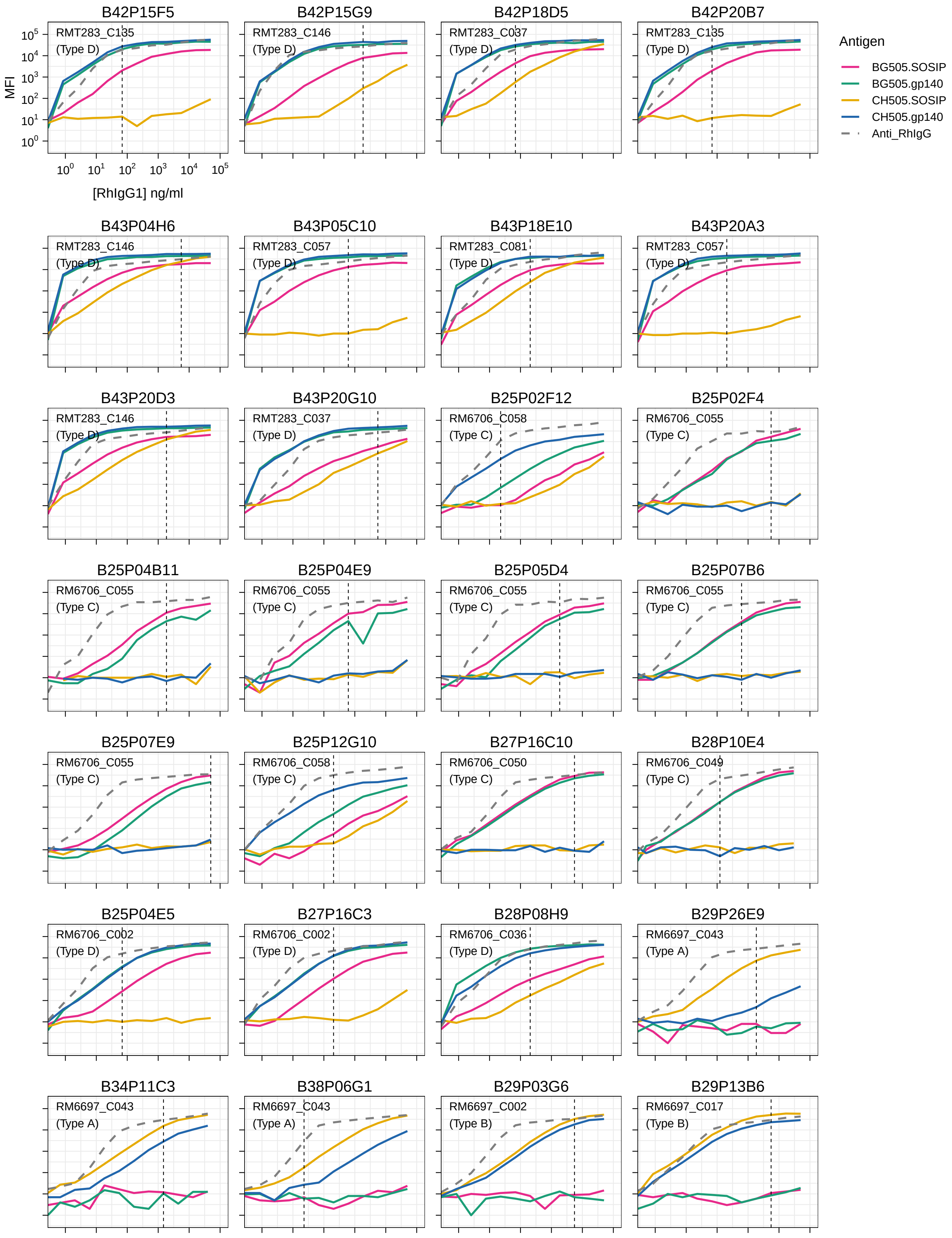

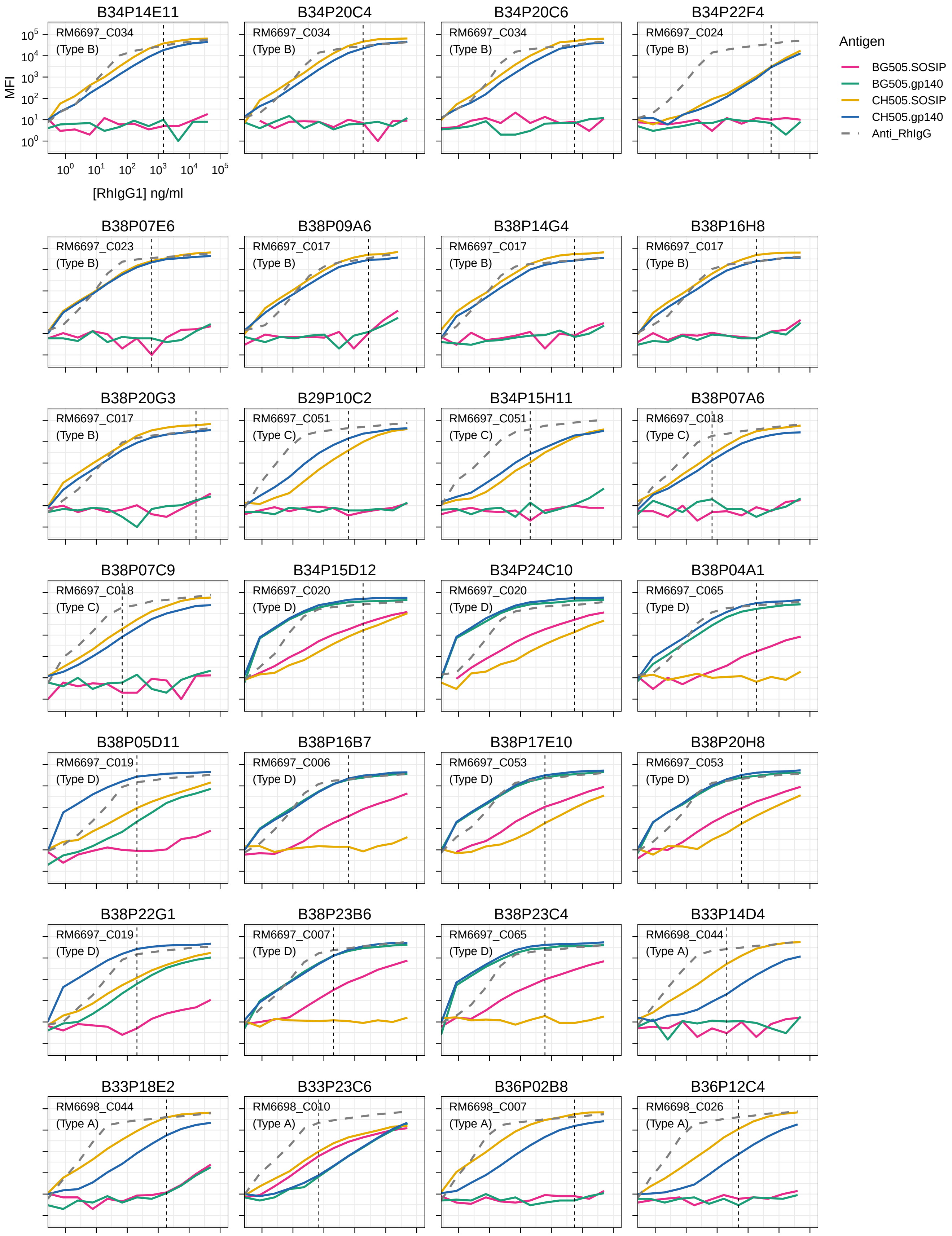

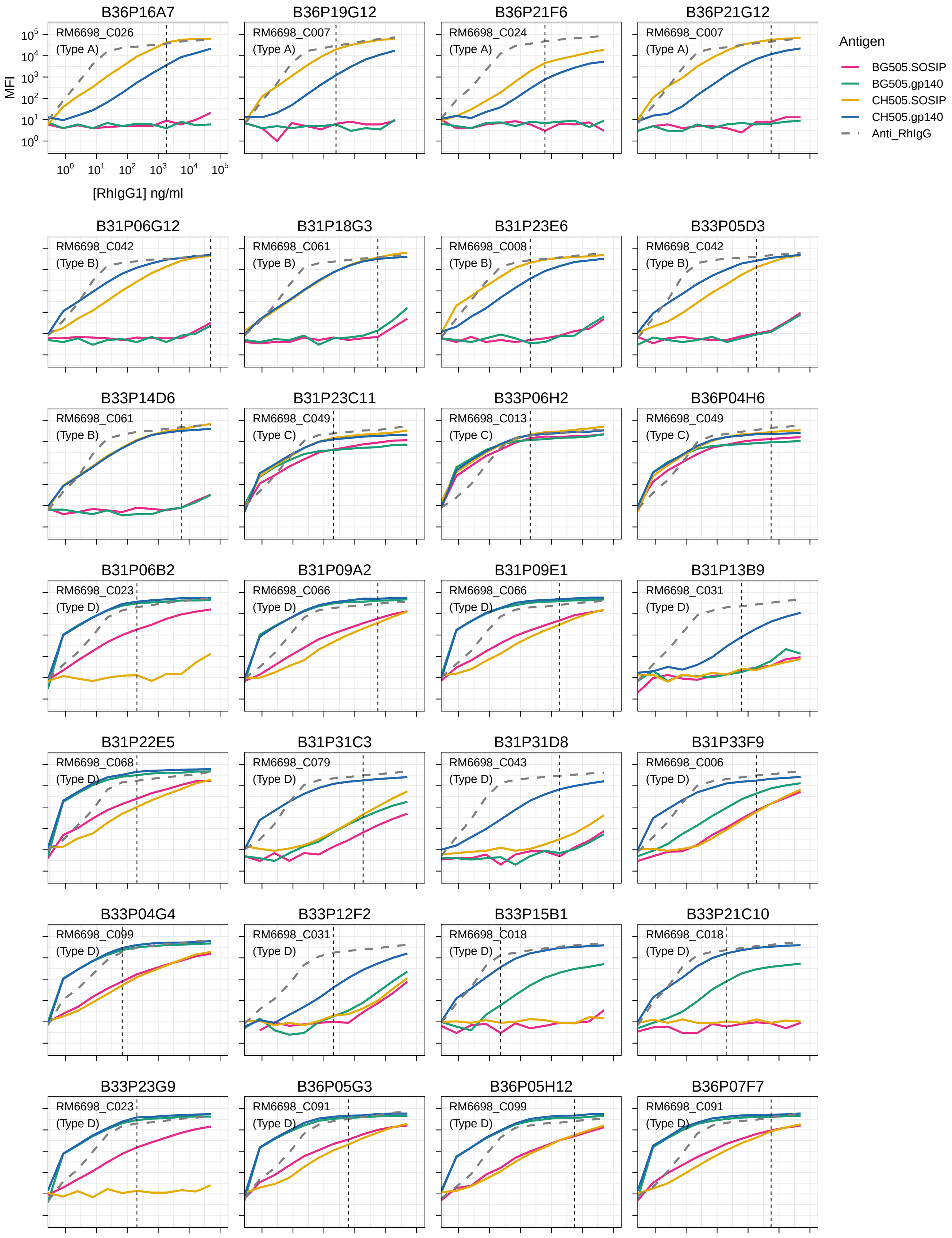

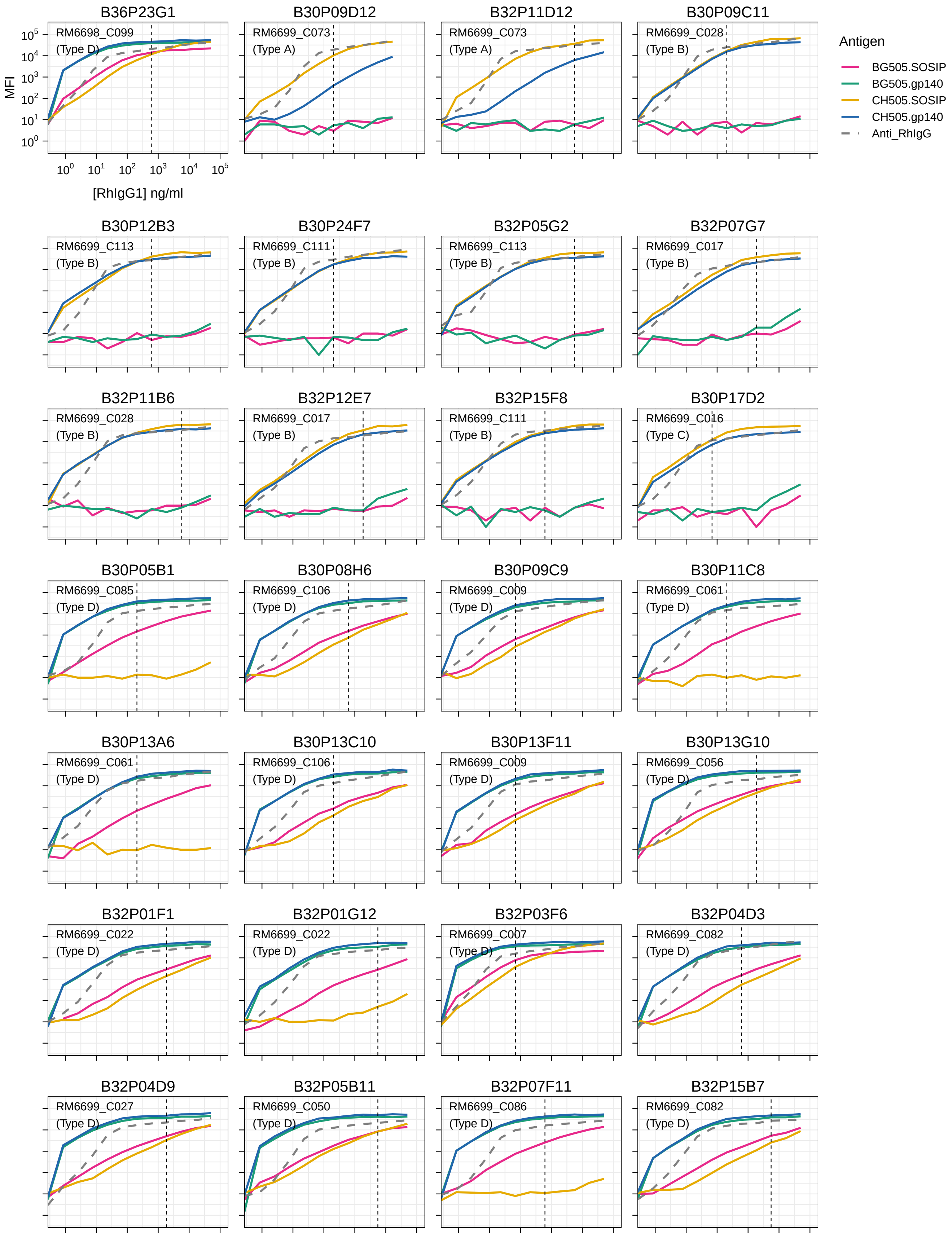

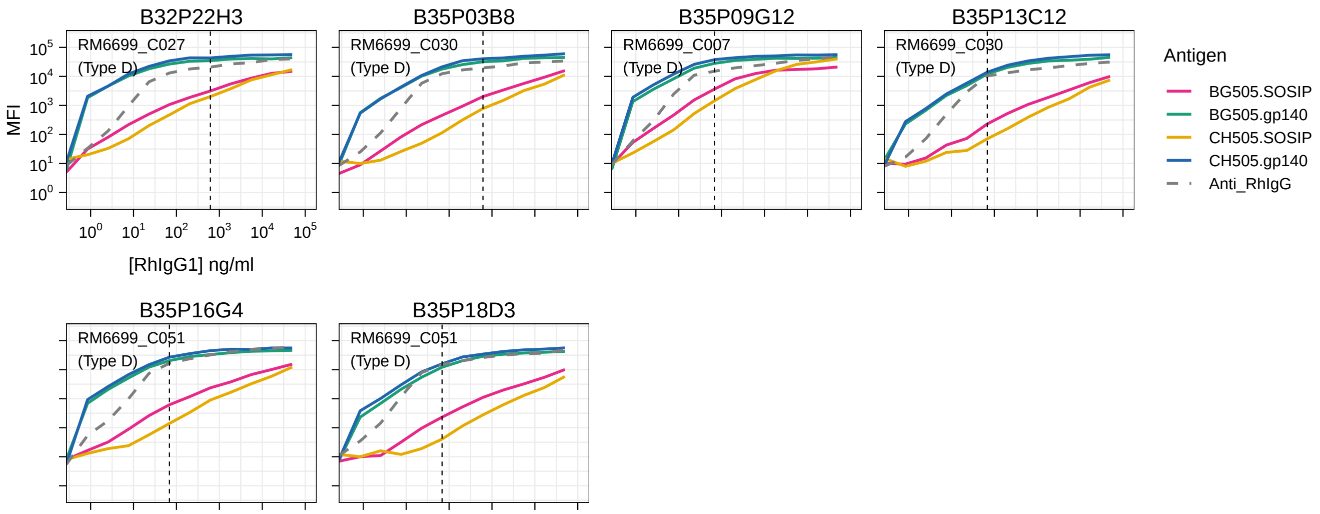
